# Probabilistic Graphical Modeling under Heterogeneity

**DOI:** 10.1101/2023.10.13.562136

**Authors:** Liying Chen, Satwik Acharyya, Chunyu Luo, Yang Ni, Veerabhadran Baladandayuthapani

## Abstract

Probabilistic graphical models are powerful and widely used tools to quantify, visualize and interpret dependencies in complex biological systems such as highthroughput genomics and proteomics. However, most existing graphical modeling methods assume homogeneity within and across samples which restricts their broad applicability to cases where sample-specific heterogeneity exists e.g. tumor heterogeneity. We propose a flexible Bayesian approach called <u>Graph</u>ical <u>R</u>egression (GraphR) which (a) allows direct incorporation of intrinsic factors of sample heterogeneity at different scales through a regression-based formulation, (b) enables sparse network estimation at a sample-specific level, (c) allows identification and uncertainty quantification of potential effects of heterogeneity on network structures, and (d) is computationally efficient through the use of variational Bayes algorithms. We illustrate the comparative efficiency of GraphR against existing methods in terms of graph structure recovery and computational cost across multiple realistic simulation settings. We use GraphR to analyze four diverse multi-omics and spatial transcriptomics datasets to study inter- and intra-sample molecular networks and delineate biological discoveries that otherwise cannot be revealed by existing approaches. We have developed a <u>GraphR R-package</u> along with an accompanying <u>Shiny App</u> that provides comprehensive analysis and dynamic visualization functions.

## Introduction

Network analyses are powerful tools for revealing and predicting underlying structures and dynamics in complex biological systems. Typically, complex biological systems consist of high-dimensional array of features including genes, proteins and other molecules that exhibit intricate interactions. The global structures and cumulative behavior of the systems cannot be inferred solely from partial factors of the system^[1]^. The increasing availability of high-throughput data and the comprehensive collation of biological data have indeed fueled the explosive interest in network-based approaches over the past few decades^[2,3]^. Specifically, biological networks capture the complex interactions between and within genomic molecular entities, enabling a holistic understanding of cellular functioning, organization and underlying etiology of disease; examples include protein-protein interaction^[4]^, gene co-expression networks^[5]^, and gene regulatory networks^[6]^. Such networks, synonymous with graphs, are compact representations of genomic interactions where features are nodes interconnected by edges (links) between them. These graphs provide a conceptual and intuitive framework to study interactions across multiple omics and spatial omics fields such as genomics, transcriptomics, and proteomics^[7]^.

Probabilistic graphical models (PGMs) are a common class of approaches to infer networks of complex systems that allow the measurement of uncertainty and provide probabilistic reasoning^[8]^. A PGM can be represented by a joint probability distribution over a high-dimensional space based on a graph which compactly encodes dependence structures among a set of nodes (variables), and the associated conditional distributions^[8]^. Furthermore, the associated probability distributions over networks formalize uncertain knowledge regarding the quantitative dependencies between variables while also enabling the incorporation of prior knowledge which further facilitates the decision making process^[9]^. Undirected graphical models are a widely-used subclass of PGMs, with edges in the graph representing conditional independence between two nodes. Specifically, if nodes follow a Gaussian distribution, one obtains Gaussian graphical models (GGMs) which extend conditional independence to partially uncorrelated structure which measures the association between two random variables after adjusting for the effect of all other variables^[10,11]^. Compared to pairwise-correlation networks, undirected graphical models tend to be sparser and more interpretable, capturing direct associations while explicitly removing the indirect associations^[12]^. GGMs are a representation of multivariate Gaussian distribution, with a sparse precision matrix where a zero entry is equivalent to conditional independence which admits attractive theoretical, interpretational, and computational properties^[13]^.

GGMs have undergone intense methodological and computational developments in recent years^[3,14,15]^. This progress has facilitated their widespread use across various biomedical domains, especially in genomics research^[16,17]^. Despite their widespread use, most of the GGM-based methods make a fundamental assumption of *homogeneity*, i.e. samples arise from a common population implying that the samples are independent and identically distributed^[14,18,19]^. This assumption may not be valid in many modern applications that generate heterogeneous data. One of the canonical examples that arise in cancer research is *tumor heterogeneity*^[20]^. Tumors usually present substantial interand intratumor heterogeneity, such as distinct genotypes and phenotypes of different regions within a single tumor^[21]^, or across same type (or subtype) of cancers^[22]^. Tumor heterogeneity plays a critical role in oncology research and clinical practice, providing valuable insights into disease progression, therapeutic selection and clinical outcomes of patients; for example, breast cancer subtype-based systemic therapy selection^[23]^, EGFR mutations and anti-EGFR therapies in lung cancer^[24]^, stem cell gene expression programs and clinical outcomes in human leukemia^[25]^ and intratumor heterogeneity levels and cancer evolution in human glioblastoma^[26]^. Tumor heterogeneity manifests itself at multiple intrinsic levels such as sub-population or sample-specific level. Hence, GGM-based methods that appropriately account for intrinsic heterogeneity in constructing genomic networks can lead to a better understanding of the underlying mechanism of tumors. Ignoring the potential impact of these intrinsic heterogeneity factors, such as cancer subtypes or dedifferentiation stages may lead to biased results which fail to reflect the true nature of associations between molecular markers, primarily due to Simpson’s paradox^[27]^. A simple illustration of Simpson’s paradox in graphical models is provided in the next section (also see Figure 2) where the assumption of homogeneity leads to spurious associations and appropriate accounting for intrinsic heterogeneity can reveal true biological signals. Hence graphical models that incorporate various types and scales of heterogeneity and allow for network inference at the sub-population or individual sample level, have the potential to uncover the true biological structures that can considerably aid precision medicine endeavors^[28]^.

In recognition of this fact, a few methods have been developed over the last few years for individualized networks estimation. These methods can be broadly classified into two categories: unsupervised (“top-down”) and supervised (“bottom-up”) approaches. Unsupervised approaches in the top-down direction typically involve de-convolution of population-level networks to infer sample-specific networks, implemented using unsupervised algorithms^[4,29,30]^. However, the top-down approaches do not allow assessments of direct impact of heterogeneity measures on network edges in a straightforward manner and thus fail to identify and quantify potential confounders due to lack of inferential procedures for individual networks. More recently, bottom-up approaches have been developed that alleviate this to some degree, as they allow a direct incorporation of heterogeneity measures and implemented by supervised algorithms via frequentist or Bayesian approaches^[31,32,33,34,35]^. In a frequentist paradigm,^[31,32]^ sample-specific graphs are inferred based on penalty functions, which are not geared to provide uncertainty quantification. Supervised Bayesian approaches allow the estimation and inference of individualized networks^[33,34,35]^ primarily through computationally-intensive sampling-based approaches, which stymies their application to high-dimensional genomic datasets. Furthermore most if not all methods, are not designed to incorporate multiple number and scales heterogeneous intrinsic factors and thus are not broadly applicable.

To address these challenges, we develop a unified, flexible and computationally scalable Bayesian framework for estimating network structures incorporating multiple types and scales of intrinsic heterogeneity, called Gaussian <u>Graph</u>ical <u>R</u>egression (GraphR). GraphR employs a regression-based formulation in the spirit of previous works^[33,35]^ to estimate graph structures in GGMs where precision matrices are linear functions of heterogeneity features which are represented by intrinsic factors (as covariates in regression setting). GraphR takes input in the form of a feature matrix (e.g. genes or proteins expression) and samples along with intrinsic factors in diverse combinations, allowing for incorporation of multiple types of heterogeneity (Figure 1). Some canonical examples of intrinsic factors and resulting graph structures include (I) binary intrinsic factors, such as disease and control group leading to binary group graphs; (II) categorical intrinsic factors, e.g. multi-category graphs for cancer subtypes; (III) a univariate continuous intrinsic factors, e.g. graphs varying over dedifferentiation levels; (IV) combinations of categorical and continuous intrinsic factors, such as graphs changing over cancer subtypes and continuous biomarkers; (V) multivariate continuous intrinsic factors such as application to spatial transcriptomics data resulting in spatially varying graphs. Additionally, GraphR can also identify and quantify the potential effects of intrinsic factors on graph structures, enhancing the understanding and interpretation of underlying mechanisms at a sample-specific level.

**Figure 1:**
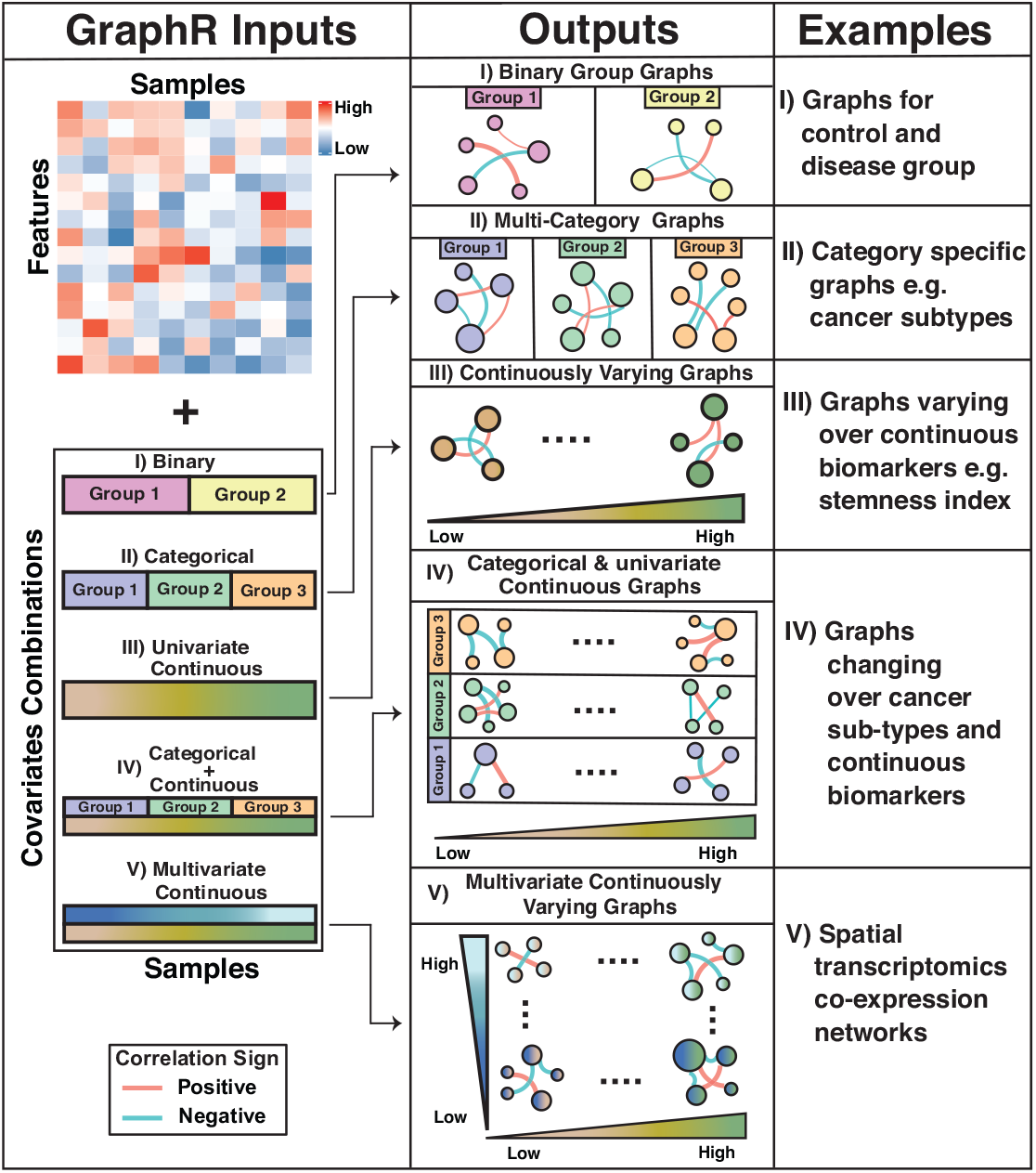
Overview of GraphR. The input for GraphR is a feature matrix (e.g. genes or proteins expression) and intrinsic factors reflecting heterogeneous structure (e.g. cancer types, dedifferentiation levels or spatial domains). The GraphR method can infer individualized graphs while accommodating different combinations of intrinsic factors. **(I)** binary intrinsic factors, such as control and disease group leading to binary group graphs; **(II)** categorical intrinsic factors, e.g. multi-category graphs for cancer subtypes; **(III)** a univariate continuous intrinsic factors, e.g. graphs varying over dedifferentiation levels; **(IV)** combinations of categorical and continuous intrinsic factors, such as graphs changing over cancer subtypes and continuous biomarkers; **(V)** multivariate continuous intrinsic factors such as application to spatial transcriptomics data resulting in graphs are changing over whole spatial domain.

We demonstrate the performance and versatility of GraphR through multiple simulation settings and multiple ‘omics data applications. In simulations, GraphR is compared with six state-of-the-art relevant methods^[14,31,36,19,37]^ in multiple scenarios consisting of multicategory, continuous varying and homogeneous graph structures. We show that GraphR precisely recovers the graph structure and accurately estimates graphical pattern changes with intrinsic factors in comparison with other methods. We further apply GraphR to four diverse breast cancer-related datasets that include including proteogenomics and spatial transcriptomics data, with intrinsic heterogeneity factors being breast cancer subtypes, dedifferentiation levels (stemness indices), pan-gynecological cancer types and spatial locations. Our key findings include: (a) identification of phosphorylated EGFR in Luminal A + B and Her2-enriched breast cancers with high co-expression with phosphorylated Her2 and Shc proteins; (b) Additionally, phosphorylated EGFR is also detected as a hub protein in endometrial carcinoma but not in cervical and ovarian cancers; (c) In terms of stemness indices, partial correlation between GAPDH and FOXM1 tends to be higher with increasing dedifferentiation levels; (d) Finally, applying GraphR to spatial transcriptomics breast cancer data reveals co-activation (positive correlation) of XBP1 and SPINT2 in the tumor region and conversely, co-repression (negative correlation) in the normal region, highlighting the method’s profitability. Furthermore, we have developed an <u>R-package</u> and an accompanying <u>Shiny App</u> that is easily accessible to the scientific and research community – to obtain, inspect, rerun, and more importantly apply the methods on their own datasets, without coding experience and implementations.

## Results

Our results are organized as follows. First, we motivate the need for GraphR through an illustration of Simpson’s paradox in graphical models. Subsequently we describe the construction of GraphR to incorporate intrinsic factor-based heterogeneity. Next, we utilize synthetic datasets to evaluate the operating characteristics of the GraphR method and compare it with other graphical modeling approaches across multiple scenarios. Finally, we illustrate the versatility of GraphR using four data sets in breast cancer: proteomic network-based characterization of breast cancer subtypes, continuously varying proteomic network over the scale of stemness indices for breast cancer, pan-cancer proteomic networks across gynecological cancers, and spatially varying gene regulatory networks for spatial transcriptomic data from a breast cancer tissue.

### Illustration of Simpson’s Paradox in Graphical Models

We motivate the development of GraphR using an illustration of Simpson’s paradox in graphical models. Simpson’s paradox broadly refers to a scenario where trends between two variables reverse or disappear after adjusting for some other variables. To illustrate the key principle, we present a simple example in a graphical model setting with three nodes. Assume we draw samples from two heterogeneous groups with distinct conditional independence structures (Figure 2 Left panel) where the partial correlation between node A and B (conditioned on node C) is negative (partial correlation = -0.61; shown in blue) in the first group and positive (partial correlation = 0.56; shown in red) in the second group, respectively (the details of construction design is discussed in Section B.3 of the Supplementary Information). However, if we treat all samples as homogeneous (i.e. coming from a single group), then the partial correlation between the two nodes is obfuscated (partial correlation = 0.20; Figure 2 Right panel). This demonstrates that even for a simple three-node graph, ignoring discrete heterogeneity leads to erroneous conclusions. In more complex scenarios, such as existence of multiple confounders at different scales (categorical and continuous), the inference of graphical structures among multiple genomic features is more challenging and motivates development of graphical models that incorporate this additional axis of information to construct heterogeneous networks.

**Figure 2:**
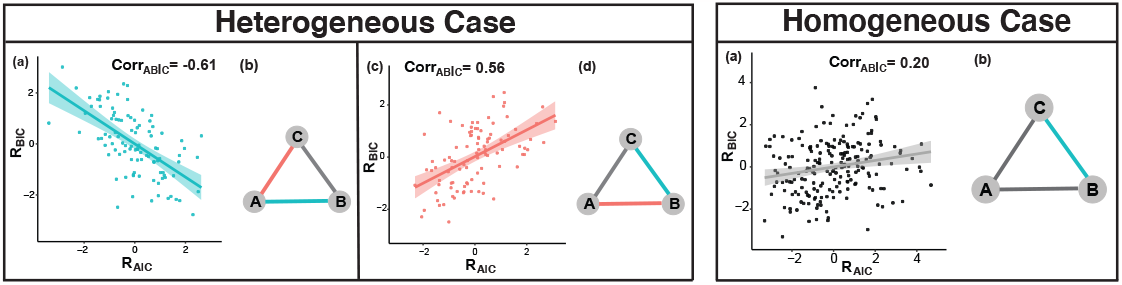
Illustration of Simpson’s Paradox in Graphical Models. Samples are drawn from two heterogeneous groups with distinct conditional independence structures. **Heterogeneous Case** considers heterogeneous group effects. Figure (a) and (c) of the left panel denotes point plots of residuals from linear regression of node A with node C and node B with node C respectively, with linear regression as the smoothing function (green and red line respectively). The observed partial correlation is -0.61 and 0.55 in the two groups. Figure (b) and (d) of the left panel shows estimated conditional independence structures from GraphR for two groups. Colors of edges represent signs of partial correlation, with green and red being negative and positive respectively, and grey indicating no connection. **Homogeneous Case** shows results when treating all samples homogeneous. Figure (a) in the right panel represents association between residuals from linear regression of node A with node C and node B with node C. The observed partial correlation is 0.20. All other settings are same as Heterogeneous Case. Figure (b) in the right panel characterizes the estimated conditional independence structures from GraphR and rest of the settings are same as Heterogeneous Case.

### GraphR construction

GraphR model construction is described in *<u>Methods</u>* <u>Section</u> with its technical details provided in Section A of *Supplementary Information* and the schematic described in Figure 1. Briefly, let **Y** denote a *p−*dimensional vector of genomic features (e.g. proteins/genes) and **X** denote a *q −* dimensional vector of intrinsic factors driving heterogeneity (e.g. cancer subtypes, spatial locations) which are assessed on the same *n* samples. Standard GGMs for **Y** can be represented as **Y***∼ N* (0, Ω^*−*1^), where *N* (*•*) is the Gaussian distribution. Elements in the precision matrix (Ω^*−*1^) encode the partial correlations, with zero elements denoting conditional independence. In GraphR we generalize this construction to **Y**| **X***∼ N* [0, Ω(**X**)^*−*1^], where-in the partial correlations now are explicit functions of the intrinsic factors **X** which modulate the dependence patterns of the feature vector **Y**. We develop a regression-based procedure for estimation that allows: a) sparse estimation via spike-and-slab priors; (b) computational efficiency through the use of scalable variational Bayes algorithms; (c) uncertainty quantification through posterior probabilities; and (d) downstream mining of the heterogeneous networks to identify hub genes, pathways patterns, degree and connectivity for subsequent biological interpretations.

### Synthetic Datasets

To demonstrate the operating characteristics of GraphR, we provide simulation results under two broad scenarios: undirected graphs with unordered pairs of nodes and directed graphs where directions of edges are apriori known. For the undirected setting, we consider the following three cases (I) pre-specified categories (discrete heterogeneity), (II) with intrinsic factors being continuous variables and (III) a homogeneous population. In terms of directed graphs scenarios, we show the performance of GraphR for varying dimensions of features and intrinsic factors. GraphR estimation involves *𝒪* (*p*^2^*q*) parameters whereas a standard homogeneous GGM estimates a graph of order *𝒪* (*p*^2^). This problem becomes considerably more challenging under high-dimensional scenarios with limited sample size. Our core hypothesis is that, by accounting for different scales of intrinsic factor-based heterogeneity, enables better network structural recovery for both undirected and directed graphs.

### Overview of simulation design

For undirected setting, we fix *n* = 151; *p* = 33 and *q* = 2, and compare our approach to six relevant methods: (I) Bayesian Gaussian graphical models (BGGM^[19]^) (II) graphical lasso (GLASSO^[14]^) (III) fused graphical lasso (FGL^[36]^) (IV) group graphical lasso (GGL^[36]^) (V) Laplacian shrinkage for inverse covariance matrices from heterogeneous populations (LASICH^[37]^) (VI) kernel graphical lasso (K-GLASSO^[31]^). A concise overview of the competing methods has been provided in Section A.6 of the Supplementary Information. We further illustrate the operating characteristics of GraphR with varying dimensions of features (50 to 500), intrinsic factors (2 to 10) and sample sizes (100 to 5000) through datasets generated from directed acyclic graphs (DAGs). To evaluate the performance of network recovery in both settings, we give comparative metrics: (1) area under the ROC curve (AUC), (2) true positive rate (TPR), (3) false positive rate (FPR), (4) Matthew’s correlation coefficient (MCC, that jointly considers both TPR and FPR) and (5) false discovery rate (FDR). Further details about simulation design are provided in the <u>Methods Section</u> and Section B of the Supplementary Information.

### Undirected graphs

Under multi-category graph setting in Figure 3A, GraphR outperforms all the competitive methods in terms of MCC (<u>GraphR: 0.892</u>; FGL: 0.559; GGL: 0.56; LASICH: 0.611) and a marginally similar AUC (<u>GraphR: 0.953</u>; FGL: 0.97; GGL: 0.971; LASICH: 0.974). In comparison with GraphR, the competitive methods have high values of TPR at the expense of higher FDR and FPR. This pattern is well captured by MCC since it is an overall assessment considering other discovery rates. Notably, GraphR performs the best in case of individual specific precision matrices (i.e. continuous intrinsic factor-based heterogeneity). Figure 3B shows that GraphR achieves the highest MCC (<u>GraphR: 0.955</u>, BGGM: 0.354, GLASSO: 0.316 and k-GLASSO: 0.333) and AUC (<u>GraphR: 0.998</u>, BGGM: 0.838, GLASSO: 0.748 and k-GLASSO: 0.648) among other methods. In the end, we assess the performance of GraphR for homogeneous graphs in Figure 3C attains the highest MCC (GraphR: 0.955; BGGM: 0.951; GLASSO: 0.644) and marginally lower AUC (GraphR: 0.985; BGGM: 1; GLASSO: 0.984). The homogeneous case serves as positive control with respect to other simulation settings. Detailed values of other discovery rates and results from different simulation settings are provided in the <u>Methods Section</u> and Section B.1 of the Supplementary Information.

**Figure 3:**
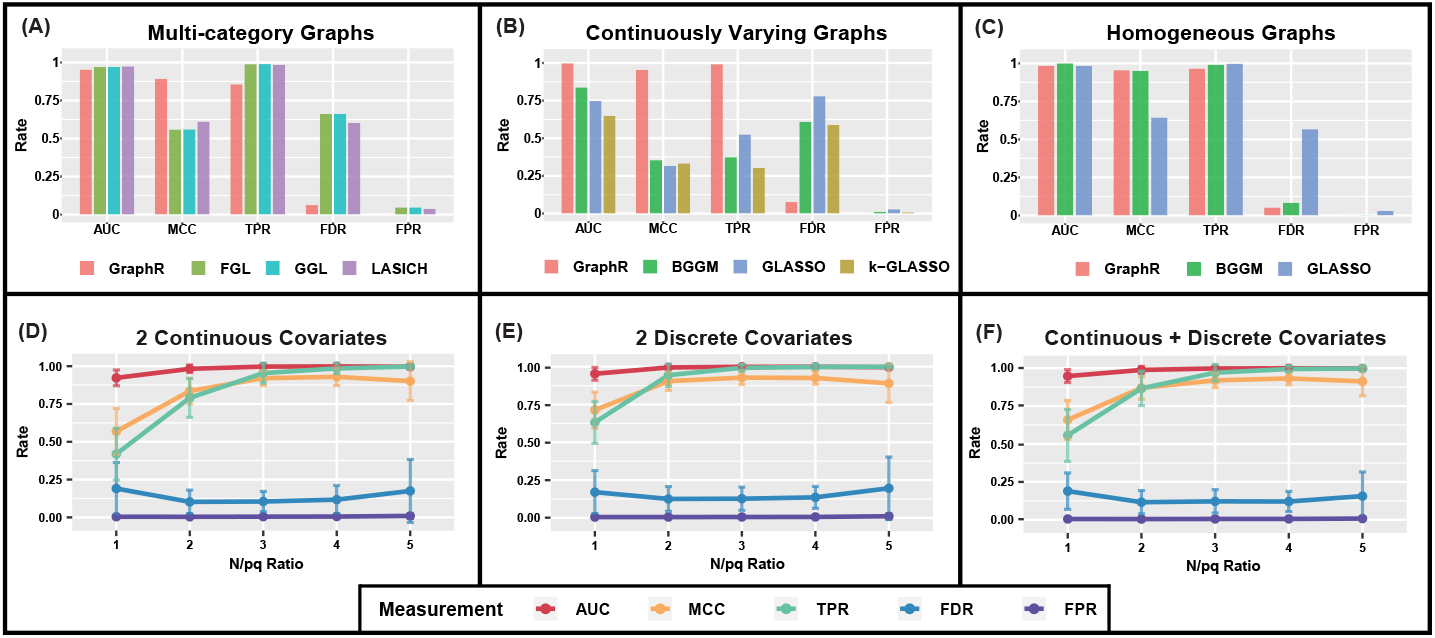
Simulation results. For each setting, we report the mean values of area under the ROC curve (AUC), Matthews correlation coefficient (MCC), true positive rate (TPR), false positive rate (FPR) and false discovery rate (FDR) based on 50 repetitions to evaluate the selection performance. **First row of the Figure 3 represents comparison of selection (recovery) performance among GraphR and several other methods under the undirected graphical settings**. We consider three cases here, where subjects are drawn from **(<u>3</u>A)** two different categories or **(<u>3</u>B)** each subjects are heterogeneous or **(<u>3</u>C)** a homogeneous population. Six relevant methods are used for comparison, involving BGGM, GLASSO, FGL, GGL, LASICH and K-GLASSO. **Second row of the Figure 3 shows selection performance for directed acyclic graphs** with number of nodes p = 50 and number of intrinsic factors q = 2. The x-axis is defined as the ratio between sample size n and p*q, ranging from 1 to 5. Effect size *β* is set to be 3 while the types of two intrinsic factors being continuous only **(<u>3</u>D)** or discrete only **(<u>3</u>E)** or continuous and discrete covariates **(<u>3</u>F)**

### DAGs

We further demonstrate the operating characteristics of GraphR in directed graph scenarios with 50 nodes (*p* = 50), 2 intrinsic factors (*q* = 2) and varying sample sizes (*n* = 100, 200, 300, 400, 500). GraphR generally performs better with increasing ratio between *n* and *pq* and effect size *β* (second row of Figure 3, Supplementary Figure B6 and B7). However, FDR increases when *n* over *pq* ratio reaches 5 (continuous: 0.174; discrete: 0.196; continuous and discrete: 0.156) compared to the case when *n* over *pq* ratio 4 (continuous: 0.117; discrete: 0.136; continuous and discrete: 0.120), leading to moderate decrease in MCC from (0.929, 0.926, 0.932) to (0.901, 0.89, 0.913) with settings in Figure 3D,3E, 3F respectively.

<u>In summary</u>, the GraphR method is robust and performs better in terms of structural recovery in presence of intrinsic factor-based heterogeneity along with well controlled FDR across multiple scenarios such as undirected and directed graphs. Under heterogeneous structure, GraphR method achieves the highest power than other methods with the lowest FDR and FPR, indicating the importance to incorporate heterogeneity.

### Illustrative Examples

We apply GraphR to four diverse datasets in breast cancer (BRCA) to characterize different aspects of heterogeneity of proteogenomic networks. BRCA is the most common cancer among women in the United States with around 3 million newly diagnosed cases and 43K deaths in 2022^[38]^. Accumulating evidence suggests that multiple factors affect the choice of treatment for breast cancer, such as subtypes and stages of cancer, as well as individual patient considerations^[23,39]^. To better dissect breast cancer heterogeneity, we utilize GraphR to explore the proteomic characterization of intrinsic subtypes of BRCA, investigate effects of dedifferentiation levels (stemness indices), assess differences and commonalities in proteomic networks across pan-gynecological cancers and effect of the tumor microenvironment using BRCA spatial transcriptomic data.

### Proteomic Network-based Characterization of Intrinsic Subtypes of Breast Cancer

We focus on characterization of the shared and subtype-specific proteomic networks of intrinsic subtypes of BRCA^[40]^. These subtypes have demonstrated distinct genomic features and have been established as crucial prognostic factors in treatment selection and clinical outcomes^[41,42]^; however, their proteomic networks have been under-explored. To this end, we apply GraphR to 190 proteins obtained from 626 BRCA patients from The Cancer Genome Atlas (TCGA^[42]^) across three subtypes (as discrete factors): Luminal A and B (n=626), Her2-enriched (n=75) and Basal-like (n=158). Proteomics data are obtained and normalized from The Cancer Proteome Atlas^[43]^ through Reverse Phase Protein Arrays (RPPAs) to quantify proteins or phosphoproteins which are involved in multiple functional and signaling pathways such as apoptosis, DNA damage response, and cell cycle^[44]^.

We aim to estimate the cancer subtype-specific networks and only include protein pairs belonging to different isoforms while showing significantly strong partial correlation (FDR-based p-value *<* 0.01 and| partial correlation|*≥*0.4) for at least one group. Proteins with top 5 connectivity degrees are used to plot cancer subtype-specific networks in Figure 4A, and networks involving all included proteins are shown in Supplementary Figure C2-C4. We find several proteins are conserved across groups such as phosphorylated EGFR i.e. EGFR* ^1^ for Luminal A + B and Her2-enriched with relatively high co-expression with Her2*, Shc and Src*. This is consistent with previous findings that high correlation of phosphorylated EGFR and Her2 is observed in Her2-containing RPPA-defined subgroup which includes most of Her2-enriched and part of luminal A + B BRCA defined by PAM50^[45]^. Furthermore, Shc has been proven as a key factor involved in EGFR downstream signaling pathway and the interaction between Shc and EGFR is related to the activation of MAPK pathway^[46,47]^, while the synergism between Src and EGFR is associated with the acceleration of oncogenesis process, resulting in the increased level of Shc^[48,49]^.

**Figure 4:**
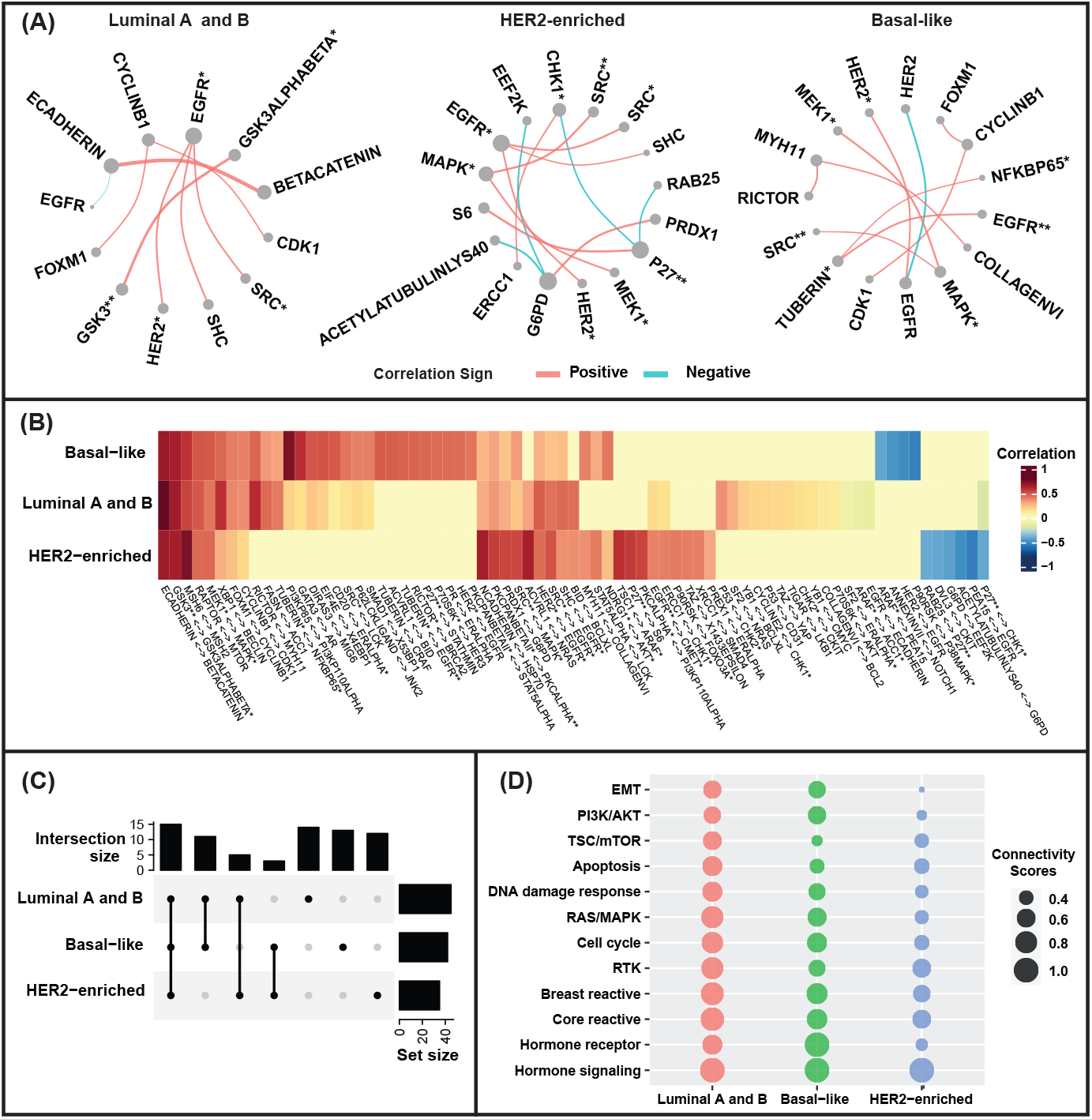
Proteomic Network-based Characterization of Intrinsic Subtypes of Breast Cancer. **(A)** PAM50 subtype-specific networks for proteins with top five connectivity degrees. **(B)** Heatmap of partial correlations between protein pairs for each subtype of breast cancer. We include protein pairs with significantly strong partial correlations (FDR based p-values *<* 0.01 and| partial correlation |*≥* 0.4) for at least one subtype. **(C)** Upset plot for sizes of significant connections within or across BRCA subtypes based on the protein pairs included in (B). **(D)** Connectivity scores of pathways for each breast cancer subtypes. Pathways are ordered based on the clustering of connectivity scores.

Figure 4B shows the heatmap of partial correlations between selective protein pairs of three groups and the corresponding heatmap of posterior inclusion probabilities (PIP) is provided in Figure C1 of the Supplementary Information. The upset plot in Figure 4C demonstrates the numbers of significant connections within or across BRCA groups. Overall, 15 connections are identified as significant across all three groups while 13, 12, 14 connections only showed significance in Basal-like, Her2-enriched, Luminal A and B respectively, indicating the similarities and differences for each group of BRCA. In the heatmap, we can observe high partial correlations of proteins belonging to the same family, e.g. MSH2 and MSH6. Furthermore, we also find that E-cadherin and *β*-catenin are positively correlated in all groups of BRCA with partial correlation being 0.69, 0.69 and 0.98 in Basal-like, Her2-enriched and Luminal A and B BRCA. We observe negative partial correlations between p27* Chk1* in Her2-enriched and Luminal A and B BRCA with moderate and weak magnitude respectively (Her2-enriched: *ρ* = -0.42; Luminal A and B: *ρ* = -0.19). Interestingly, we do not identify this negative partial correlation in Basal-like BRCA.

We further combine information of 12 established proteomics pathways^[44]^ to transform connections between protein pairs into an integrated functional analysis where we group functional proteins belonging to the same pathway into GraphR algorithm. In order to gain insight into pathway-level connectivities within or across each subtype, we use connectivity score (CS) which is defined as the ratio of the connected edges to the total number of possible edges. Higher CS corresponds to denser connections in the pathway. The dot sizes of the bubble plot in Figure 4D are proportional to the CS, and clustering on CS is conducted to decide the order of pathways in Figure 4D. Generally, CS for Luminal A and B tend to be the largest among the three groups and smallest for Her2-enriched BRCA, except for PI3K/AKT pathway, TSC/mTOR pathway, hormone receptor pathway and hormone signaling pathway. With respect to hormone receptor pathway, CS for Basal-like BRCA is highest (CS = 1) and is equal to 0.67 and 0.33 in Luminal A and B and Her2-enriched cancer, partly due to the reason that all hormone receptors are likely to have a downexpression in Basal-like BRCA while the associations among hormone receptors are more complicated and diverse in the other two groups. Compared to Basal-like BRCA, Luminal A and B and Her2-enriched cancer had a slightly higher CS in terms of TSC/mTOR pathway (Basal-like: 0.3; Luminal A and B: 0.6; Her2-enriched: 0.4).

### Characterization of Stemness-induced Heterogeneity on Proteomics Networks in Breast Cancer

Our aim is to investigate changes in proteomic network patterns across various dedifferentiation levels of BRCA. Tumor cells exhibit dedifferentiation, a reversed growing process by which cells lose the specialized phenotype and gain properties of stem cells^[50]^. It is commonly accepted that the degrees of differentiation of tumor cells are significantly related to tumor behavior and thus have been widely used in the grading system for BRCA which provides guidance for treatment and outcomes^[51]^. To this end, two independent stemness indices are obtained based on DNA methylation (mDNAsi) and mRNA expression (mRNAsi) which provide the degrees of dedifferentiation on epigenetic and gene expression levels respectively, across BRCA tumors^[50]^. The mDNAsi and mRNAsi values range from 0 to 1 with lower values implying a tendency towards normal-like cells. We apply GraphR using the same proteomics dataset as in the previous example along with two stemness indices across 616 BRCA patients’ data from TCGA^[42]^, treating mRNAsi, mDNAsi and patients’ ages (as a potential confounder) as three continuous intrinsic factors. Details of the preprocessing procedure can be found in Section D.2 of the Supplementary Information.

We estimate proteomic networks across varying levels of mRNAsi, mDNAsi and age. Figure 5A shows the networks for proteins with high degrees of connectivity across different quartiles of mRNAsi while setting mDNAsi and age at their median (analogous plots for mDNAsi are shown in supplementary Figure D.3.). The network connections are included based on the following three criteria: (1) protein pairs with different isoforms; (2) significant partial correlation (FDR based p-values *<* 0.01) in more than half of the cases. (3) | partial correlation |*≥* 0.45 for at least one case. The median value of partial correlation between significant protein pairs are obtained for each quartile of mRNAsi. We identify shared hub proteins across all quartiles of mRNAsi such as ERCC1, and the connectivity degree for ERCC1 in the first to fourth quartiles are 1.37, 0.37, 0.43 and 1.33. In each quartile, there are also some specific hub proteins, such as P27* in the first and fourth quartiles and RAB11 in the second and third quartiles. Figure 5B shows the heatmap of partial correlations between specific protein pairs with varying mRNAsi. We observe distinct clusters of positive and negative associations on the right and left side of the heatmap respectively. In Figure 5C, we provide the line plots for five selected protein pairs to demonstrate their co-expression patterns with change in mRNAsi, e.g. partial correlations between GAPDH FOXM1 tend to have a positive association with dedifferentiated tumor cells (high mRNAsi levels). GAPDH is also identified as a key factor with respect to proliferation and aggressiveness of breast cancer cells which have significantly positive correlations with higher histoprognostic grades^[52]^. At the same time, FOXM1 is a pro-stemness transcription factor which is suggested to have a driving effect on cell dedifferentiation and proliferation in BRCA^[50]^. Our results are consistent with previous findings and suggest interaction patterns between the two proteins in cell dedifferentiation and proliferation process.

**Figure 5:**
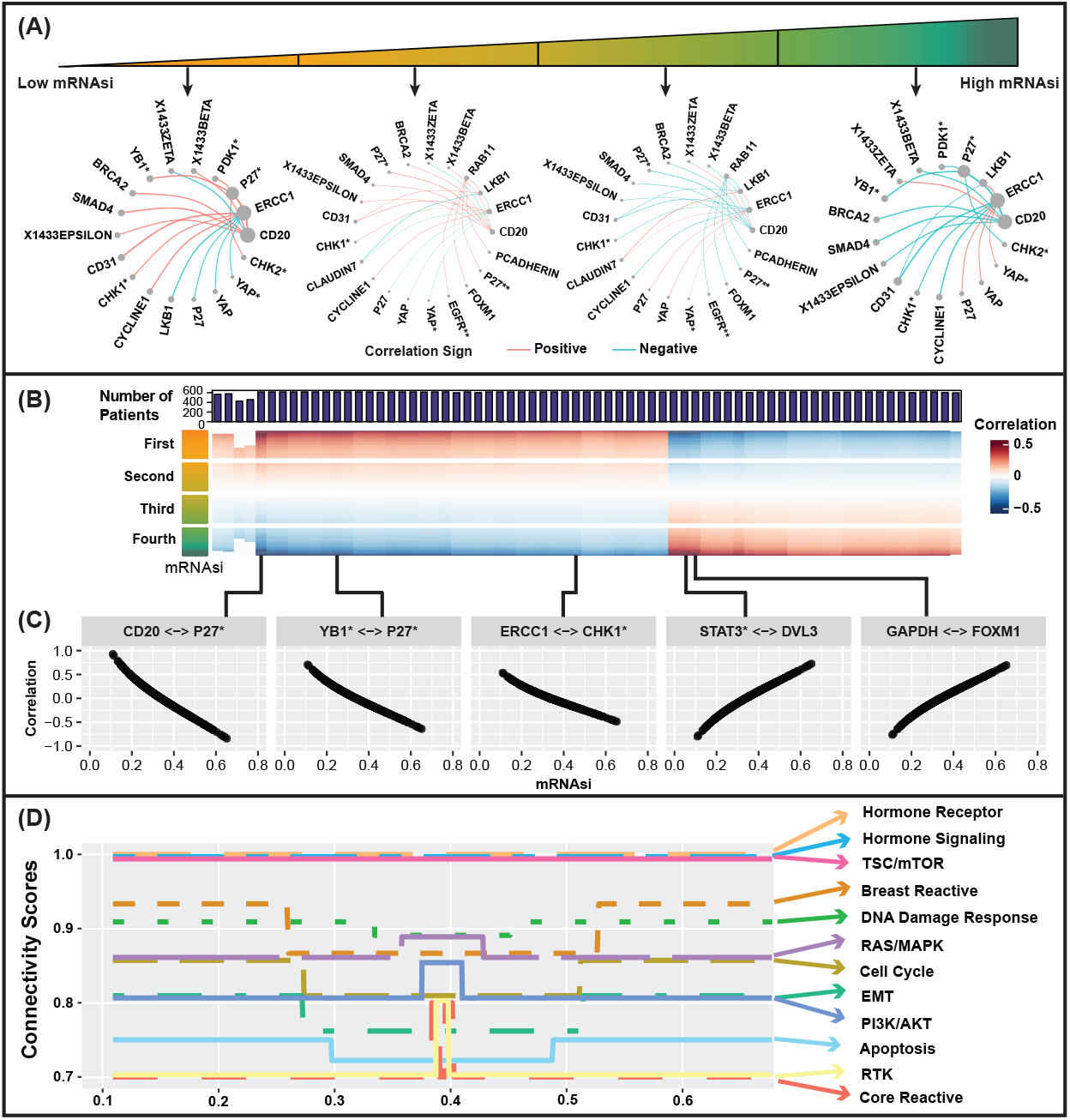
Characterization of Stemness-induced Heterogeneity on Proteomics Networks in Breast Cancer. We apply GraphR with three intrinsic factors including age, logit transformed mRNA and DNA methylation stemness indices (mRNAsi and mDNAsi respectively). We only include protein pairs with significant strong partial correlation (FDR based p-values *<* 0.01) for more than 50% cases and | partial correlation |*≥* 0.45 for at least one case. The results are presented with varying mRNAsi while age and mDNAsi are fixed at median. **(A)** Networks for proteins with top five connectivity degrees corresponded to each quartile of mRNAsi. **(B)** Heatmap of partial correlations between protein pairs with varying mRNAsi. The color bar on the left shows quarterly divided mRNAsi. The barplot on top represents the number of significant cases for the corresponding protein pair. **(C)** Line plots showing association between partial correlation of selective protein pairs and mRNAsi of the original scale. **(D)** Connectivity scores of pathways changing along mRNAsi.

Finally, we conduct a proteomics pathway integrated analysis to show various patterns of connectivity scores along with mRNAsi in Figure 5D, reflecting the heterogeneity of BRCA with the change of dedifferentiation levels. Three pathways – TSC/mTOR pathway, hormone receptor pathway and hormone signaling pathway – are found to be insensitive to the changes of mRNAsi, maintaining a constant CS of one. We also find several pathways varying with mRNAsi, such as breast reactive pathway (CS = 0.87 when mRNAsi is around 0.21 - 0.5; otherwise CS = 0.93); core reactive pathway (CS = 0.8 when mRNAsi is around 0.33 - 0.34; otherwise CS = 0.7). The mDNAsi based analysis is discussed in Section D of the Supplementary Information.

### Pan-Cancer Proteomic Network-based Characterization of Breast and Gynecologic Cancers

The previous two applications mainly focused on the influence of intra-tumor heterogeneity in BRCA. In contrast, this application seeks to elucidate the impact of inter-tumor heterogeneity among BRCA and a selection of gynecologic cancers. Breast and gynecologic cancers exhibit several shared characteristics in terms of biology and evolutionary traits, e.g. their growth and progression are highly correlated with female hormones^[53]^. Despite the molecular-level similarities observed between breast and gynecologic cancers^[54]^, substantial diversities exist among various cancer types, including distinct gene expression patterns that are unique to each type^[53,55]^. In our pursuit of identifying both shared and distinct proteomic features within these cancer types, we employ GraphR to analyze proteomic data from BRCA (n = 892) and three gynecologic cancers: cervical squamous cell carcinoma and endocervical adenocarcinoma (CESC, n = 173), ovarian serous cystadenocarcinoma (OV, n = 436), and uterine corpus endometrial carcinoma (UCEC, n = 440).

Four cancer type-specific networks are shown in Figure 6A with the same inclusion criteria as PAM50 case except magnitude| *ρ* |*≥* 0.45. The network plots of all protein pairs are provided in Figure E2-E5 in the Supplementary Information. We identify the following hub proteins across cancer types, e.g. EGFR* in BRCA and UCEC. EGFR* which has been identified as hub protein in Luminal A, B and Her2-enriched breast cancers, again plays a key role in BRCA and UCEC with the positive partial correlation with Her2* and Src*. Interestingly, MAPK*, a hub protein for Her2-enriched and Basal-like BRCA, is not detected as a hub protein with respect to the combined groups of BRCA, partly due to the fact that Luminal A and B group (sample size(%): 626 (70.17%)) is the dominant subtype in BRCA. Additionally, AMPK*α*\*, a member of AMPK family, is also identified as a hub protein in BRCA and CESC, showing negative partial correlation with ACC1 and positive partial correlation with phosphorylated ACC1 (ACC1*), which aligns with the previous finding that AMPK has been regarded as a key regulator in cellar metabolism which is involved in the phosphorylation of ACC^[56]^. Figure 6B represents the heatmap of partial correlations between selective protein pairs (the corresponding PIP plot is shown in Figure E1 of the Supplementary Information), and the upset plot in Figure 6C summarizes the numbers of significant connections within or across the four cancer types. We find 18 significant connections across all cancer types including connection between mTOR and raptor, EGFR* and HER2*. We also observe several protein pairs with distinguished correlation signs in different cancer types, for example FOXO3A and CD31 display a positive correlation in CESC but a negative relationship in BRCA. In Figure 6D, we present the pathway-related CS for the four cancer types, where-in BRCA has high connectivity scores in hormone signaling (CS = 1) and breast reactive (CS = 0.9) pathways. Simultaneously, the CS of hormone signaling pathway in OV is high (CS = 1).

**Figure 6:**
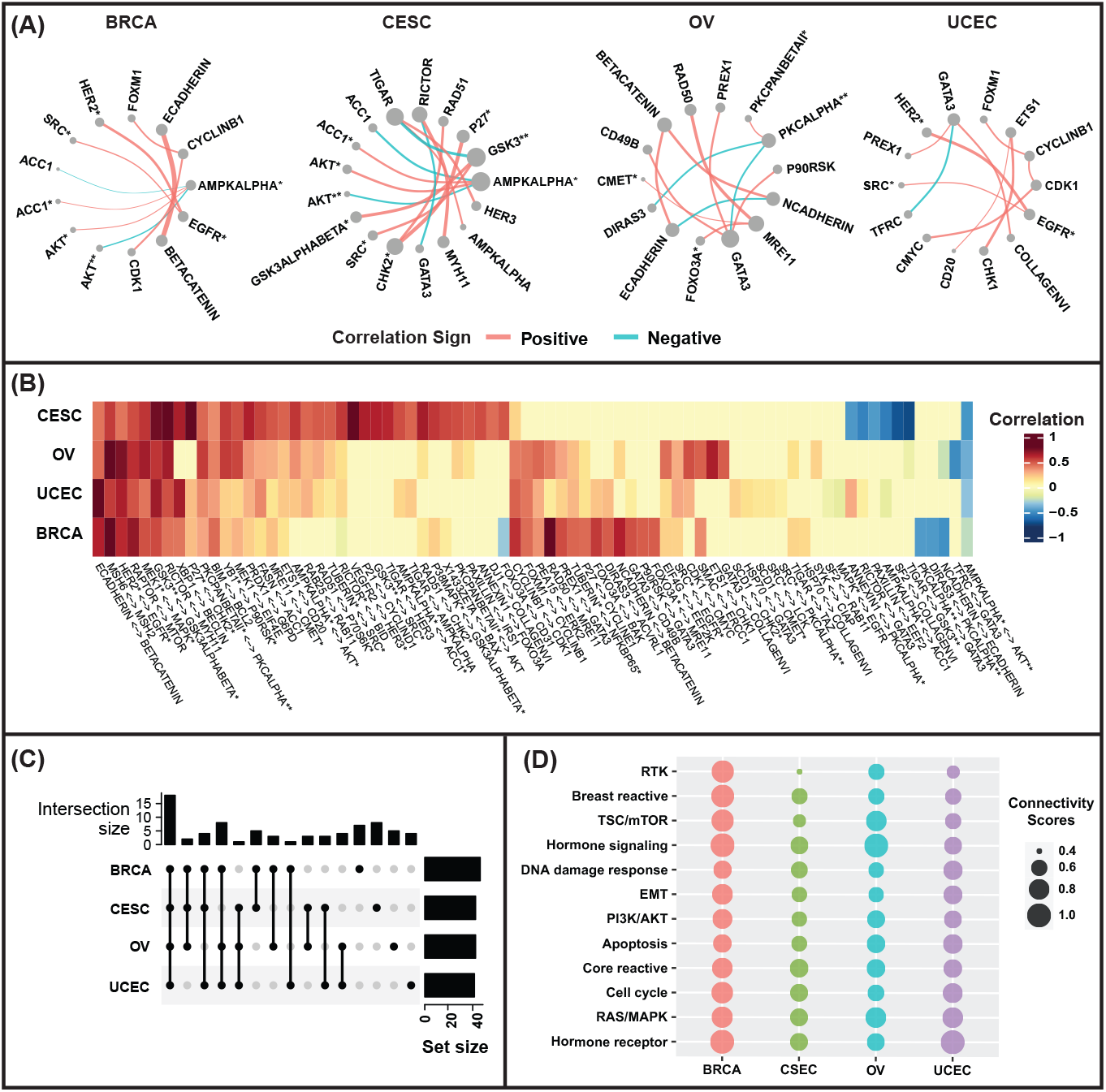
Pan-Cancer Proteomic Network-based Characterization of Breast and Gynecologic Cancers. The results are based on BRCA and three types of gynecologic cancers including cervical squamous cell carcinoma and endocervical adenocarcinoma (CESC), ovarian serous cystadenocarcinoma (OV) and uterine corpus endometrial carcinoma (UCEC). **(A)** Cancer-specific networks for proteins with top five connectivity degrees. **(B)** Heatmap of partial correlations between protein pairs for each cancer type. We only include significant protein pairs which are highly correlated with FDR based p-values *<* 0.01 and | partial correlation | *>* 0.45 for at least one type of cancer. **(C)** Upset plot for numbers of significant connections within or across cancer types based on the protein pairs included in (B). **(D)** Connectivity scores of pathways for each cancer type. Pathways are ordered based on the clustering of connectivity scores.

### Gene Expression Network Estimation incorporating Spatial Heterogeneity

We demonstrate the usage of GraphR in the estimation of gene expression networks incorporating spatial heterogeneity. We use a spatial transcriptomics dataset in BRCA, characterized by simultaneous measurements of gene expression and spatial coordinates which serve as intrinsic factors in the model. Our underlying hypothesis is that nearby spots/cells will share “closer” connectivity than farther ones. We utilize a human breast cancer dataset collected from a BRCA biopsy at a thickness of 16 *µ*m^[57]^. Based on the Hematoxylin and Eosin staining image in Figure 7A (Top), we classify locations into three spatial regions: tumor, intermediate, and normal with the sizes 114, 67, and 69 spots respectively (Figure 7A Bottom). GraphR is implemented to the data which contains measurement of 100 spatially expressed genes at 250 spot locations to obtain spatially dependent networks (preprocessing steps are deferred to Section F2 of the Supplementary Information). All the results are explained with respect to intermediate region which serves as the baseline.

**Figure 7:**
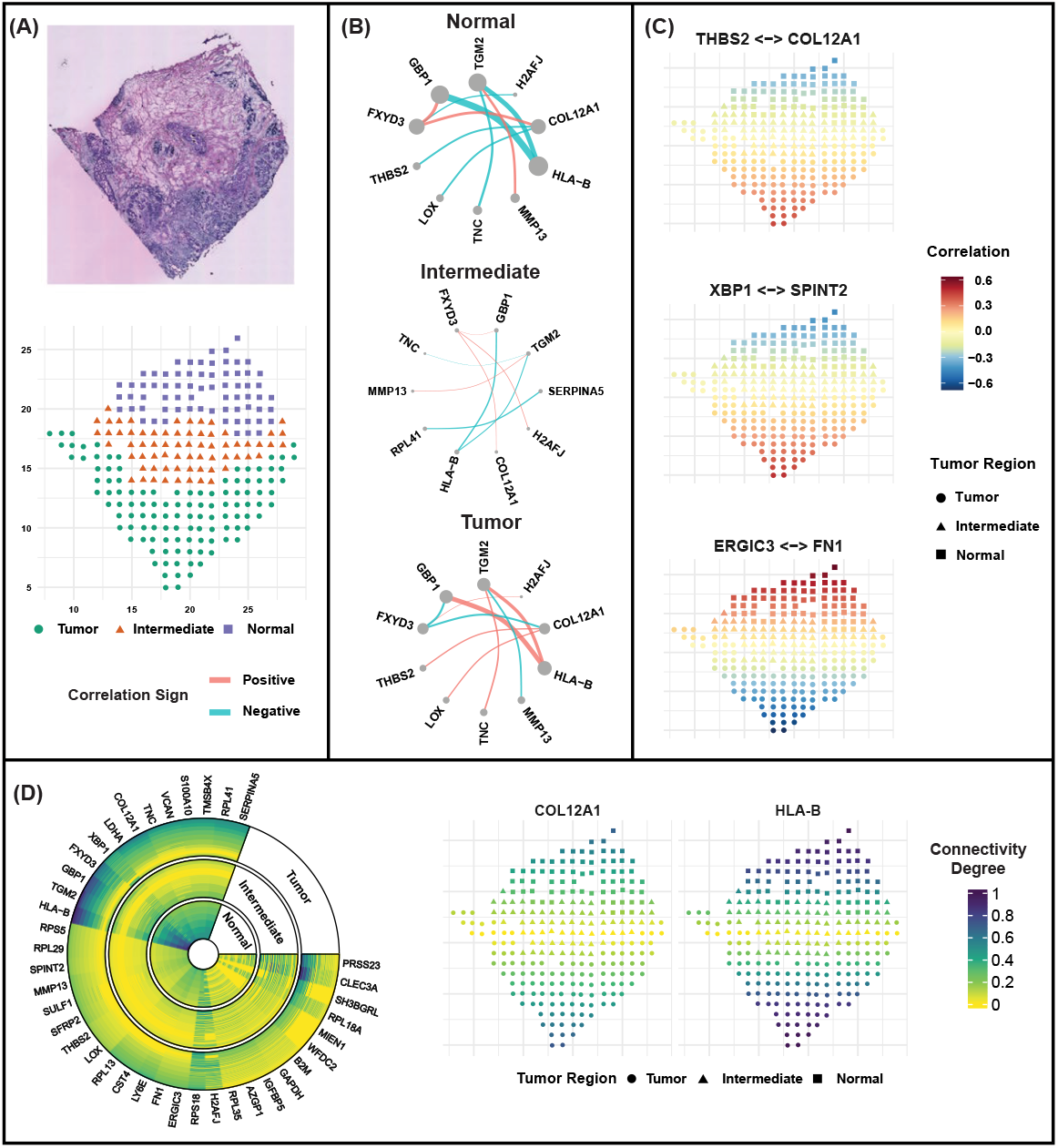
Gene Expression Network Estimation incorporating Spatial Heterogeneity. **(A)** The hematoxylin and eosin staining image (Top) and the spatial distribution of measured spots (Bottom) which are classified into tumor, intermediate and normal region^[57]^. **(B)** The networks of genes with top five weighted connectivity degrees on the right w.r.t. each spatial region. Weighted connections of each spatial region are defined as the weighted mean of significant partial correlations (FDR based p-values *>* 0.01) for all spots in the corresponding region with weights being posterior inclusion probability of these connections. Width of edges represent weighted partial correlations and sizes of nodes represent weighted connectivity degrees **(C)** Spatial pattern of partial correlations for selective gene pairs (THBS2 COL12A1; XBP1 SPINT2; ERGIC3 FN1) where point shapes defer for each spatial region. **(D)** Circular heatmap of connectivity degrees of genes for each cluster (left) and spatial pattern of connectivity degrees of selected genes COL12A1 and XBP1 (right).

Figure 7B shows the networks of genes with the top five weighted connectivity degrees, defined as weighted mean of significant correlations (details in Section F3 of the Supplementary Information) within each spatial region. Based on the networks, we find several hub genes shared across spatial regions, e.g. HLA-B (weighted connectivity degrees: 0.94 for tumor; 0.23 for intermediate; 1.35 for normal) which shows negative partial correlation with GBP-1 in normal region and positive in tumor region. In addition to the shared genes, we also find several genes specific to each region, e.g. COL12A1 in normal and tumor regions (weighted connectivity degrees: 0.54 for tumor; 0.76 for normal).

We present the spatially varying partial correlations of selected gene pairs in Figure 7C, demonstrating the ability of GraphR to incorporate spatial information and explicitly estimate spatially varying dependence structures. We observe negative partial correlations in the normal region and positive partial correlations in the tumor region for gene pairs THBS2 - COL12A1 and XBP1 SPINT2 and the inverse partial correlation pattern for ERGIC3 - FN1. The spatial pattern between XBP1 - SPINT2 aligns with previous findings^[58]^ which have identified SPINT2 as a hub gene for tumor and intermediate regions whereas XBP1 is a hub gene for tumor region only. XBP1 can induce cell invasion and metastasis in breast cancer cells and high expression of XBP1 can promote the proliferation of tumour cells^[59]^. We further show the overall connectivity degree patterns for selected genes via circular heatmap (Figure 7D left). Genes such as HLA-B, TGM2 and GBP1 tend to exhibit high connectivity degrees across most tumor and normal regions. Meanwhile, there are no clear patterns for some genes within each cluster, such as CLEC3A and H2AFJ, partly reflecting the spatial heterogeneity of cancers. The right panel of Figure 7D displays the spatial distribution of connectivity degrees with respect to COL12A1 (hub genes in normal and tumor regions) and HLA-B (hub genes in all regions). Networks containing all included proteins for each region (Figure F1-F3) and the spatial pattern of partial correlations (Figure F4) and connectivity (Figure F5) degrees for more gene pairs are discussed in Section F of the Supplementary Information.

## Discussion

We present GraphR, a scalable and adaptable probabilistic graphical modeling framework designed to infer heterogeneous networks for high-dimensional multivariate data. Standard GGM approaches assume homogeneity among samples, failing to incorporate intrinsic heterogeneity and potential confounding variables, which might lead to erroneous conclusions as illustrated via the Simpson’s paradox. To this end, GraphR enables seamless integration with intrinsic factors which simultaneously accounts for group-specific or sample-specific heterogeneity and thus providing more precise and robust estimation of graph structures. Furthermore, GraphR employs sparsity-inducing priors facilitating efficient estimation with limited sample sizes along with providing uncertainty quantifications (controls multiplicity) on the selection of both edges and intrinsic factors. As we show through multiple simulation settings, this achieves the right balance between sensitivity and specificity – high accuracy in graph estimation along with low false positive rates – compared to multiple other competing methods.

GraphR can empower researchers to obtain a more systematic and unbiased understanding of the underlying biological mechanisms and their changes along with inter- and intra-tumor heterogeneity in tumor microenvironment. For example, the connectivity levels of RAS/MAPK pathway are found to be relatively high in Luminal A and B and Basal-like BRCA. It has been shown that RAS/MAPK pathway is closely associated with BRCA, and several FDA-approved prescription drugs that regulate or inhibit the MAPK pathway including Farnesyltransferase inhibitors and Sorafenib may reduce metastatic progression in BRCA^[60]^, especially for Basal-like BRCA^[61,62]^. Furthermore, the RAS/MAPK pathway is also shown as a critical signaling pathway involved in various cellular processes, including cell differentiation, proliferation, and apoptosis^[63]^, which partly explains the changing pattern of CS along with the stemness indices and thereby implies potential variations in behaviors among BRCA characterized by varying levels of differentiation. Beyond BRCA, we also find RAS/MAPK pathway has relatively high connectivity scores in UCEC and OV. In the case of UCEC, insulin has been linked to tumor development through the PI3K/Akt and RAS/MAPK pathways^[64]^, which further elaborates the potential anticancer effects of metformin, a common treatment for type II diabetes, through RAS/MAPK pathway^[65,66]^. Furthermore, the elevated CS observed in OV aligns with previous research suggesting that inhibitors of the MAPK/RAS may improve patient survival^[67]^.

In the context of tumor microenvironment, GraphR can be used to infer spatially varying networks and subsequently identify hub genes and connections which are specific to regions or cell types when applied to spatial transcriptomic data. In our breast cancer example, We have found HLA-B as a hub gene across all regions and is negatively correlated with GBP1 in normal region and positive partial correlation in tumor region. HLA-B, belonging to MHC class I, has been shown to be associated with favorable outcome in breast cancers through increased immune T cell activation^[68,69]^. Moreover, GBP-1, has also been identified as a regulator of T cell activation^[70,71]^. GBP-1 displays relatively strong expression in infiltrating cells while exhibiting weak expression in tumor cells^[72]^, which elucidates the spatial interaction pattern between HLA-B and GBP-1.

A major advantage of GraphR, over existing Bayesian approaches that allow for the estimation of covariate-dependent graphs^[35,33]^, is its computational efficiency. Instead of applying MCMC, one of the gold standards for posterior inference, we apply variational Bayes algorithms with exchangeable priors and the mean-field assumption^[73,74]^ in GraphR, which significantly improves the scalability in high-dimensional settings. For instance, a full Bayesian estimation^[33]^ of 50-node networks with 4 intrinsic factors required 15.6 hours as opposed to 9.32 seconds using a high-computing cluster with single CPU core (see <u>Methods Section</u> for further computational comparisons). This promising feature positions GraphR for extensive use in biomedical research, especially when dealing with high-throughput multi-omics data. To facilitate further analysis in the biomedical research field, we have developed the *<u>GraphR</u>* <u>package</u>. This package provides functions for estimation, prediction and network visualization. Additionally, we have constructed a *<u>GraphR</u>* <u>Shiny App</u> that provides users with a dynamic platform for both exploring and visualizing graph patterns using datasets used in this paper, as well as applying the GraphR method to their own datasets, all without the need for prior knowledge or coding in R.

The GraphR method can be extended in several directions. One can use a non-Gaussian distribution to model graphical structures for discrete data. Another interesting avenue is the extension to non-linear function mapping between covariates and precision matrix^[33]^. The GraphR method also holds potential for expansion to the study of spatial genomic and proteomic networks across multiple tissue samples. We posit that these extensions will likely improve broad applicability of GraphR.

## Methods

### GraphR Overview and Construction

#### Preliminaries

Gaussian graphical models are standard approach to study dependent structure of multivariate data. A p-dimensional graph for vector **Y** = (*Y*_1_, …, *Y*_*p*_) *∈* ℝ ^*p*^ is defined with a tuple *G*_*y*_ = *{G, P*(**Y**)*}*, where *G* is the corresponding graph with associated distribution *P*(**Y**). A graph *G* = (*V, E*) contains a vertex set *V* = *{*1, …, *p}* and an edge set *E ⊂ V × V*, representing conditional relationship among each vertex. For an undirected graph, an edge is included if (*i, j*) *∈ E ⇔* (*j, i*) *∈ E*. A Gaussian graphical model is defined by setting the associated distribution *P*(**Y**) to a Gaussian distribution with mean *µ ∈* ℝ^*p*^and covariance matrix Σ *∈* ℝ^*p×p*^ i.e. **Y** *∼ N* (*µ*, Σ). We denote the precision matrix with Ω = Σ^*−*1^ = (*ω*_*ij*_)_*p×p*_ which is a positive definite (PD) matrix. The partial correlation between random variable *Y*_*i*_ and *Y*_*j*_ is represented as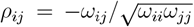. Notably, an undirected edge exits iff (*a*) *ρ*_*ij*_ ≠ 0 and (*b*) *ρ*_*ij*_ = *ρ*_*ji*_ (reason being (*i, j*) *E* (*j, i*) *E*). The condition in (*b*) is not necessary for a directed graph. One can infer the conditional graph structure based on the estimated precision matrix Ω.

To maintain the parsimonious structure of the network model, we introduce sparsity in the partial correlation. The non-zero entries *ω*_*ij*_ in the precision matrix correspond to the existing edge between *Y*_*i*_ and *Y*_*j*_. Furthermore, under the normality assumption, zero entries of the precision matrix indicate the conditional independence between the corresponding nodes given all other variables *Y*_*−*(*i*,*j*)_. Thus learning the conditional independence structure is equivalent to estimating and identifying zero entries in the precision matrix, which is also known as covariance selection problem. Under the assumption of Gaussian distribution, *Y*_*i*_ ∑ *j?* ≠ *i γ*_*ij*_*Y*_*j*_ + *ϵ*_*i*_, with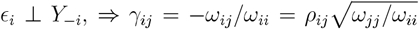, allows to connect identification of zero entries in precision matrix to a coefficient selection problem in a regression setting^[11]^. Based on the restated problem, a non-zero *γ*_*ij*_ will lead us to the existence of an edge i.e. (*i, j*) *∈ E*.

In practice, it is common to have the case where conditional independence pattern for each individual has structural differences which can be reflected by some intrinsic factors, such as cancer types or continuous biomarkers. Previous research works^[33,35]^ have shown that given intrinsic factors **X**, the pairwise Markov property holds while modeling the dependent structure as a function of intrinsic factors in both directed acyclic and undirected graphical settings. Suppose precision matrix is a function of intrinsic factors **X** = (*X*_1_, …, *X*_*q*_) *∈* ℝ^*q*^, denoted as Ω(**X**) = [*ω*_*ij*_(**X**)]_*p×p*_. We will have *Y*_*i*_ *⊥ Y*_*j*_|*Y*_*−*(*i*,*j*)_, **X** if *ω*_*ij*_(**X**) = 0. For simplicity and interpretation, we here only consider the situation where diagonal entries of precision matrix are independent of intrinsic factors while off-diagonal entries are linear function of intrinsic factors. This aligns with the covariate dependent varying-coefficient model^[35]^ which is an extension of the covariance selection problems^[11]^.

#### GraphR Model

Suppose we have a random vector **Y** = (*Y*_1_, …, *Y*_*p*_), and intrinsic factors **X** = (*X*_1_, …, *X*_*q*_) with *N* samples of (**Y, X**). We regress *Y*_*i*_ (1 *≤ i ≤ p*) on other nodes *Y*_*−I*_ where regression coefficients are function of intrinsic factors, namely,

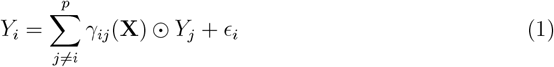

Here *⊙* represents the element-wise multiplication between vectors. Under the normality assumption, we have *ϵ*_*i*_ is independent of *Y*_*j*_ given X if and only if *γ*_*ij*_(**X**) = *−ω*_*ij*_(**X**)*/ω*_*ii*_(**X**) where *ω*_*ij*_(**X**) and *ω*_*ii*_(**X**) being the off-diagonal and diagonal entries of precision matrix Ω(**X**) and *ϵ*_*i*_ ∼ *N* (0, 1*/ω*_*ii*_). As we only consider the case where diagonal entries of precision matrix are independent of intrinsic factors while off-diagonal entries are linear function of intrinsic factors, we can further express the *γ*_*ij*_ as:

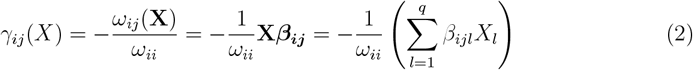

where ***β***_***ij***_ = [*β*_*ij*1_, …, *β*_*ijq*_]^*T*^ are coefficients of intrinsic factors.

Notably, the functional form of *γ*_*ij*_(**X**) enables the flexibility in our models, with the following special cases (I) we have a standard Gaussian graphical model if **X** = (1, …, 1)^*T*^; (II) multiple/group-specific Gaussian graphical models are obtained when **X** is categorical; and (III) with a continuous covariate, our model can be viewed as a Gaussian graphical model varying over the domain of the continuous covariate. Moreover, if there is prior knowledge on the directionality in the nodes **Y**, the methodology can be adapted to a directed acyclic graph (DAG) models along with the incorporation of sample heterogeneity (via **X**).

#### Priors

As opposed to GGM models that estimate networks of order *𝒪*(*p*^2^) parameters, GraphR network parameters are on the order of *𝒪* (*p*^2^*q*). Hence, we need to induce sparse estimation especially with limited sample sizes (relative to number of parameters). Furthermore, we assume a covariate dependent sparse precision matrices since genomic and proteomic networks are typically sparse in nature. To this end, note that the coefficients *β*_*ijl*_ = 0 for all *ℓ* = 1, …, *q* implies the edge between random variable *Y*_*i*_ and *Y*_*j*_ doesn’t exist which leads to a sparse linear regression problem. From a Bayesian perspective, the most widely used sparse regression technique is to impose a model selection prior, such as spike-and-slab prior, which assigns probabilistic weights to each variable and shrinks noise variables to exact 0^[75]^. It yields automatic edge selection property for graphical models by providing exact zero estimates for several variables, as well as inherits some desirable theoretical aspects^[76,77]^. Therefore we employ a spike-and-slab on the coefficient of intrinsic factors with different slab precision and mixing parameters, as follows:

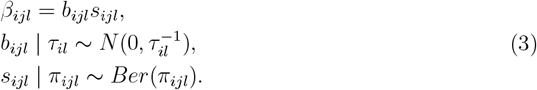

With probability *π*_*ijl*_, *β*_*ijl*_ are drawn from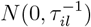 i.e. “slab” density with precision parameter *τ*_*il*_, and *β*_*ijl*_ is 0 with probability 1 *π*_*ijl*_. More specifically for *π*_*ijl*_ = 1, we will have a normal prior on the coefficient of intrinsic factor, which is equivalent to ridge regression^[78]^. Conjugate priors for hyper-parameters *τ* and *π* are selected:

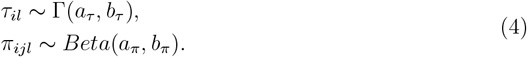

We chose a non-informative prior for variance parameter *ω*_*ii*_ *∝* 1^[35]^.

### Model fitting via variational Bayes

MCMC is a standard technique for posterior inference of model parameters^[79,80]^. However, since the posterior distribution contains high dimensional integrals and 2^*p∗*(*p−*1)*∗q*^ potential models, MCMC will be computationally challenging for large, even moderate p and q. Throughout this article, we consider a VB method^[73,74]^ with exchangeable priors and mean-field assumption. Mean-field VB (MFVB) aims to find an approximation belonging to a family of distributions and the posterior distribution by minimizing the Kullback–Leibler divergence. The target distribution is assumed to have independent partitions such that parameters with strong dependencies, i.e., magnitude parameters *b* and selection indicators *s* in our case, will be estimated jointly. More details about implementation of the VB method and posterior inference are provided in Section A of the Supplementary Information. The GraphR algorithm is also provided in Section A.5 of the Supplementary Information.

### Network Construction and Summaries

#### Network Construction

Networks are constructed for each group/individual via: (1) posterior expectation of magnitude (E*b*) and (2) posterior inclusion probability (PIP, E*s*) for each coefficient and (3) expected precision parameter (𝔼*ω*_*ii*_) obtained from GraphR results. Considering an individual with intrinsic factor (*x*_1_, …, *x*_*q*_), we define PIP of the edge between node *j* and *i* as a weighted mean of 𝔼(*s*_*ijl*_) _1*≤l≤q*_ with weights proportional to (*x*_*l*_𝔼*b*_*ijl*_)^2^. An (significant) edge is included in the network if the corresponding PIP is larger than a FDR based cutoff^[81,82]^. Partial correlation is calculated as 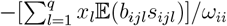, indicating the sign and strength of the corresponding edge.

#### Network Summaries

Downstream analyses, e.g. hub node detection and pathway-based functional analysis, are conducted based on constructed networks. Hub genes are detected based on the *connectivity degree* which is defined as the sum of magnitudes of significant edges (i.e. absolute values of partial correlations) between the corresponding node and others. Aiming to evaluate the connectivity status of a pathway, we apply *connectivity score (CS)*, which is calculated as the ratio of the observed number of edges between proteins belonging to the corresponding pathway to the total number of possible edges.

**Simulated Data** In this section, we assess the performance of the GraphR method in different simulation settings from undirected and directed graphs. In the first scenario, we consider undirected graphs with unordered pairs of vertices, and directed graphs in the second scenario where directions of edges are decided by some prior knowledge. For both settings, the numbers of parameters to be estimated are *𝒪* (*p*^2^*q*) which are significantly larger than sample sizes. We focus on graph structural recovery across different settings.

#### Comparative Methods

We compare our approach to six relevant methods: (I) BGGM^[19]^ (II) GLASSO^[14]^ (III) FGL^[36]^ (IV) GGL^[36]^ (V) LASICH^[37]^ (VI) K-GLASSO^[31]^. Under categorical intrinsic factors, the proposed GraphR is equivalent to multiple graphical model which assumes that samples are drawn from *G* distinct groups. Thus, we compare our method with the following three commonly used methods for multiple graphical models: FGL, GGL, and LASICH. In case of continuously varying graph structures over the domain of continuous intrinsic factors, we compare our method with BGGM, GLASSO, and k-GLASSO. K-GLASSO integrates intrinsic factors by kernel smoothing function and is regarded as a fair comparison in terms of GLASSO and BGGM. A standard Gaussian graphical model can be treated as a special case of GraphR with intrinsic factors being intercept only. For comparison, we consider one Bayesian and one non-Bayesian approach (i.e. BGGM and GLASSO respectively), and both methods are designed for independent and identically distributed subjects from homogeneous populations. Other than the methods mentioned above, there are more methods that can be used to infer the graph structure varying with intrinsic factors, such as undirected Gaussian graphical models conditional on covariates (GGMx,^[83]^) and Bayesian Edge Regression^[35]^. However, these approaches use MCMC algorithm leading to the lack of scalability for large even moderate *n* and *p*, and thus are excluded from the comparison methods.

#### Comparative Measures

We use the following comparative measures to evaluate the graph structural recovery performance across different settings. We calculate the true positive rate (TPR), false positive rate (FPR), false discovery rate (FDR) and Matthews correlation coefficient (MCC),

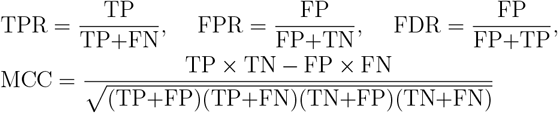

where TP, FP, TN, and FN denote true positives, false positives, true negatives, and false negatives. MCC^[84]^ measures the quality of binary classification, ranging from +1 (perfect classification) to -1 (total mismatch). In addition to the previous measures, we construct the receiver operating characteristic (ROC) curve to compare the settings ability to recover significant edges. The sensitivity (TPR) and 1-specificity (FPR) for each setting are computed at each threshold parameter value [*α∈* (0, 1)] with the area under the ROC curve (AUC) to compare the structural recovery in each setting. Higher AUC values imply better recovery of edges. Specially we calculate the measures for each subject in the continuous case and mean values of measures are reported. Next, we discuss the simulation design and performance of the GraphR method w.r.t. undirected and directed graphs.

**Scenario I: Undirected Graphs** We consider undirected graphs with unordered pairs of vertices in the first scenario which contain three simulation settings. In each setting, we fix sample size *N* = 151, number of nodes *p* = 33, and connection probability *π* = 2%, with the types and numbers of intrinsic factors changing with different cases. In <u>case A</u>, we consider discrete intrinsic factors which reduce the GraphR to multiple graphical models. <u>Case B</u> is comprised with individual specific GGM with continuous intrinsic factors. We assume that all individuals are homogeneous in <u>case C</u>. The detailed simulation design of cases A, B, and C are discussed in Section B.1.1, B.1.2, and B.1.3 of the Supplementary Information respectively.

In <u>Case A</u>, all four approaches yield relatively high AUC as reported in the *<u>Results</u>* <u>Section</u>. TPR for FGL, GGL and LASICH are higher than GraphR (GrphR: 0.856, FGL: 0.988, GGL: 0.989, LASICH: 0.985), but at the cost of large values of FDR (<u>GraphR: 0.062</u>, FGL: 0.662, GGL: 0.662, LASICH: 0.602) and FPR (<u>GraphR: 0.001</u>, FGL: 0.047, GGL: 0.047, LASICH: 0.038). GraphR provides the lowest FDR and FPR values which are almost nine times smaller than FGL, GGL, and LASICH. Figure 3A shows the MCC from GraphR is higher than other three methods (values mentioned in *<u>Results</u>* <u>Section</u>), implying a better performance in terms of graph structural recovery.

In <u>Case B</u>, we have a heterogeneous structure with individual specific precision matrix. The GraphR significantly outperforms three comparison methods in such heterogeneous structure with respect to all measures. Figure 3B denotes the outstanding overall performance in terms of AUC and MCC. Moreover, GraphR achieves high TPR (<u>GraphR: 0.992</u>, BGGM: 0.374, GLASSO: 0.525, k-GLASSO: 0.304) accompanying with well-controlled FDR (<u>GraphR: 0.076</u>, BGGM: 0.61, GLASSO: 0.779, k-GLASSO: 0.589) and FPR (<u>GraphR: 0.001</u>, BGGM: 0.011, GLASSO: 0.028, k-GLASSO: 0.008) values in comparison with BGGM, GLASSO, k-GLASSO.

In <u>Case C</u>, with homogeneous graph for all individuals, GraphR achieves marginally similar TPR (GraphR: 0.966; BGGM: 0.992; GLASSO: 0.998) alongwith the lowest values of FDR (GraphR: 0.052, BGGM: 0.084, GLASSO: 0.567) and FPR (GraphR: 0.001, BGGM: 0.002, GLASSO: 0.03). In Figure 3C, we observe that the MCC value of GraphR is higher than BGGM and GLASSO (values reported in *<u>Results</u>* <u>Section</u>), leading to a better performance of GraphR in overall measurement.

To summarize, the robust GraphR method achieves high graph structural recovery rates and controls FDR across all three cases with different settings of covariance matrices. GraphR achieves the highest MCC values in case of heterogeneous structure at the presence of discrete or continuous intrinsic factors which are routinely present in various omics datasets. Additionally, GraphR has more power than any other methods, with the lowest FDR and FPR. This indicates that incorporating intrinsic factor-based information leads to better structural recovery.

### Scenario II

**Directed Acyclic Graphs** In scenario II, we consider a special case of our GraphR in which pre-knowledge based directions of edges are assumed, leading to a directed acyclic graph model. We consider different simulation settings to illustrate the performance of variable selection and scalability in case of moderate and/or large number of nodes (p) and intrinsic factors (q). Details of the simulation design are deferred to Section B.2 of the Supplementary Information.

In general, GraphR performs better as the effect size (*β*) and the ratio between *n* and *pq* increase. In a moderate scenario where p=50 and q=2, GraphR has an average performance with a ratio of 1. However, our method’s accuracy improves when the ratio is greater than or equal to 2 and *β* equals 3, especially for MCC and TPR. For instance, when there is one continuous and one discrete intrinsic factor, the values of MCC and TPR for a ratio of 1 are 0.66 and 0.56, respectively. When the ratio is 2, the values jump to 0.87 for both, indicating significant improvement. Moreover, GraphR tends to perform slightly better with more discrete than continuous intrinsic factors when *n/pq* are not large, ie when ratio = 1, MCC for two discrete, one discrete, and one continuous, and two continuous intrinsic factors are 0.71, 0.66 and 0.56 respectively. However, we don’t detect the impact of types of intrinsic factors on the performance of GraphR when *n/pq* is relatively large. As shown in Figure 3D,3E, 3F, when ratio is equal to five, all five measures for models with different types of intrinsic factors are close to each other, with AUC and TPR close to 1, MCC ranging from 0.89 to 0.91, FDR around 0.17 and FPR smaller than 0.01. We also find a similar pattern for the setting with large number of nodes (Section B.2 of the Supplementary Information). With ratio *n/pq≥* 2, GraphR achieves good performance with respect to almost all criteria, except for FDR which has a relative increase when ratio is three. In case of relatively large number of intrinsic factors, GraphR performs well (e.g., *β* = 2 and *n/pq* = 5), with MCC values of 0.84 and 0.71 with the five or ten intrinsic factors setting. Additionally, as the number of intrinsic factors increases, more edges are considered to be significant which results in increased TPR and FPR.

**Computational efficiency** In all cases, GraphR achieves improved performance in computation time which is expected to have a linear relationship with sample size n, number of nodes p, and number of intrinsic factors q. On average, computation times for case A (one intrinsic factor), B (two intrinsic factors) and C (two intrinsic factors) are 0.978s, 1.848s and 2.055s, which are slightly longer than these frequentist approaches such as GLASSO (0.484s for case A and 0.587 for case C), FGL (0.154s for case B) and GGL (0.155s for case B) in a high-computing cluster with single CPU core. However, compared to BGGM and LASICH which take around 13 seconds, GraphR achieves better efficiency in computation. More time comparisons are deferred to Section B of the Supplementary Materials.

GraphR is also competitive in computation efficiency. For instance, bayesian graphical regression^[33]^, which applies MCMC to infer structure of directed acyclic graphs with intrinsic factors, spends 15.6 hours on case with n = 200, p = 50, and q = 4. However, in the case where n = 5000, p = 500, and q = 2, leading to estimation of 499,000 parameters and 250,500 hyperparameters, the average computation time for GraphR is less than 50 seconds, showing a large improvement in scalability.

## Data Availability

Datasets analyzed in this paper are publicly available. The proteomics data are obtained from patients involved in TCGA^[42]^, and can be downloaded at tcpaportal.org/tcpa/ download.html. The demographic information including intrinsic subtypes of breast cancer, cancer types, and ages can be downloaded by using the “TCGAbiolinks” R package www.bioconductor.org/packages/release/bioc/html/TCGAbiolinks.html. Stemness indices are available at gdc.cancer.gov/about-data/publications/PanCanStemness-2018. Spatial transcriptomics data from human breast cancer are available at www.spatialtranscriptomicsresearch.org.

## Code Availability

The GraphR package is available at github.com/bayesrx/GraphR and the shiny app is available at bayesrx.shinyapps.io/GraphR/.

## Acknowledgements

Veera Baladandayuthapani was supported by NIH grants R01-CA160736, R01CA244845-01A1, and P30 CA46592, NSF grant 1463233, and start-up funds from the U-M Rogel Cancer Center and School of Public Health. Satwik Acharyya was supported by the Postdoctoral Fellows Grant of the University of Michigan Rogel Cancer Center which helps us to host the shiny app server. Yang Ni was supported by NIH grant 1R01GM148974-01 and NSF grant DMS-2112943.

## Additional information

**Supporting Information** file is available <u>here</u>.

## Author Contributions

V.B. conceived the idea and provided funding support. L.C., S.A. and V.B. designed the experiments. L.C. and S.A. developed the method, implemented the software, performed simulations and analyzed real data. C. L. developed the accompanying Shiny App. L.C., S.A., Y.N. and V.B. wrote the manuscript.

## Footnotes

1 Note: Throughout this text, we denote phosphorylated forms of proteins with an asterisk (*)

