## Supplementary Information for "Probabilistic Graphical Modeling under Heterogeneity"

#### Introduction

Network modeling are widely used in biomedical research, aiming to estimate and visualize complicated dependency structures in various fields and at different level. Graphically, networks compromise a set of variables (nodes) and relationships among nodes which are referred as edges. Under the assumption that: (1) edges represent partial correlation between nodes; (2) nodes follow Gaussian distribution, leading to a Gaussian graphical models (GGM)[7]. GGM can be represented as a multivariate Gaussian distribution, usually with a sparse precision matrix of which a zero entry is equivalent to conditional independence. Most current probabilistic GGM-based methods assume homogeneous samples which limits the applicability of these models to incorporate heterogeneity across samples that is routinely present in many scientific contexts. We propose a flexible and computationally efficient approach called Graphical Regression (GraphR) which allows for covariate-dependent graphs and enables incorporation of sample heterogeneity. The Figure below provides an overview of the GraphR method.

Here we provide supplementary materials for Probabilistic Graphical Modeling under Heterogeneity, which are organized as following:

1. In Section A, we provide a detailed introduction for mean-field variational Bayes method and the derivation of the methodology.

2. In Section B, additional simulation results for undirected and directed settings are discussed.

3. In Section C, we present the additional results from the PAM50 protemoics dataset.

*These authors contributed equally.

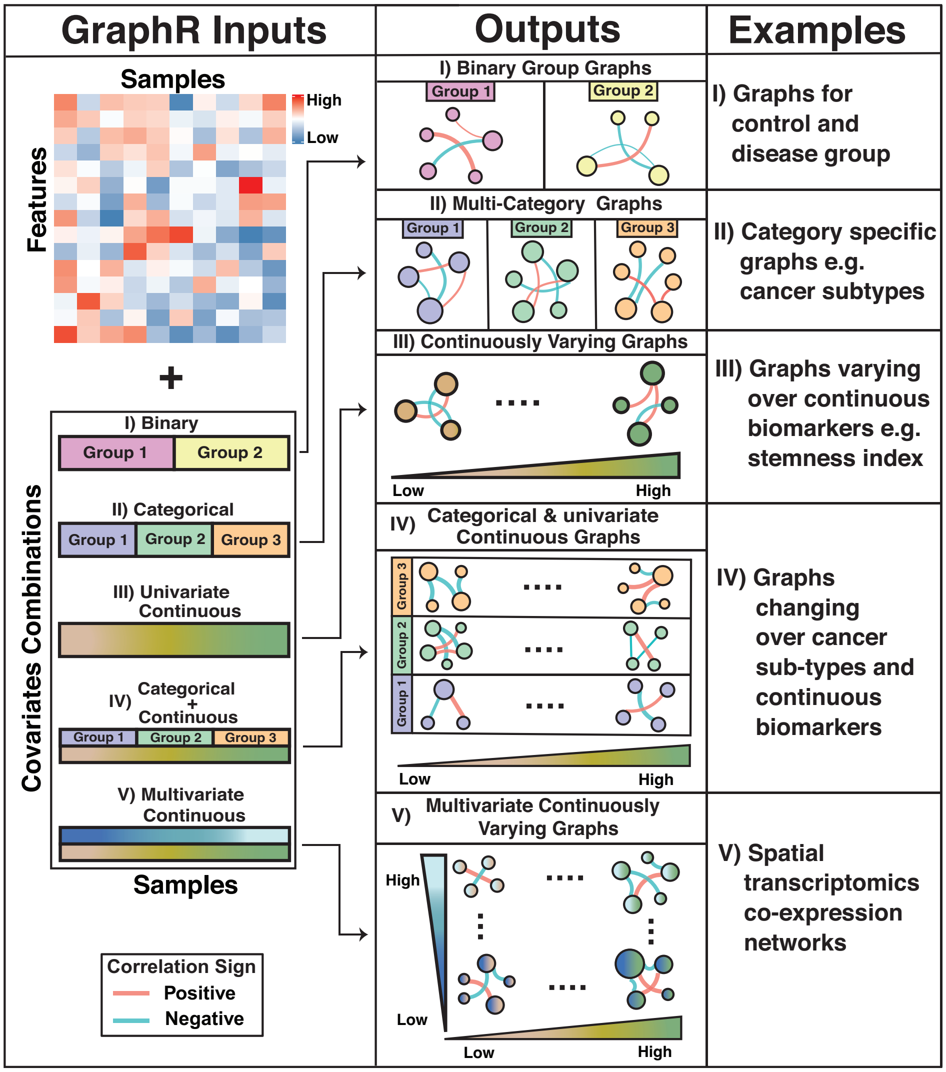


4. In Section D, more results from stemness and age based breast cancer data are provided.

5. In SectionE, we added further analysis from pan-gynecological breast cancer data.

6. In Section F, additional results from from spatial transcriptomics breast cancer data are presented.

7. In Section G, we layout the implementation related details of the GraphR package.

### A Methodology

In this Section, we discuss the GraphR model and priors in Section A.1 and brief introduction to variational Bayes inference method with mean-field assumption in Section A.2 followed by detailed derivation of the update equations and evidence lower bound (ELBO) for our GraphR model in Section A.3 and Section A.4 respectively. In Section A.5, we provide a comprehensive overview of the GraphR and competing methods. Algorithm can be found in Section A.6.

For notational purposes, we consider $\mathbf{Y} = (Y_1, ..., Y_p) \in \mathbb{R}^{n \times p} \sim \backslash(\mu, \Omega^{-1}(\mathbf{X}))$, where the precision matrix $\Omega(\mathbf{X}) = [\omega_{ij}(\mathbf{X})]_{p \times p}$ is a function of intrinsic factors $\mathbf{X} = (X_1, ..., X_q) \in \mathbb{R}^{n \times q}$ and denote $\boldsymbol{\theta}$ as the parameters of estimation. Here $n$, $p$, $q$ denotes sample size, number of features and covariates respectively.

#### A.1 Model and priors

The GraphR model is expressed as follows:

$$
\begin{aligned}
Y_i &= \sum_{j \neq i}^{p} \gamma_{ij}(\mathbf{X}) \odot Y_j + \epsilon_i, \qquad \epsilon_i \sim N(0, \frac{1}{\omega_{ii}}), \\
\gamma_{ij}(X) &= -\frac{\omega_{ij}(\mathbf{X})}{\omega_{ii}} = -\frac{1}{\omega_{ii}} \mathbf{X} \boldsymbol{\beta_{ij}} = -\frac{1}{\omega_{ii}} \left( \sum_{l=1}^{q} \beta_{ijl} X_l \right)
\end{aligned}
\tag{A.1}
$$

where $\odot$ represents the element-wise multiplication between vectors.

Priors on the GraphR model are given below:

$$
\begin{aligned}
&\beta_{ijl} = b_{ijl} s_{ijl}, \\
&b_{ijl} \mid \tau_{il} \sim N(0, \tau_{il}^{-1}), \\
&s_{ijl} \mid \pi_{ijl} \sim \text{Ber}(\pi_{ijl}), \\
&\tau_{il} \sim \text{Gamma}(a_\tau, b_\tau) \\
&\pi_{ijl} \sim \text{Beta}(a_\pi, b_\pi) \\
&\omega_{ii} \propto 1.
\end{aligned}
\tag{A.2}
$$

Parameters of estimation are $\boldsymbol{\theta} = \{\boldsymbol{b}, \boldsymbol{s}, \boldsymbol{\omega}, \boldsymbol{\pi}, \boldsymbol{\tau}\}$.

#### A.2 Mean field variational Bayes

Varitional Bayes method aims to obtain the optimal approximation of true posterior distribution from a class of tractable distributions $Q$, called variational family, by minimizing the Kullback-Leibler (KL) divergence between the approximate $q_{\text{vb}}(\boldsymbol{\theta})$ and the true posterior distribution $p(\boldsymbol{\theta}|\mathbf{Y}, \mathbf{X})$ [2]. A common choice of $Q$, known as mean-field approximation, assumes that $q_{\text{vb}}(\boldsymbol{\theta})$ can be expressed as $\prod_{k=1}^{K} q_{\text{vb}}^{k}(\boldsymbol{\theta_k})$ for some partition of $\boldsymbol{\theta}$. One can write $q_{\text{vb}}(\boldsymbol{\theta})$ as

$$q^*_{\text{vb}}(\boldsymbol{\theta}) \in \arg\min_{q_{\text{vb}}(\boldsymbol{\theta})\in\mathbb{Q}} KL(q_{\text{vb}}(\boldsymbol{\theta})\|p(\boldsymbol{\theta}|\boldsymbol{Y},\boldsymbol{X})), \text{ where}$$

$$KL(q_{\text{vb}}(\boldsymbol{\theta})\|p(\boldsymbol{\theta}|\boldsymbol{Y},\boldsymbol{X})) = \int q_{\text{vb}}(\boldsymbol{\theta})\,\log\left(\frac{q_{\text{vb}}(\boldsymbol{\theta})}{p(\boldsymbol{\theta}|\boldsymbol{Y},\boldsymbol{X})}\right)d\boldsymbol{\theta}$$

$$= -\int q_{\text{vb}}(\boldsymbol{\theta})\,\log\left(\frac{p(\boldsymbol{\theta}|\boldsymbol{Y},\boldsymbol{X})}{q_{\text{vb}}(\boldsymbol{\theta})}\right)d\boldsymbol{\theta} = -\int q_{\text{vb}}(\boldsymbol{\theta})\,\log\left(\frac{p(\boldsymbol{\theta},\boldsymbol{Y},\boldsymbol{X})}{q_{\text{vb}}(\boldsymbol{\theta})}\right)d\boldsymbol{\theta} + \log\left[p(\boldsymbol{Y},\boldsymbol{X})\right]. \tag{A.3}$$

We denote $L[q_{\text{vb}}(\boldsymbol{\theta})] = \int q_{\text{vb}}(\boldsymbol{\theta})\log\left(p(\boldsymbol{\theta},\boldsymbol{Y},\boldsymbol{X})/q_{\text{vb}}(\boldsymbol{\theta})\right)d\boldsymbol{\theta}$ which is the lower bound of the model log-likelihood, and write $KL(q_{\text{vb}}(\boldsymbol{\theta})\|p(\boldsymbol{\theta}|\boldsymbol{Y},\boldsymbol{X})) = -L[q_{\text{vb}}(\boldsymbol{\theta})] + \log\left[p(\boldsymbol{Y},\boldsymbol{X})\right]$. Minimizing KL-divergence is equivalent to maximizing $L[q_{\text{vb}}(\boldsymbol{\theta})]$ since $\log\left[p(\boldsymbol{Y},\boldsymbol{X})\right]$ doesn't involve $\boldsymbol{\theta}$. We can further show that

$$\begin{aligned}
L[q_{\text{vb}}(\boldsymbol{\theta})] &= \int q_{\text{vb}}(\boldsymbol{\theta})\left[\log p(\boldsymbol{\theta},\boldsymbol{Y},\boldsymbol{X}) - \log q_{\text{vb}}(\boldsymbol{\theta})\right]d\boldsymbol{\theta}\\
&= \int q^k_{\text{vb}}(\boldsymbol{\theta_k})\int\left[\left(\log p(\boldsymbol{\theta},\boldsymbol{Y},\boldsymbol{X}) - \log q_{\text{vb}}(\boldsymbol{\theta_k})\right)\right]\prod_{i\neq k} q_{\text{vb}}(\boldsymbol{\theta_i})d\boldsymbol{\theta_{-k}}d\boldsymbol{\theta_k}\\
&\quad -\int\sum_{i\neq k}\log q_{\text{vb}}(\boldsymbol{\theta_i})\prod_{i\neq k} q_{\text{vb}}(\boldsymbol{\theta_i})\int q_{\text{vb}}(\boldsymbol{\theta_k})d\boldsymbol{\theta_{-k}}d\boldsymbol{\theta_k}\\
&= \int q^k_{\text{vb}}(\boldsymbol{\theta_k})\left[\mathbb{E}_{-k}(\log p(\boldsymbol{\theta},\boldsymbol{Y},\boldsymbol{X})) - \log q_{\text{vb}}(\boldsymbol{\theta_k})\right]d\boldsymbol{\theta_k} - \text{const}\\
&= -KL(q^k_{\text{vb}}(\boldsymbol{\theta_k})\|\exp\left[\mathbb{E}_{-k}(\log p(\boldsymbol{\theta},\boldsymbol{Y},\boldsymbol{X}))\right]).
\end{aligned} \tag{A.4}$$

Therefore, we have $q^k_{\text{vb}}(\boldsymbol{\theta_k}) \propto \exp\left[\mathbb{E}_{-k}(\log p(\boldsymbol{\theta},\boldsymbol{Y},\boldsymbol{X}))\right]$.

#### A.3 Update equations

The likelihood is expressed as

$$\begin{aligned}
p(\boldsymbol{\theta},\boldsymbol{Y},\boldsymbol{X}) \propto \prod_{i=1}^{p}&\left\{\left|\frac{1}{\omega_{ii}}I_n\right|^{-\frac{1}{2}}\exp\left[-\frac{\omega_{ii}}{2}\left\|Y_i - \sum_{j\neq i}\gamma_{ij}(X)\odot Y_j\right\|^2\right]\right\}\times\\
&\prod_{i=1}^{p}\prod_{j\neq i}^{p}\prod_{l=1}^{q}\left\{\left(\frac{1}{\tau_{il}}\right)^{-\frac{1}{2}}\exp\left[-\frac{\tau_{il}}{2}(b_{ijl})^2\right](\pi_{ijl})^{s_{ijl}}(1-\pi_{ijl})^{1-s_{ijl}}\right\}\times\\
&\prod_{i=1}^{p}\prod_{l=1}^{q}\left\{(\tau_{il})^{a_\tau-1}\exp\left[-b_\tau\tau_{il}\right]\right\}\times\\
&\prod_{i=1}^{p}\prod_{j\neq i}^{p}\prod_{l=1}^{q}\left\{(\pi_{ijl})^{a_\pi-1}(1-\pi_{ijl})^{b_\pi-1}\right\}
\end{aligned} \tag{A.5}$$

$$\begin{aligned}\log p(\boldsymbol{\theta}, \boldsymbol{Y}, \boldsymbol{X}) = &\text{Const} + \sum_{i=1}^{p} \left\{ \frac{n}{2}\log(\omega_{ii}) - \frac{\omega_{ii}}{2} \left\| Y_i + \frac{1}{\omega_{ii}} \sum_{j \neq i}^{p} \sum_{l=1}^{q} b_{ijl} s_{ijl} X_l \odot Y_j \right\|^2 \right\} \\ &+ \sum_{i=1}^{p} \sum_{l=1}^{q} \left\{ \left( \frac{p-1}{2} + a_\tau - 1 \right) \log(\tau_{il}) - \left( b_\tau + \frac{1}{2} \sum_{j \neq i}^{p} (b_{ijl})^2 \right) \tau_{il} \right\} \\ &+ \sum_{i=1}^{p} \sum_{j \neq i}^{p} \sum_{l=1}^{q} \left\{ (s_{ijl} + a_\pi - 1) \log(\pi_{ijl}) + (b_\pi - s_{ijl}) \log(1 - \pi_{ijl}) \right\}. \quad (A.6)\end{aligned}$$

Due to the dependence between $\boldsymbol{b}$ and $\boldsymbol{s}$[24], the mean-field assumption is considered as:

$$q_{\text{vb}}(\boldsymbol{b}, \boldsymbol{s}, \boldsymbol{\omega}, \boldsymbol{\pi}, \boldsymbol{\tau}) = q_{\text{vb}}(\boldsymbol{b}, \boldsymbol{s}) q_{\text{vb}}(\boldsymbol{\omega}) q_{\text{vb}}(\boldsymbol{\pi}) q_{\text{vb}}(\boldsymbol{\tau}).$$

We can obtain the update equation for each parameter as:

$$q_{\text{vb}}^{k}(\boldsymbol{\theta_k}) \propto \exp \left[ \mathbb{E}_{-k}( \log p(\boldsymbol{\theta}, \boldsymbol{Y}, \boldsymbol{X})) \right].$$

**a. Update of** $\tau_{il}$:

$$\begin{aligned}\log q_{\text{vb}}(\tau_{il}) &= \mathbb{E}_{-\tau_{il}}(l) \\ &= C + \left[ \frac{p-1}{2} + a_\tau - 1 \right] \log \tau_{il} + \left[ b_\tau + \frac{1}{2} \sum_{j \neq i}^{p} \mathbb{E}_{-\tau_{il}} (b_{ijl})^2 \right] \tau_{il}. \quad (A.7)\end{aligned}$$

$$q_{\text{vb}}(\tau_{il}) \sim \Gamma \left( a_\tau + \frac{p-1}{2}, b_\tau + \frac{1}{2} \sum_{j \neq i}^{p} \mathbb{E}_{-\tau_{il}} (b_{ijl})^2 \right)$$

**b. Update of** $\pi_{ijl}$:

$$\begin{aligned}\log q_{\text{vb}}(\pi_{ijl}) &= \mathbb{E}_{-\pi_{ijl}}(l) \\ &= C + \left[ \mathbb{E}_{-\pi_{ijl}}(s_{ijl}) + a_\pi - 1 \right] \log(\pi_{ijl}) + \left[ b_\pi - \mathbb{E}_{-\pi_{ijl}}(s_{ijl}) \right] \log(1 - \pi_{ijl}). \quad (A.8)\end{aligned}$$

$$q_{\text{vb}}(\pi_{ijl}) \sim \text{Beta} \left( \mathbb{E}_{-\pi_{ijl}}(s_{ijl}) + a_\pi, b_\pi - \mathbb{E}_{-\pi_{ijl}}(s_{ijl}) + 1 \right)$$

**c. Update of** $\omega_{ii}$:

$$\begin{aligned}\log q_{\text{vb}}(\omega_{ii}) &= \mathbb{E}_{-\omega_{ii}}(l) \\ &= C + \frac{n}{2}\log(\omega_{ii}) - \frac{\|Y_i\|^2}{2}\omega_{ii} - \frac{\mathbb{E}_{-\omega_{ii}} \| \sum_{j \neq i}^{p} \sum_{l=1}^{q} b_{ijl} s_{ijl} X_l \odot Y_j \|^2}{2} \left( \frac{1}{\omega_{ii}} \right). \quad (A.9)\end{aligned}$$

$$q_{\rm vb}(\omega_{ii}) \sim \text{GIG}\left(\frac{n+2}{2}, \|Y_i\|^2, \mathbb{E}_{-\omega_{ii}}\|\sum_{j\neq i}^{p}\sum_{l=1}^{q} b_{ijl}s_{ijl}X_l \odot Y_j\|^2\right)$$

Here **GIG** represents generalized inverse Gaussian distribution.

**d. Update of** $\beta_{ijl} = b_{ijl}s_{ijl}$:

We Denote $M_{-(m,n)}^{-k} = \sum_{j\neq m}^{p}\sum_{l=1}^{q} b_{mjl}s_{mjl}X_l \odot Y_j - b_{mnk}s_{mnk}X_k \odot Y_n$ and

$$\begin{aligned}\log\, q_{\rm vb}(b_{ijl}|s_{ijl}) &= \mathbb{E}_{-b_{ijl}|s_{ijl}}(l)\\ &= C - \frac{1}{2}\left[\mathbb{E}_{-b_{ijl}|s_{ijl}}(\tau_{il}) + \mathbb{E}_{-b_{ijl}|s_{ijl}}\left(\frac{1}{\omega_{ii}}\right)s_{ijl}\|X_l \odot Y_j\|^2\right](b_{ijl})^2\\ &- \left[Y_i + \mathbb{E}_{-b_{ijl}|s_{ijl}}\left(\frac{1}{\omega_{ii}}\right)\mathbb{E}_{-b_{ijl}|s_{ijl}}M_{-(i,j)}^{-l}\right]^T[X_l \odot Y_j]\, s_{ijl}b_{ijl}. \end{aligned} \tag{A.10}$$

$$q_{\rm vb}(b_{ijl}|s_{ijl}) \sim \mathbb{N}\left(\mu(s_{ijl}), \sigma^2(s_{ijl})\right)$$

$$\sigma^2(s_{ijl}) = \left[\mathbb{E}_{-b_{ijl}|s_{ijl}}(\frac{1}{\omega_{ii}})\|X_l \odot Y_j\|^2 s_{ijl} + \mathbb{E}_{-b_{ijl}|s_{ijl}}(\tau_{il})\right]^{-1}$$

$$\mu(s_{ijl}) = -\sigma^2(s_{ijl})\left\{\left[Y_i + \mathbb{E}_{-b_{ijl}|s_{ijl}}\left(\frac{1}{\omega_{ii}}M_{-(i,j)}^{-l}\right)\right]^T[X_l \odot Y_j]s_{ijl}\right\}$$

The mariginal density of $q_{\rm vb}(s_{ijl})$ is obtained by integrating the joint density of $q_{\rm vb}(b_{ijl}, s_{ijl})$ as

$$\begin{aligned}&q_{\rm vb}(s_{ijl}) = \int \exp\left\{\log\, q_{\rm vb}(b_{ijl}, s_{ijl})\right\} db_{ijl}\\ &= \exp\left\{s_{ijl}\ \mathbb{E}_{-s_{ijl}}\text{logit}(\pi_{ijl})\right\}\int \mathbb{N}_{b_{ijl}}\left(\mu(s_{ijl}), \sigma^2(s_{ijl})\right)\sigma(s_{ijl})\text{exp}\left(\frac{\mu^2(s_{ijl})}{2\sigma^2(s_{ijl})}\right)db_{ijl}\\ &= \sigma(s_{ijl})\text{exp}\left\{s_{ijl}\ \mathbb{E}_{-s_{ijl}}\text{logit}(\pi_{ijl}) + \left(\frac{\mu^2(s_{ijl})}{2\sigma^2(s_{ijl})}\right)\right\}.\\ &\log[q_{\rm vb}(s_{ijl})] = C + \log(\sigma(s_{ijl})) + s_{ijl}\ \mathbb{E}_{-s_{ijl}}\text{logit}(\pi_{ijl}) + \frac{\mu^2(s_{ijl})}{2\sigma^2(s_{ijl})}\\ &\log[q_{\rm vb}(s_{ijl}=0)] = C - \frac{1}{2}\text{log}\mathbb{E}_{-s_{ijl}}\tau_{ijl}\\ &\log[q_{\rm vb}(s_{ijl}=1)] = C + \mathbb{E}_{-s_{ijl}}\text{logit}(\pi_{ijl}) - \frac{1}{2}\text{log}\left[\mathbb{E}_{-s_{ijl}}(\frac{1}{\omega_{ii}})\|X_l \odot Y_j\|^2 + \mathbb{E}_{-s_{ijl}}(\tau_{il})\right]\\ &+ \frac{1}{2}\left[\mathbb{E}_{-s_{ijl}}(\frac{1}{\omega_{ii}})\|X_l \odot Y_j\|^2 + \mathbb{E}_{-s_{ijl}}(\tau_{il})\right]^{-1}\left[(X_l \odot Y_j)^T(Y_i + \mathbb{E}_{-s_{ijl}}\left(\frac{1}{\omega_{ii}}\right)\mathbb{E}_{-s_{ijl}}M_{-(i,j)}^{-l})\right]^2\end{aligned}$$

$$
\begin{aligned}
&s_{ijl} \sim \text{Ber}(\psi_{ijl}) \\
&\log\, q_{\text{vb}}(s_{ijl}) = C + s_{ijl}\text{logit}(\psi_{ijl}) \\
&\psi_{ijl} = \mathbb{E}_{-s_{ijl}}\text{logit}(\pi_{ijl}) - \frac{1}{2}log\left[\mathbb{E}_{-s_{ijl}}(\frac{1}{\omega_{ii}})\|X_l \odot Y_j\|^2 + \mathbb{E}_{-s_{ijl}}(\tau_{il})\right] + \frac{1}{2}log\mathbb{E}_{-s_{ijl}}\tau_{il} \\
&+ \frac{1}{2}\left[\mathbb{E}_{-s_{ijl}}(\frac{1}{\omega_{ii}})\|X_l \odot Y_j\|^2 + \mathbb{E}_{-s_{ijl}}(\tau_{il})\right]^{-1}\left[(X_l \odot Y_j)^T(Y_i + \mathbb{E}_{-s_{ijl}}\left[\frac{1}{\omega_{ii}}\right]\mathbb{E}_{-s_{ijl}}M^{-l}_{-(i,j)})\right]^2
\end{aligned}
$$

#### A.4 Evidence lower bound (ELBO)

As shown previously, the evidence lower bound is defined as:

$$
\begin{aligned}
L[q_{\text{vb}}(\boldsymbol{\theta})] &= \int q_{\text{vb}}(\boldsymbol{\theta})\log\left(p(\boldsymbol{\theta}, \boldsymbol{Y}, \boldsymbol{X})/q_{\text{vb}}(\boldsymbol{\theta})\right) d\boldsymbol{\theta} \\
&= \mathbb{E}_{q_{\text{vb}}(\boldsymbol{\theta})}\ \log(p(\boldsymbol{\theta}, \boldsymbol{Y}, \boldsymbol{X})) - \mathbb{E}_{q_{\text{vb}}(\boldsymbol{\theta})}[q_{\text{vb}}(\boldsymbol{\theta})] \\
&= \mathbb{E}_{q_{\text{vb}}(\boldsymbol{\theta})}\ \log(p(\boldsymbol{\theta}, \boldsymbol{Y}, \boldsymbol{X})) - \sum_{i=1}^{p}\mathbb{E}_{q_{\text{vb}}(\boldsymbol{\theta})}[q_{\text{vb}}(\omega_{ii})] - \sum_{i=1}^{p}\sum_{l=1}^{q}\mathbb{E}_{q_{\text{vb}}(\boldsymbol{\theta})}[q_{\text{vb}}(\tau_{il})] \\
&- \sum_{i=1}^{p}\sum_{j\neq i}^{p}\sum_{l=1}^{q}\left\{\mathbb{E}_{q_{\text{vb}}(\boldsymbol{\theta})}[q_{\text{vb}}(b_{ijl}, s_{ijl})] + \mathbb{E}_{q_{\text{vb}}(\boldsymbol{\theta})}[q_{\text{vb}}(\pi_{ijl})]\right\}.
\end{aligned}
$$

Notably, $-\mathbb{E}_{q_{\text{vb}}(\boldsymbol{\theta})}[q_{\text{vb}}(\omega_{ii})]$, $-\mathbb{E}_{q_{\text{vb}}(\boldsymbol{\theta})}[q_{\text{vb}}(\tau_{il})]$, $-\mathbb{E}_{q_{\text{vb}}(\boldsymbol{\theta})}[q_{\text{vb}}(b_{ijl}, s_{ijl})]$, $-\mathbb{E}_{q_{\text{vb}}(\boldsymbol{\theta})}[q_{\text{vb}}(\pi_{ijl})]$ are entropies of GIG, Gamma, Normal, Bernoulli and Beta distributions, which have a close form. We denote entropy as $H(\cdot)$ and all the expectations below are taken w.r.t $q_{\text{vb}}(\boldsymbol{\theta})$.

**a. Derivation of**$\mathbb{E}\ \log(p(\boldsymbol{\theta}, \boldsymbol{Y}, \boldsymbol{X}))$:

$$
\begin{aligned}
\mathbb{E}\,\log(p(\boldsymbol{\theta},\boldsymbol{Y},\boldsymbol{X})) = \sum_{i=1}^{p}\Bigg\{ & -\frac{n+(p-1)q}{2}\log 2\pi + q\left[a_\tau \log b_\tau - \log\Gamma(a_\tau)\right] \\
& + (p-1)q\left[\log\Gamma(a_\pi+b_\pi) - \log\Gamma(a_\pi b_\pi)\right] \\
& + \frac{n}{2}\mathbb{E}(\log\omega_{ii}) - \frac{\|Y_i\|^2}{2}\mathbb{E}\omega_{ii} - \mathbb{E}(\omega_{ii}^{-1})\mathbb{E}\left\|\sum_{j\neq i}^{p}\sum_{l=1}^{q}\beta_{ijl}Z_l\odot Y_j\right\|^2 \\
& - Y_i^T\left(\sum_{j\neq i}^{p}\sum_{l=1}^{q}\mathbb{E}\beta_{ijl}Z_l\odot Y_j\right) \\
& + \sum_{l=1}^{q}\left[\left(\frac{p-1}{2}+a_\tau-1\right)\mathbb{E}\left(\log\tau_{il}\right) - \left(b_\tau + \frac{\sum_{j\neq i}^{p}\mathbb{E}b_{ijl}^2}{2}\right)\mathbb{E}\tau_{il}\right] \\
& + \sum_{j\neq i}^{p}\sum_{l=1}^{q}\left[\left(\mathbb{E}s_{ijl}+a_\pi-1\right)\mathbb{E}(\log\pi_{ijl}) + (b_\pi - \mathbb{E}s_{ijl})\right]\mathbb{E}\left(\log(1-\pi_{ijl})\right)\Bigg\}
\end{aligned}
$$

**b. Derivation of $H(\tau_{il})$**

$$
\begin{aligned}
H(\tau_{il}) = & -\left[a_\tau + \frac{p-1}{2}\right]\log\left[b_\tau + \frac{1}{2}\sum_{j\neq i}^{p}\mathbb{E}b_{ijl}^2\right] + \log\,\Gamma(a_\tau + \frac{p-1}{2}) \\
& - \left[a_\tau + \frac{p-1}{2} - 1\right]\mathbb{E}(\log\,\tau_{il}) + \left[b_\tau + \frac{1}{2}\sum_{j\neq i}^{p}\mathbb{E}b_{ijl}^2\right]\mathbb{E}\tau_{il}
\end{aligned}
$$

**c. Derivation of $H(\pi_{ijl})$**

$$
\begin{aligned}
H(\pi_{ijl}) = & -\log\,\Gamma(a_\pi+b_\pi+1) + \log\,\Gamma(\mathbb{E}s_{ijl}+a_\pi) + \log\,\Gamma(b_\pi+1-\mathbb{E}s_{ijl}) \\
& - (\mathbb{E}s_{ijl}+a_\pi-1)\mathbb{E}\log\,\pi_{ijl} - (b_\pi - \mathbb{E}s_{ijl})\mathbb{E}\log\,(1-\pi_{ijl})
\end{aligned}
$$

**d. Derivation of $H(\omega_{ii})$**

Denote $a_{\omega_{ii}} = \|Y_i\|^2$ and $b_{\omega_{ii}} = \mathbb{E}\|\sum_{j\neq i}^{p}\sum_{l=1}^{q}\beta_{ijl}Z_l\odot Y_j\|^2$

$$
H(\omega_{ii}) = -\frac{n+2}{4}\log\frac{a_{\omega_{ii}}}{b_{\omega_{ii}}} + \log(2K_{(n+2)/2}\sqrt{a_{\omega_{ii}}b_{\omega_{ii}}}) - \frac{n}{2}\mathbb{E}(\log\omega_{ii}) + \frac{a_{\omega_{ii}}}{2}\mathbb{E}\omega_{ii} + \frac{b_{\omega_{ii}}}{2}\mathbb{E}\omega_{ii}^{-1}
$$

**e. Derivation of $H(b_{ijl}, s_{ijl})$**

$$
H(b_{ijl}, s_{ijl}) = H(b_{ijl}|s_{ijl}) + H(s_{ijl})
$$

where

$$
H(b_{ijl}|s_{ijl}) = \frac{1}{2}\log\left[2\pi\sigma^2(s_{ijl})\right] + \frac{1}{2}
$$

$$H(s_{ijl}) = -\psi_{ijl}\log(\psi_{ijl}) - (1-\psi_{ijl})\log(1-\psi_{ijl})$$

Combining all the previous entropy derivations, we have the following expression of evidence lower bound (ELBO) as

$$\begin{aligned}
L[q_{\text{vb}}(\boldsymbol{\theta})] =& -\frac{np+p(p-1)q}{2}\text{log}2\pi \\
&+ pq\left[a_\tau \text{log} b_\tau - \text{log}\Gamma(a_\tau) + \text{log}\Gamma(a_\tau + \frac{p-1}{2})\right] \\
&+ p(p-1)q\left[\text{log}\Gamma(a_\pi + b_\pi) - \text{log}\Gamma(a_\pi b_\pi) - \text{log}\Gamma(a_\pi + b_\pi + 1)\right] \\
&- \sum_{i=1}^{p}\left\{Y_i^T\left(\sum_{j\neq i}^{p}\sum_{l=1}^{q}\mathbb{E}\beta_{ijl}Z_l \odot Y_j\right) - \frac{n+2}{4}\text{log}\frac{a_{\omega_{ii}}}{b_{\omega_{ii}}} + \text{log}(2K_{(n+2)/2}\sqrt{a_{\omega_{ii}}b_{\omega_{ii}}})\right\} \\
&- \left[a_\tau + \frac{p-1}{2}\right]\sum_{i=1}^{p}\sum_{l=1}^{q}\left\{\text{log}\left[b_\tau + \frac{1}{2}\sum_{j\neq i}^{p}\mathbb{E}b_{ijl}^2\right]\right\} \\
&+ \sum_{i=1}^{p}\sum_{j\neq i}^{p}\sum_{l=1}^{q}\Bigg\{\log\,\Gamma(\mathbb{E}s_{ijl} + a_\pi) + \log\,\Gamma(b_\pi + 1 - \mathbb{E}s_{ijl}) \\
&+ \frac{1}{2}\text{log}\left[2\pi\sigma^2(s_{ijl})\right] + \frac{1}{2} - \psi_{ijl}\log(\psi_{ijl}) - (1-\psi_{ijl})\log(1-\psi_{ijl})\Bigg\}.
\end{aligned}$$

#### A.5 Overview of competing methods

Here we only consider methods which allows for multiple graphical models or covariate-dependent graphs for individual. The GraphR model A.1 estimates covariate-dependent graphs based on Gaussian likelihood with incorporation of discrete and/or continuous variables which encode heterogeneity of samples. Mean-field Variational Bayesian (MFVB) algorithm is implemented in the GraphR in order to achieve computational efficiency. The FGL, GGL[4] and LASICH[19] can only be used in group-specific settings. The Gaussian graphical regression model proposed by Zhang and Li (2022)[27] apply penalized likelihood functions for the inference which allows scenarios for both multiple graphical models and individual covariate-dependent graphs. However those methods fail to provide measurements of uncertainty and probabilistic reasoning. Peterson, Stingo, and Vannucci (2015)[18] propose a Bayesian model for multiple graphical models and implemented a MCMC based method. Wang et al. (2021)[25] and Ni, Stingo, and Baladandayuthapani (2019)[16] also develop methods to estimate conditional dependencies as a function of individual-level or group-level covariates in directed or undirected graphs. However both methods apply MCMC based algorithm and thus fail to scale with high dimensional data set. The Table below summerizes the comparison among these methods.

| | Frequentist or Bayesian | Multi-category graphs | Continuously-varying graphs | Scalability | Uncertainty quantification |
|---|---|---|---|---|---|
| GraphR | Bayesian | ✓ | ✓ | ✓ | ✓ |
| FGL, GGL [4] | Frequentist | ✓ | X | ✓ | X |
| LASICH [19] | Frequentist | ✓ | X | ✓ | X |
| Zhang and Li (2022) [27] | Frequentist | ✓ | ✓ | ✓ | X |
| Peterson, Stingo and Vannucci (2015) [18] | Bayesian | ✓ | X | X | ✓ |
| Ni, Stingo and Baladandayuthapani (2019) [16] | Bayesian | ✓ | ✓ | X | ✓ |
| Wang et al. (2021) [25] | Bayesian | ✓ | ✓ | X | ✓ |

Table A.1: Methodological comparison among different methods.

#### A.6 Algorithm

**Algorithm 1** GraphR algorithm

**Input:** $\mathbf{Y}, \mathbf{X}$, tolerance

**Output:** Covariate dependent edges

**while** $\zeta >$ tolerance **do**

  **for** $i$ in $1:p$ **do**

    **for** $l$ in $1:q$ **do**

      **Set** $q_{\text{vb}}(\tau_{il}) \sim \Gamma\left(a_\tau + \frac{p-1}{2}, b_\tau + \frac{1}{2}\sum_{j\neq i}^{p} \mathbb{E}_{-\tau_{il}}(b_{ijl})^2\right)$

    **end for**

    **for** $j$ in $1:p$ and $j \neq i; l$ in $1:q$ **do**

      **Set** $q_{\text{vb}}(\pi_{ijl}) \sim \text{Beta}\left(\mathbb{E}_{-\pi_{ijl}}(s_{ijl}) + a_\pi, b_\pi - \mathbb{E}_{-\pi_{ijl}}(s_{ijl}) + 1\right)$

    **end for**

    **Set** $q_{\text{vb}}(\omega_{ii}) \sim \text{GIG}\left(\frac{n+2}{2}, \|Y_i\|^2, \mathbb{E}_{-\omega_{ii}}\|\sum_{j\neq i}^{p}\sum_{l=1}^{q} b_{ijl}s_{ijl}X_l \odot Y_j\|^2\right)$

    **for** $j$ in $1:p$ and $j \neq i; l$ in $1:q$ **do**

      $\beta_{ijl} = b_{ijl}s_{ijl}$

      **Set** $q_{\text{vb}}(b_{ijl}|s_{ijl}) \sim \mathbb{N}\left(\mu(s_{ijl}), \sigma^2(s_{ijl})\right)$ where

      $\sigma^2(s_{ijl}) = \left[\mathbb{E}_{-b_{ijl}|s_{ijl}}(\frac{1}{\omega_{ii}})\|X_l \odot Y_j\|^2 s_{ijl} + \mathbb{E}_{-b_{ijl}|s_{ijl}}(\tau_{il})\right]^{-1}$

      $\mu(s_{ijl}) = -\sigma^2(s_{ijl})\left\{\left[Y_i + \mathbb{E}_{-b_{ijl}|s_{ijl}}\left(\frac{1}{\omega_{ii}}M_{-(i,j)}^{-l}\right)\right]^T [X_l \odot Y_j]s_{ijl}\right\}$

      **Set** $q_{vb}(s_{ijl}) \sim \text{Ber}(\psi_{ijl})$ where:

      $\psi_{ijl} = \mathbb{E}_{-s_{ijl}}\text{logit}(\pi_{ijl}) + \frac{1}{2}log\mathbb{E}_{-s_{ijl}}\tau_{il} -$

      $\frac{1}{2}log\left[\mathbb{E}_{-s_{ijl}}(\frac{1}{\omega_{ii}})\|X_l \odot Y_j\|^2 + \mathbb{E}_{-s_{ijl}}(\tau_{il})\right] +$

      $\frac{1}{2}\left[\mathbb{E}_{-s_{ijl}}(\frac{1}{\omega_{ii}})\|X_l \odot Y_j\|^2 + \mathbb{E}_{-s_{ijl}}(\tau_{il})\right]^{-1} \times$

      $\left[(X_l \odot Y_j)^T(Y_i + \mathbb{E}_{-s_{ijl}}\left[\frac{1}{\omega_{ii}}\right]\mathbb{E}_{-s_{ijl}}M_{-(i,j)}^{-l})\right]^2$

    **end for**

  **end for**

  $\zeta$: maximum value of expectation difference for parameters before and after updates.

**end while**

### B Simulation studies

In this Section, we provide more simulation results for undirected (Section B.1) and directed (Section B.2) scenarios with different parameters settings such as: sample size ($n$), number of nodes ($p$), number of intrinsic factors ($q$), type of intrinsic factors, connection probability ($\pi$), regression coefficients of intrinsic factors ($\beta$) and etc. We further discuss the simulation study for illustration of Simpson's paradox in Gaussian graphical models in Section B.3.

#### B.1 Undirected graphical models

**Metrics for comparison:** We use the following measures to compare with other methods (I) true positive rate (TPR); (II) false positive rate (FPR); (III) false discovery rate (FDR); (IV) Matthews correlation coefficient (MCC); (V) the area under the receiver operating characteristic (ROC) curve (AUC). MCC[12] measures the quality of binary classification, ranging from +1 (perfect classification) to -1 (total mismatch).

**Comparative methods:** We compare the GraphR method with the following methods (I) Bayesian Gaussian graphical models (BGGM[14]) (II) graphical lasso (GLASSO[5] (III) fused graphical lasso (FGL[4] (IV) group graphical lasso (GGL[4]) (V) Laplacian shrinkage for inverse covariance matrices from heterogeneous populations (LASICH[19]) (VI) kernel graphical lasso (K-GLASSO[9]).

We run BGGM for 10,000 iterations and discard the first 5,000 as burn-in. The tuning parameters of GLASSO is selected based on a stability approach[10]. For FGL, GGL, k-GLASSO and LASICH, approximated Akaike Information Criterion (AIC) is used for the choice of tuning parameters. We set the hyperparameters for the GraphR method as $a_\tau = 0.005, b_\tau = 0.005, a_\pi = 1, b_\pi = 4$.

**Posterior inference:** We are interested in the parameter selection on both intrinsic factors and edges level. Point estimators of each model parameters are obtained by using expectation with respect to the approximation of posterior distribution, for example the point estimates $\hat{\pi}_{ijl} = \int \pi_{ijl}\ \hat{q}(\pi_{ijl}) d\pi_{ijl} = E_{\hat{q}}(\pi_{ijl})$. With respect to selection of intrinsic factors, we use posterior inclusion probability (PIP) of intrinsic factors, which are defined as $\hat{\boldsymbol{p}}_{\boldsymbol{ij}} := [\hat{p}_{ij1}, ...\hat{p}_{ijq}]^T = [E_{\hat{q}}(s_{ij1}), ...E_{\hat{q}}(s_{ijq})]^T$. For the edge selection between $Y_i$ and $Y_j$, we take both the magnitude of external coefficient $E_{\hat{q}}(\boldsymbol{b}_{ij})$ and PIP of intrinsic factors $E_{\hat{q}}(\boldsymbol{s}_{ij})$ into consideration, and thus define PIP of edges as $\phi_{ij} := \sum_{l=1}^{q} \left[ \frac{\|E_{\hat{q}}(b_{ijl}s_{ijl})X_l\|^2}{\sum_{m=1}^{q}\left(\|E_{\hat{q}}(b_{ijm}s_{ijm})X_m\|^2\right)} \hat{p}_{ijl} \right]$. In the undirected graphical models, we have $\omega_{ij}(\mathbf{X}) = \omega_{ji}(\mathbf{X})$ due to the symmetric property of precision matrix, implying that $s_{ijl}$ and $s_{jil}$ should be 1 or 0 simultaneously. However we parallelize the regressions which leads to significant increase in time-efficacy and also breaks the dependence between $s_{ijl}$ and $s_{jil}$. Thus $s_{ijl} = s_{jil}$ are not necessarily held during the estimation procedure. To ensure the symmetry of selection indicator, namely $\boldsymbol{s_{ij}} = \boldsymbol{s_{ji}}$, we use a post-processing based approach following the steps from[13]. We denote $\kappa_{ex}$ and $\kappa_{edge}$ as the threshold of intrinsic factors and edge detection, where intrinsic factors will be selected or conditional dependence exists between edges if the corresponding PIP is larger than $\kappa_{ex}$ and $\kappa_{edge}$. $\beta_{ijl}$ is non-zero if and only if $\hat{p}_{ijl}^{min} := min(\hat{p}_{ijl}, \hat{p}_{jil}) > \kappa_{ex}$. Similarly, in the edge detection, $\phi_{ij}^{min} := min(\phi_{ij}, \phi_{ji}) > \kappa_{edge} \Leftrightarrow$ edge $\{i, j\}$ exists for $1 \leq i \neq j \leq p$. In terms of the selection of threshold on both external covariate level and edge level, we considered a Bayesian local FDR-based inference approach aiming to control the average Bayesian FDR at some level $\alpha$[3,15]. Denote the threshold on external covariates or on edges at level $\alpha$ as $\kappa_{ex,\alpha}$ and $\kappa_{edge,\alpha}$ respectively, and denote $q_{ijl} = 1 - \hat{p}_{ijl}^{min}$ and $\tilde{q}_{ij} = 1 - \phi_{ij}^{min}$. Notably, $q_{ijl} = q_{jil}$ and $\tilde{q}_{ij} = \tilde{q}_{ji}$ due to the definition of $\hat{p}_{ijl}^{min}$ and $\phi_{ij}^{min}$. PIPs of external covariates and edges represented that posterior probability that the corresponding external covariates or edges were presented from the model, then $q_{ijl}$ and $\tilde{q}_{ij}$ can be regard as estimates of the local FDR for external covariates and edges, which are also known as Bayesian q-values[21]. This statement holds even if we have correlated data[6,3]. Based on q-values of external covariates, $\kappa_{ex,\alpha}$ is determined using follow procedure: (1) sort $\{q_{ijl}\}_{1\leq i<j\leq p, 1\leq l\leq q}$ in an ascending order and notated the ordered set as $\{q^{(t)}\}_{1\leq t\leq p(p-1)q/2}$ (2) Calculate the

cumulative mean of $\{q^{(t)}\}$, and find the maximum $t^*$ such that the cumulative mean of $q^{(t^*)}$ is smaller than $\alpha$, a given significance level. (3) Set $\kappa_{ex,\alpha} = 1 - q^{(t^*)}$. Similar procedure were proposed when choosing $\kappa_{edge,\alpha}$ by replacing $\{q_{ijl}\}_{1\leq i<j\leq p, 1\leq l\leq q}$ with $\{\tilde{q}_{ij}\}_{1\leq i<j\leq p}$.

##### B.1.1 Multi-category Graphs

**Simulation design:** The samples are evenly classified into $q$ groups, and each sample has $q$ discrete intrinsic factors, indicating the group allocations. Graphs for different groups are generated as following:

(I) Graph of the first group $G_1$ is obtained from a Erdos-Renyi graphs with connection probability being $\pi$.

(II) $G_q$ is constructed by randomly excluding three existing edges and including three new edges from $G_{q-1}$ for $q \geq 2$.

(III) The precision matrices and observations in three groups are generated in the same way as step (II) in the homogeneous case.

We **fix** the simulation parameters $n = 151$, $p = 33$ and **vary** (I) $q = 2$ (two group indicators) or $q = 3$ (three group indicators); (II) $\pi = 2\%$ or $5\%$.

**Selection performance:** Figure B.1 shows the selection performance of GraphR and comparison methods with respect to multi-category graphs (groups 2 and 3) with different sparsity levels (2% and 5%). Mean values of AUC, MCC, TPR, FPR and FDR based on 50 repetitions are reported. The GraphR method performs the best in terms of MCC while producing very similar performance for TPR like other methods under all the simulation settings mentioned in the simulation design. The proposed method takes a hit on TPR to produce the lowest FDR and FPR than any other methods for all the simulation settings.

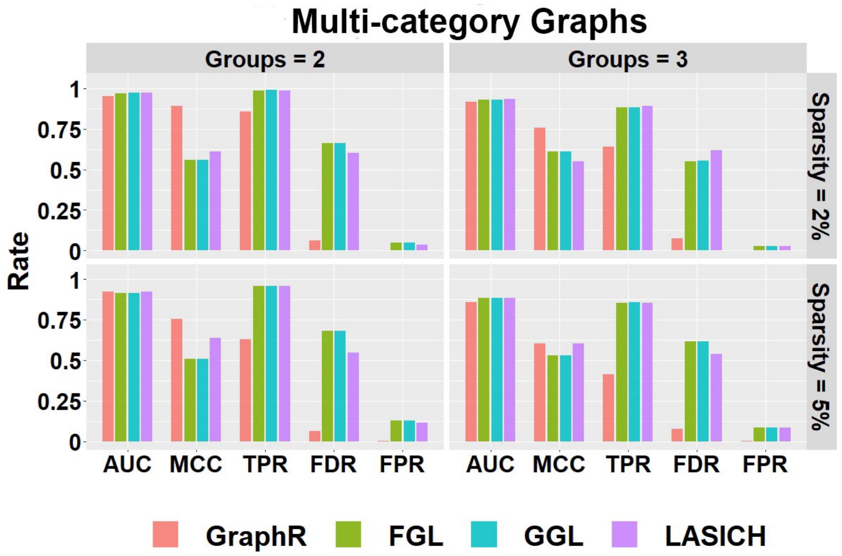


Figure B.1: Selection Performance in multi-category graphs setting with varying number of groups and sparsity.

**Computation time:** Figure B.2 shows the mean of computation time in seconds for GraphR and other competing methods w.r.t. 50 replications along with different groups and sparsity levels. GraphR is more computationally efficient than LASICH for all the settings. The LASSO based methods (FGL and GGL) are marginally faster than GraphR but those methods are unable to quantify uncertainty.

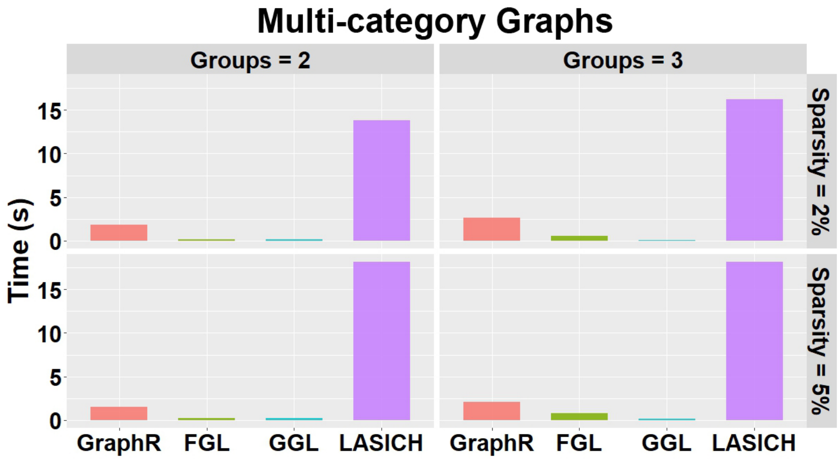


Figure B.2: Computation time(s) in group-specific setting with varying number of groups and varying sparsity.

**Summary:** For the multi-category graph settings, GraphR has marginally similar performance with three comparison methods in terms of AUC. However, in case of other overall performance indicator MCC, GraphR outperforms all other methods. Though GraphR is inferior to comparison methods in TRP, FDR and FPR for GraphR is much lower, which can be viewed as a trade-off. GraphR is also computationally efficient with $< 2.5$s computation time on average for all cases.

##### B.1.2 Continuously Varying Graphs

**Simulation design:** Data are generated based on the assumed model by introducing two continuous intrinsic factors. (I) For $i < j$, 2% of coefficients for continuous intrinsic factors $\beta_{ijl}$ are uniformly chose from -1 or 1 while others are set to be 0. Let the corresponding $\beta_{jil}$ equal to $\beta_{ijl}$. Two continuous intrinsic factors are drawn from Uniform $(-1, 1)$. We construct an individual-specific precision matrix $\Omega_n$ by fixing the diagonal entries $\omega_{ii}$ to be 1 and calculating $\omega_{nij} = \sum_{l=1}^{2} \beta_{nijl} X_{nl}$. (II) Repeat step (I) until $\Omega_n$ is positive definite for all $n$. (III) Generate $y_n$ from $N(0, \Omega_n^{-1})$.
We set the parameters as $n = 151$, $p = 33$, $q = 2$, $\pi = 2\%$.

**Computation time:** Figure B.3 shows the mean of computation time in seconds of GraphR and comparison methods based on 50 repetitions.

**Summary:** Consistent with previous findings, GraphR is computationally efficient than

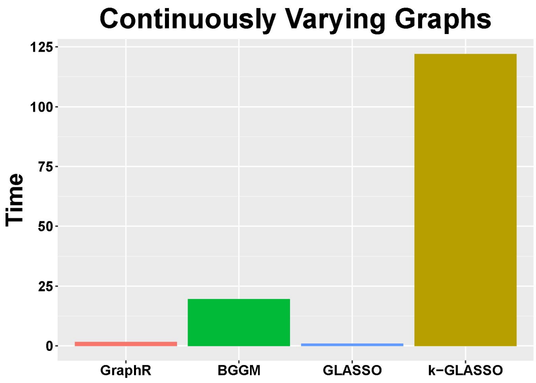


Figure B.3: Computation time(s) in continuously-varying case.

BGGM and k-GLASSO, though takes more computation time than GLASSO. However GLASSO cannot accommodate for heterogenous settings and provide probabilistic reasoning.

##### B.1.3 Homogeneous Graphs

**Simulation design:** We assume that all individuals are homogeneous, implying $\Omega_n = \Omega$, $\forall\ n \in \{1, ..., 151\}$. We only include one constant effect as the intrinsic factor, and generate data as following:

(I) Generate a Erdős–Rényi graph G by setting connection probability 2%. If the edge $\{i, j\}$ are connected in G, then the corresponding off-diagonal entries $\omega_{ij} = \omega_{ji}$ are uniformly drawn from $[-1, -0.5] \cup [0.5, 1]$ otherwise $\omega_{ij}$ are set to be 0. The diagonal entries $\omega_{ii}$ are set to be 1.

(II) The simulated $\Omega$ from (I) is not necessarily to be positively definite, and thus we will add $0.1I_{33}$ to $\Omega$ until it is positively definite.

(III) Generate 151 independent observations from $N(0, \Omega^{-1})$ and set intrinsic factors to be intercept only.

We fix the simulation parameters as $n = 151$, $p = 33$, $q = 1$ (intercept only model), $\pi =$ 2% or 5%.

**Selection performance:** Figure B.4 shows the selection performance of GraphR and comparison methods by using mean values of AUC, MCC, TPR, FPR and FDR based on 50 repetitions with sparsity being 5%. GraphR performs better than GLASSO except for TPR. BGGM performs marginally better than GraphR in the homogeneous setting. GraphR produces marginally better or at par with reduced level of TPR.

**Computation time:** Figure B.5 shows the mean of computation time in seconds for GraphR and other competitive methods with different levels pf sparsity. The results are based on 50 replications. GraphR is significantly faster than BGGM since GraphR is VB based algorithm whereas BGGM is based on Markov Chain Monte Carlo (MCMC). The frequentist method GLASSO is computationally efficient than GraphR but GLASSO is unable to do the uncertainty quantification.

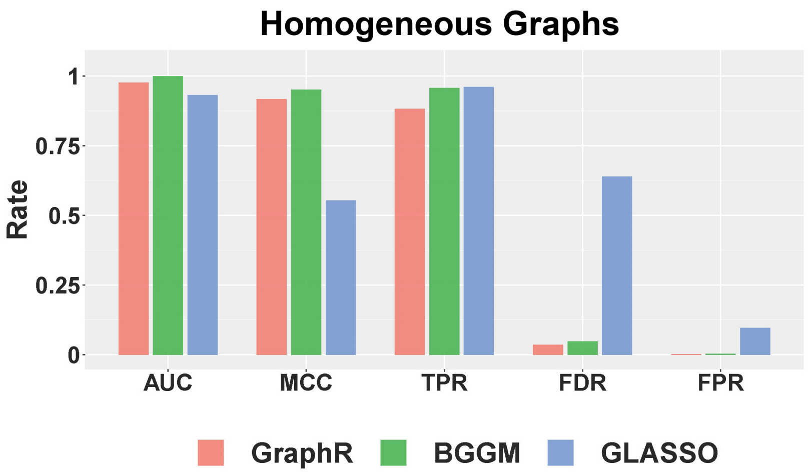


Figure B.4: Selection Performance in homogenuous setting for 5% sparsity level.

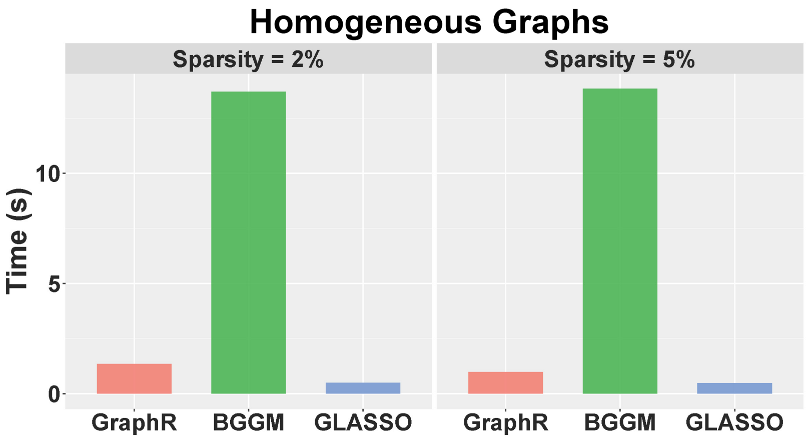


Figure B.5: Computation times for homogenuous setting with sparsity level at 2% and 5%.

**Summary:** By design GraphR method is favorable for heterogeneous settings. For the homogeneous setting, GraphR outperforms the frequentist method GLASSO and marginally similar performance with the Bayesian method BGGM at the cost of reduced FDR level. GraphR is computationally efficient than Bayesian methods. The method produces better efficacy rates while enabling uncertainty quantification unlike the frequentist methods.

#### B.2 Directed acyclic graphs (DAGs)

Here we consider a special case of our GraphR where directions of edges are pre-assumed, leading to a directed acyclic graph model. Three simulation settings are discussed here to illustrate the performance the GraphR method in case of variable selection and scalability for moderate and/or large number of nodes (p) and intrinsic factors (q).

**Simulation design:** Data is generated as following:

(1) Based on the proposed types of intrinsic factors, we generate continuous intrinsic factors from Uniform(0,1) and discrete intrinsic factors from Bernoulli(0.5). In the settings with large number of intrinsic factors, 70% of intrinsic factors are set to be continuous.

(2) For each external variables, 2% second layer of regression coefficients $\beta_{ij}^{(k)}$ are randomly selected to be non-zero and are set to be 3 when $i > j$.

(3) The first node ($i = 1$) is generated from $N(0, 1)$. In terms of $i^{th}$ node $Y_i$ ($i \geq 2$), we standardize $Y_j$ for all $j < i$, and denote these standardized nodes as $\tilde{Y}_j$. Draw $Y_i$ from Normal distribution with mean being $\sum_{j<i} \left[ (\beta_{ij}^{(1)} Z^{(1)} + \beta_{ij}^{(2)} Z^{(2)}) \odot \tilde{Y}_j \right]$ and standard deviation being 1.

We use the same 5 metrics (TPR, FPR, FDR, MCC, AUC) to compare across multiple settings and set the hyperparameters $a_\tau = 0.005, b_\tau = 0.005, a_\pi = 1, b_\pi = 1$. The probability cutoff is selected to control the Bayesian FDR at 1%.

##### B.2.1 Moderate dimensions

**Simulation parameters:** We **fix** the simulation parameters $p = 50$, $q = 2$, $\pi = 2\%$ and **vary** the ratio $n/pq = 1, 2, 3, 4, 5$ and type of intrinsic factors (discrete/continuous).

**Selection performance:** Figure B.6 shows the selection performance of intrinsic factors for the GraphR method in DAG where number of nodes and intrinsic factors are moderate. Each column of the panel indicates types of intrinsic factors while each row represent magnitude of $\beta$. Mean values and 95% confidence interval (CI) of AUC, MCC, TPR, FPR and FDR based on 50 repetitions are reported.

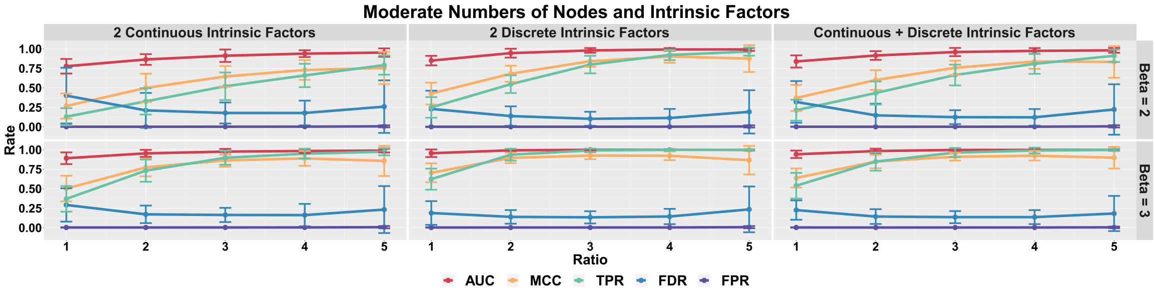


Figure B.6: Selection performance in terms of intrinsic factors in directed acyclic graphs with moderate setting.

Figure B.7 shows the edge selection performance of GraphR in the DAG setting where number of nodes and intrinsic factors are moderate. All other settings are same as ref(fig:modext).

The selection performance of GraphR in terms of intrinsic factors and edges is generally better with increasing effect size ($\beta$) and ratio between $n$ and $pq$. However when $n/pq$ ratio is 5, FDR has is slightly larger than the case when $n/pq$ ratio is 3 or 4, leading to decrease in MCC. One of the potential reason is that estimates are local optimum. Moreover, GraphR tended to perform slightly better with more discrete continuous intrinsic factors when $n/pq$ is not large enough.

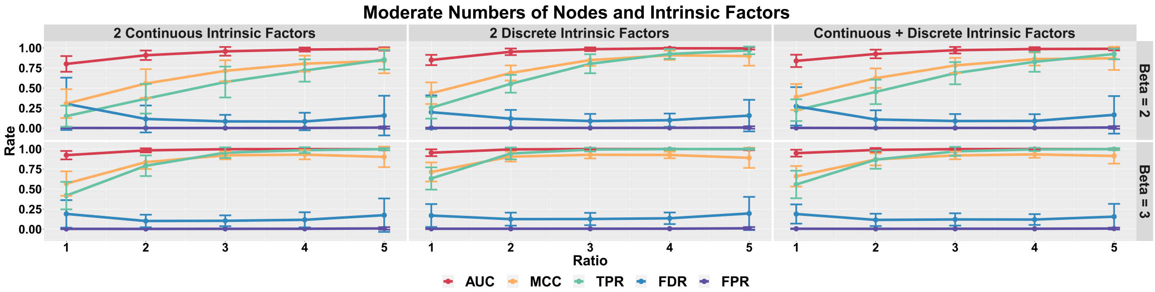


Figure B.7: Selection performance in terms of edges in directed acyclic graphs with moderate setting.

**Computation times:** Figure B.8 shows the mean and 95% CI of computation time in seconds for GraphR with different types of intrinsic factors and $n/pq$ ratio. Results are based on 50 repetitions. We observe an increasing pattern till the ratio is 4 and a sudden drop at 5, partly due to the fact that convergence to normality is easier to fulfilled with larger sample size.

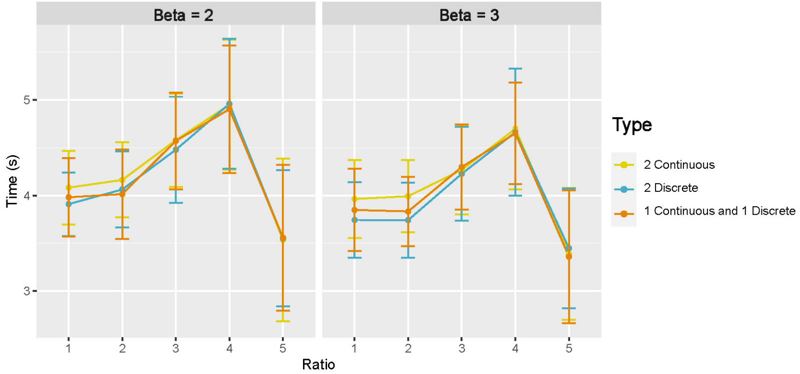


Figure B.8: Computation time in directed acyclic graphs with moderate setting.

##### B.2.2 Large number of nodes

**Simulation parameters** We **fix** the simulation parameters $q = 2$, $\pi = 2\%$ and **vary** the ratio $n/pq = 1, 2, 3, 5$ (i.e. $p = 100, 200, 300, 400, 500$), type of intrinsic factors (discrete/continuous) and magnitude of regression coefficients ($\beta = 2, 3$).

**Selection performance:** Figure B.9 and B.10 show the selection performance of GraphR in terms of intrinsic factors and edges in directed acyclic graphs. Number of nodes are set to be 100, 200, 300 and 500 as shown on the top of each plot. Other settings are same as B.6 and B.7.

Generally, we can observe an increasing trend in AUC, MCC and TPR and decreasing trend in FDR and FPR with larger $n/pq$ ratio and effect size which is similar as the pattern in the moderate setting. However, there are some exception. For example when number

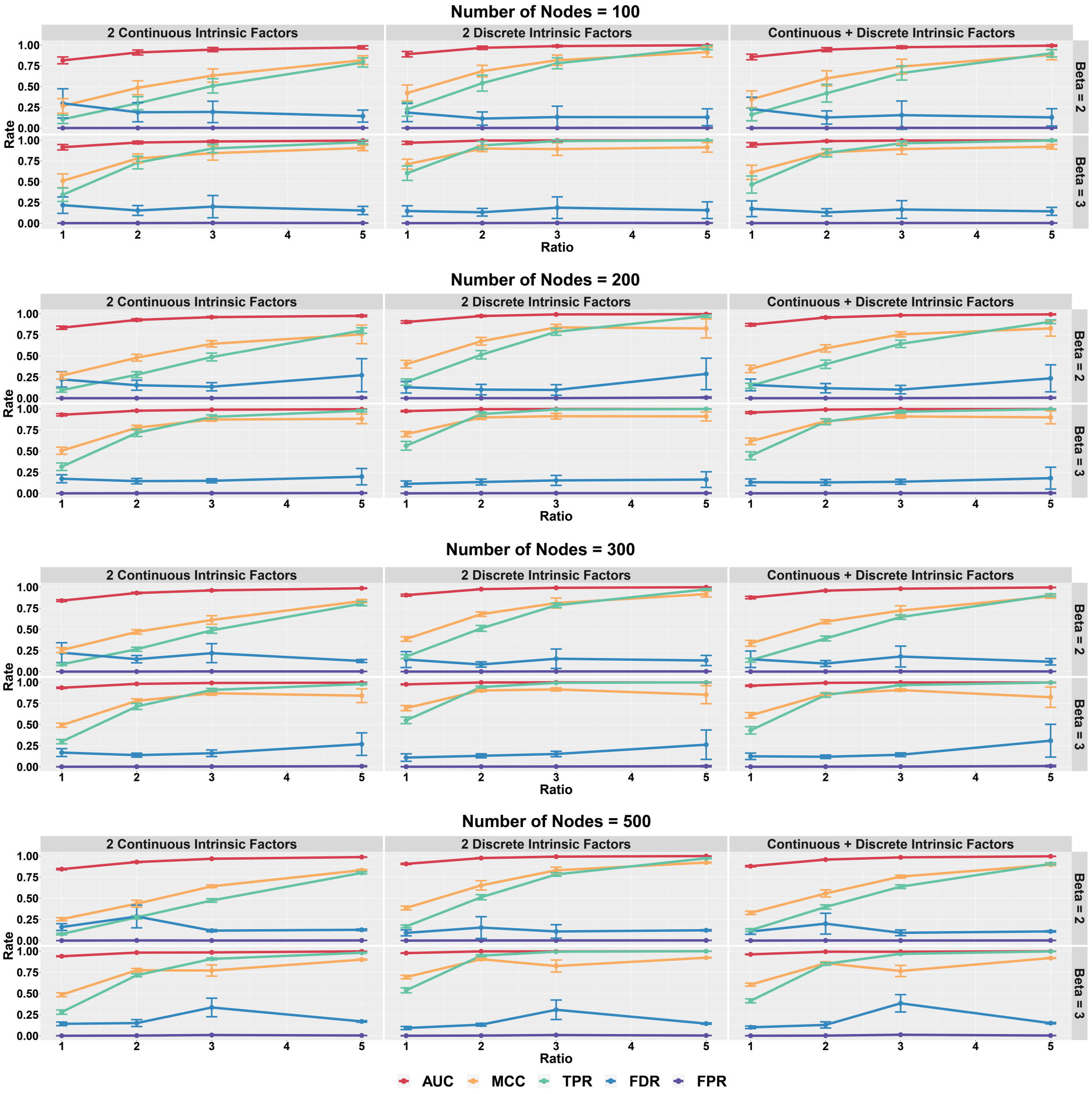


Figure B.9: Selection performance in terms of intrinsic factors in directed acyclic graphs with large number of nodes.

of nodes is 500, there is a sudden jump of FDR at $n/pq = 3$ with relatively large standard deviation, which also can be reflected the decrease in MCC. This phenomena might be caused by some cases where only local optimum are found.

**Computation times:** Figure B.11 shows the mean and 95% CI of computation time in seconds of GraphR based on 50 repetitions. Each row of panel represent types of intrinsic factors while each column indicates the magnitude of coefficients. Number of nodes are set to be 100, 200, 300 and 500, represented by different colors. In most of the cases, longer computation times are needed when sample size and number of nodes increase.

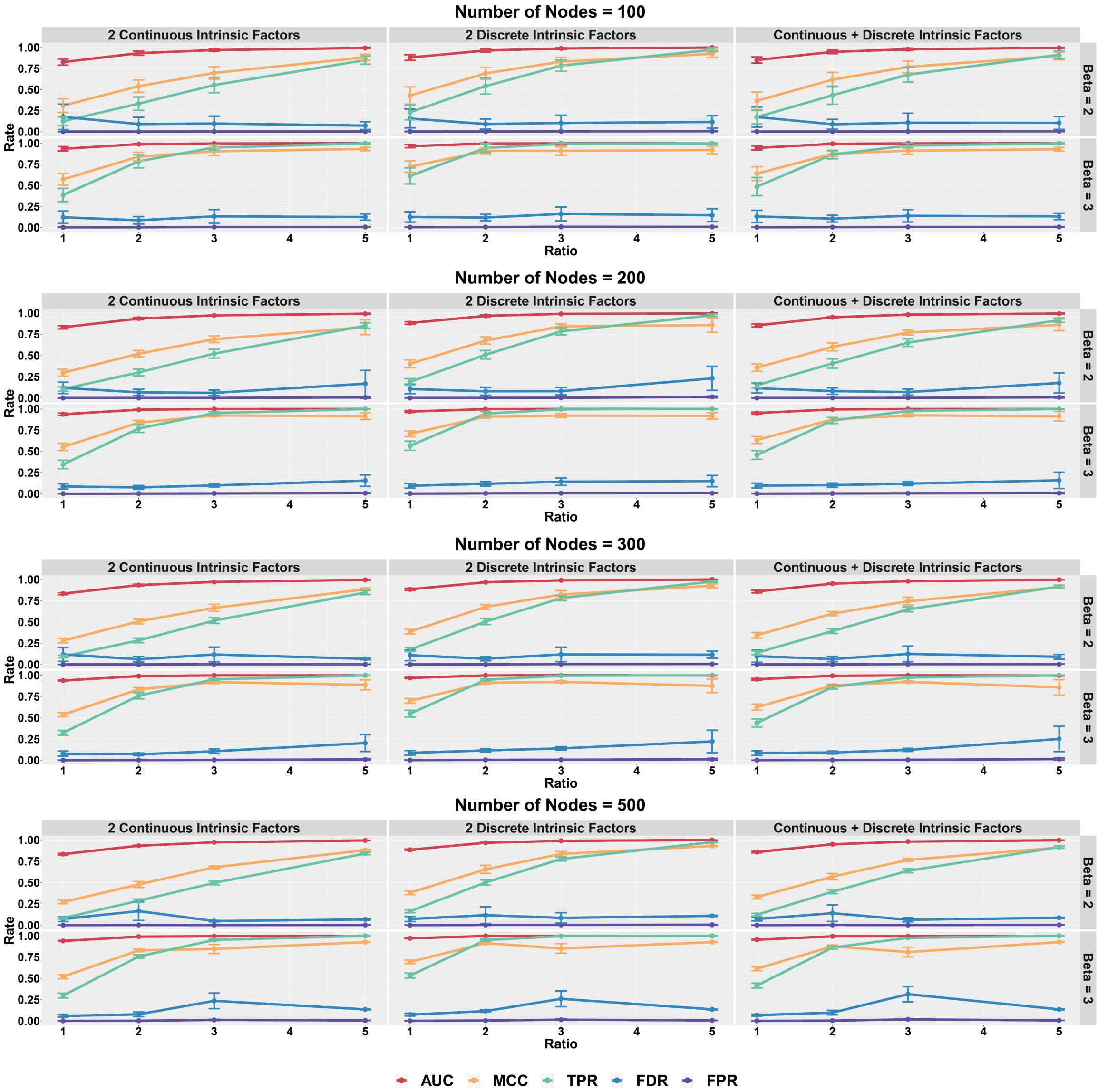


Figure B.10: Selection performance in terms of edges in directed acyclic graphs with large number of nodes.

##### B.2.3 Large number of intrinsic factors

**Simulation parameters:** We **fix** the simulation parameters $p = 50$, $\pi = 2\%$, and 70% of intrinsic factors are set to be continuous. We also **vary** the ratio $q = 2, 5, 10$, $n/pq = 1, 2, 3, 5$ and magnitude of regression coefficients ($\beta = 2, 3$).

**Selection performance:** Figure B.12 and B.13 show the selection performance of GraphR in terms of intrinsic factors and nodes in directed acyclic graphs. Number of intrinsic factors are set to be 2, 5 and 10 and 70% of covariates are continuous. Other settings are same as B.6 and B.7.

Similar as previous findings, GraphR performs better with larger $n/pq$ ratio and effect

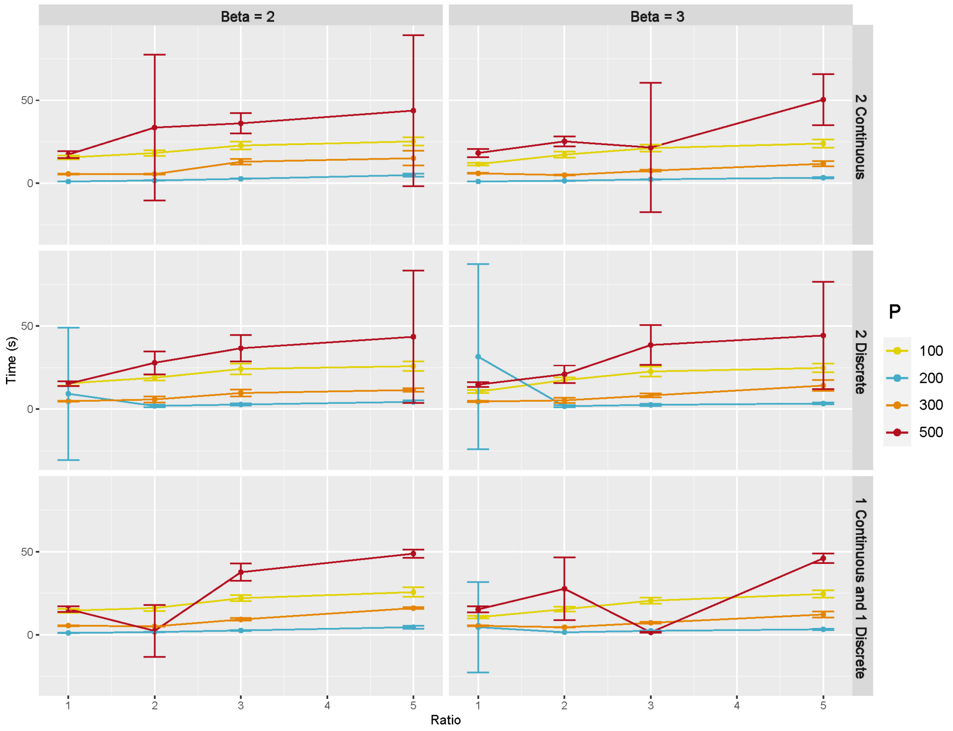


Figure B.11: Computation time in directed acyclic graphs with large number of nodes.

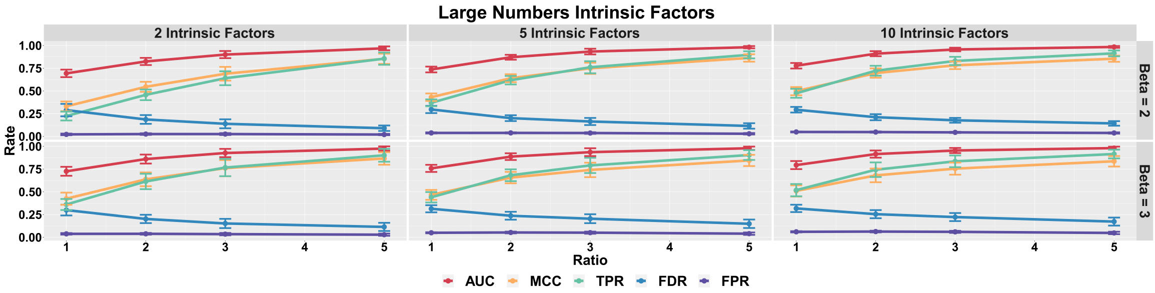


Figure B.12: Selection performance in terms of intrinsic factors in directed acyclic graphs with large number of intrinsic factors.

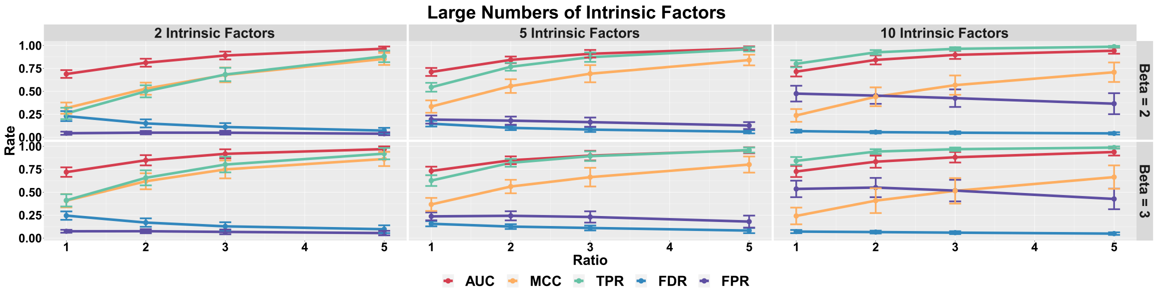


Figure B.13: Selection performance in terms of edges in directed acyclic graphs with large number of intrinsic factors.

size. Importantly, the accuracy of selection also decreases when q is relatively large, suggesting that higher $n/pq$ is needed in order to ensure the quality of estimation.

**Computation time:** Figure B.14 shows the mean and 95% CI of computation time in seconds for GraphR based on 50 repetitions. Number of intrinsic factors are set to be 2, 5 and 10, represented by different colors.

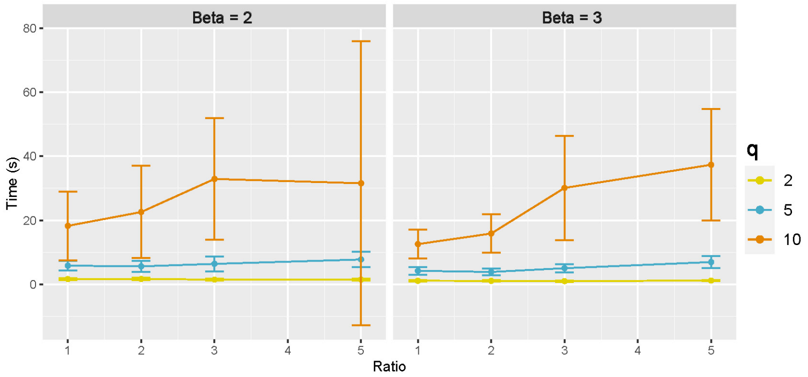


Figure B.14: Computation time in directed acyclic graphs with large number of intrinsic factors.

Computation time increase with larger number of intrinsic factors regardless of the effect sizes. Moreover, variability are also greater when number of intrinsic factors are larger.

#### B.3 Illustration of Simpson's paradox

**Simulation design:** We set overall sample size as 200, number of nodes as 3 and number of groups as 2. Samples are evenly drawn from two groups with distinct conditional independence structures. Generation of precision matrices $\Omega, \tilde{\Omega} \in \mathbb{R}^{3\times 3}$ for the first and the second group is described as follow:

(I) Denote $\Omega = [\omega_{ij}]_{3\times 3}$. Three diagonal entries are set as 1. For off-diagonal entries, $\omega_{12} = \omega_{21}$ are uniformly drawn from $[0.5, 1]$, $\omega_{13} = \omega_{31}$ are uniformly drawn from $[-1, -0.5]$, and $\omega_{23} = \omega_{32}$ are set to be 0.

(II) Denote $\tilde{\Omega} = [\tilde{\omega}_{ij}]_{3\times 3}$. Three diagonal entries are set as 1. For off-diagonal entries, $\tilde{\omega}_{12} = \tilde{\omega}_{21}$ are uniformly drawn from $[-1, -0.5]$, $\tilde{\omega}_{23} = \tilde{\omega}_{32}$ are uniformly drawn from $[0.5, 1]$, and $\tilde{\omega}_{13} = \tilde{\omega}_{31}$ are set to be 0.

Repeat (I) and (II) until both $\Omega, \tilde{\Omega}$ are positive definite matrices.

(III) Generate 100 observations from $N(0, \Omega^{-1})$ and 100 observations from $N(0, \tilde{\Omega}^{-1})$, and set intrinsic factors to be intercept only for homogeneous case and to be group indicators for heterogeneous case.

### C Proteomic Network-based Characterization of Intrinsic Subtypes of Breast Cancer

#### C.1 Data description

The PAM50[17] proteomics dataset was obtained from The Cancer Genome Atlas (TCGA)[26] and The Cancer Proteome Atlas (TCPA)[8]. Reverse Phase Protein Arrays (RPPAs) were used to quantify 190 proteins or phosphoproteins which were involved in apoptosis, DNA damage response, cell cycle[1] over 859 breast cancer (BRCA) patients where normal-like BRCA patients are excluded due to small sample sizes. BRCA of Luminal A and Luminal B are combined in one category to achieve higher power which leads to three groups with 626 BRCA patients in Luminal A and B, 75 patients in Her2-enriched and 158 in Basal-like.

#### C.2 Preprocessing and application

Proteomics data is standardized before applying GraphR. The indicators for cancer subtype (Luminal A+B, Her2-enriched and Basal-like breast cancers) are used as intrinsic factors.

We set hyperparameters as follows: $a_\tau = b_\tau = 0.005, a_\pi = 1, b_\pi = 4$. Notably, in case of integrated functional pathway analysis, we change the hyper-parameters: $a_\pi = b_\pi = 0.05$. This provides us potential to consider high density of connections between proteins in the same pathway.

Moreover, we here only include proteins pairs belonging to different isoforms while showing significantly strong correlation (FDR based p-values $< 0.01$ and magnitude $| \rho | \geq 0.4$) for at least one group, allowing plots to be more readable.

#### C.3 Results

Figure C.1 shows heatmap of posterior inclusion probability (PIP) of selected edges in each PAM50 subtype of BRCA.

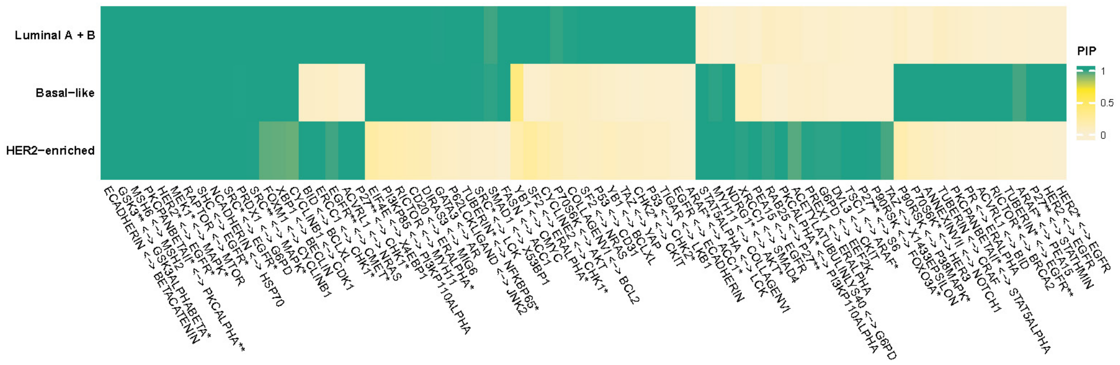


Figure C.1: Posterior inclusion probability (PIP) of selected edges in each PAM50 subtype of BRCA.

Figure C.2, C.3 and C.4 show networks of selected protein pairs in Luminal A+B, Her2-enriched and Basal-like BRCA respectively. The widths of edges are proportional to partial correlations. The sizes of nodes are proportional to connectivity degrees which are defined as the sum of magnitudes of the partial correlations. Sign of partial correlations are represented by color with red being positive and blue being negative.

**Luminal A + B breast cancer**

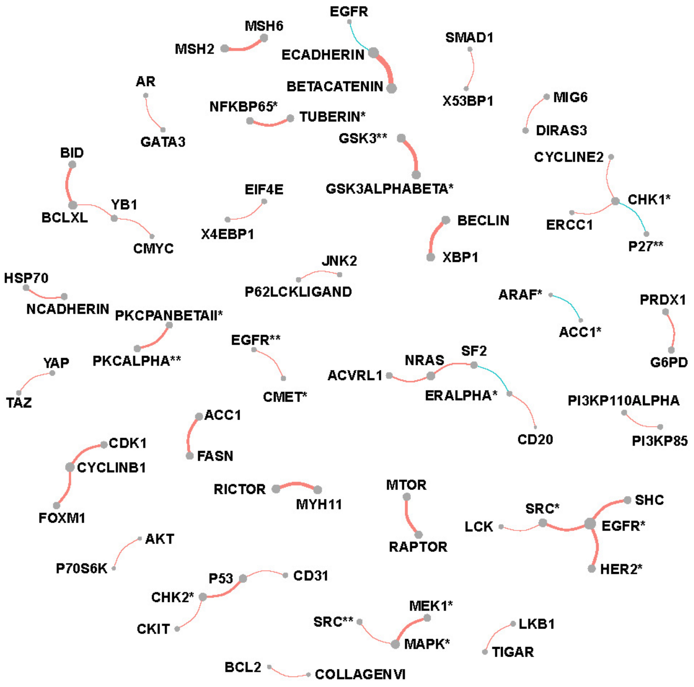


Figure C.2: Network of selected protein pairs in Luminal A + B breast cancer patients.

#### Her2-enriched breast cancer

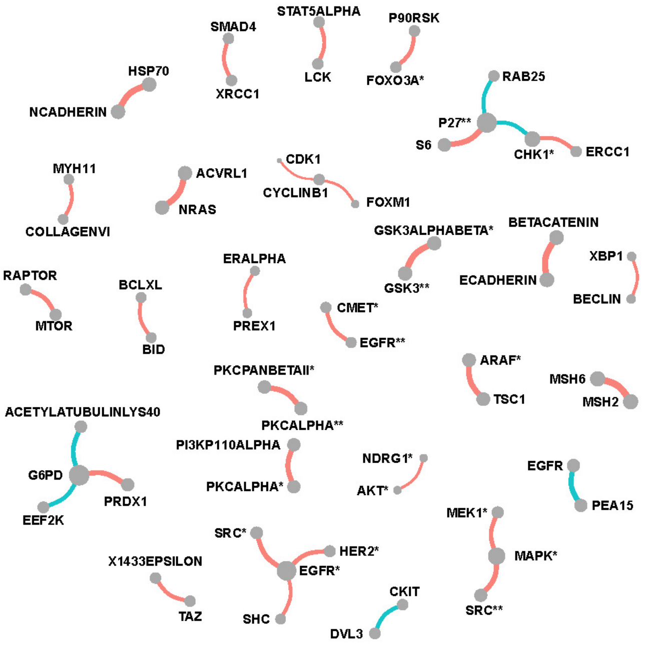


Figure C.3: Network of selected protein pairs in Her2-enriched breast cancer patients.

#### Basal-like breast cancer

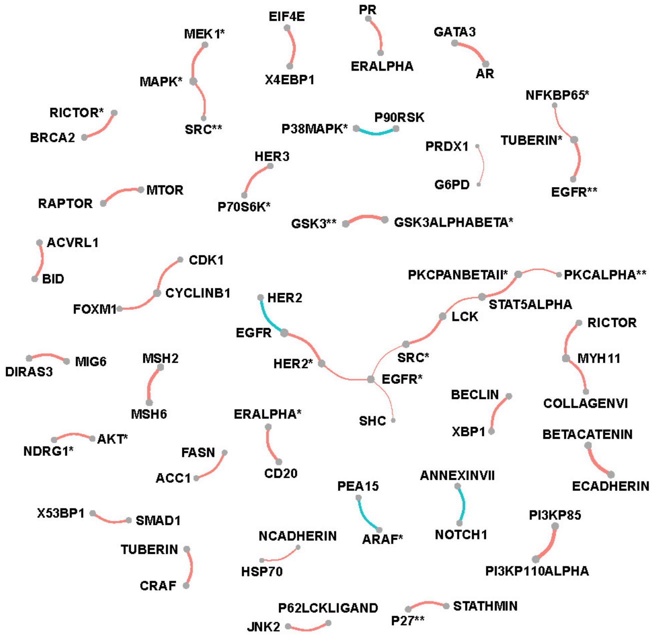


Figure C.4: Network of selected protein pairs in Basal-like breast cancer patients.

### D Characterization of Stemness-induced Heterogeneity on Proteomics Networks in Breast Cancer

#### D.1 Data description

The dataset comes from TCGA[26] and TCPA[8], and is processed by[11] to derive two independent stemness indices based on DNA methylation (mDNAsi) and mRNA expression (mRNAsi). Aim to provide the degrees of dedifferentiation on epigenetic and gene expression level, mDNAsi and mRNAsi range from 0 to 1 with lower values implying tendency to normal-like cells. 189 protein abundance are measured across 616 breast cancer (BRCA) from TCGA and can be downloaded from the NIH Genomic Data Commons (GDC) at https://gdc.cancer.gov/about-data/publications/PanCanStemness-2018.

#### D.2 Preprocessing and application

The mRNAsi, mDNAsi and patients' ages are treated as three intrinsic factors. Logit transformation is used to the stemness indices to ensure the same scale as age, and all three intrinsic factors and proteomics data are standardized before plugging into the model. Hyperparameters are set as previous section C.2.

Here we present proteomics data analysis with mDNAsi in breast cancers where mRNAsi and patients' age are set to be median, and mRNAsi related result can be found in the Result Section. For mDNAsi case, we limit the protein connections by using the following criteria

(1) proteins pair with different isoforms,
(2) significant correlation (FDR based p-values $< 0.01$) in more than half of the cases,
(3) the magnitude of correlations exceeded 0.2 for at least one case.

#### D.3 Results

Figure D.1 and D.2 are two heatmaps which show posterior inclusion probability (PIP) and partial correlation of selected edges with varying mDNAsi while mRNAsi and age are set to be median. The color bar on the left shows quarterly divided mDNAsi. The barplot on top represents the number of significant cases for the corresponding protein pair.

Figure D.3 presents networks for proteins with top five connectivity degrees corresponded to each quarter of mDNAsi. The width of edges are proportional to the median value of partial correlation between selected protein pairs for each quarter of mDNAsi. The node sizes reflects median values of connectivity degrees.

As shown in Figure D.4 we also present changes of correlation along with mDNAsi of the original scale for selected protein pairs when mRNAsi and age are fixed to be median.

Finally, the results based on integrated functional analysis of pathways are shown in D.5. This plot illustrates the changing patterns of connectivity scores of pathways along with

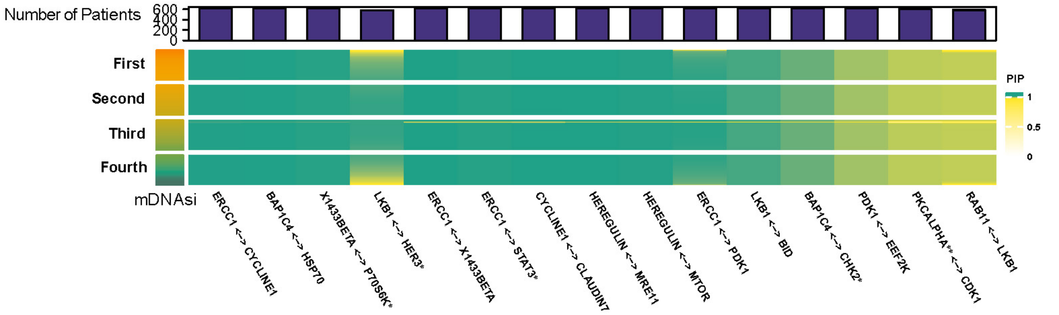


Figure D.1: Posterior inclusion probability (PIP) of selected edges with varying mDNAsi and median value of mRNAsi and age.

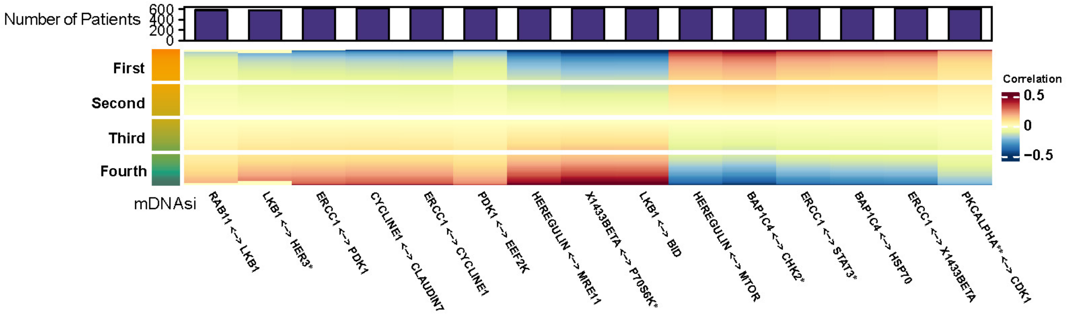


Figure D.2: Partial correlation of selected edges with varying mDNAsi and median value of mRNAsi and age.

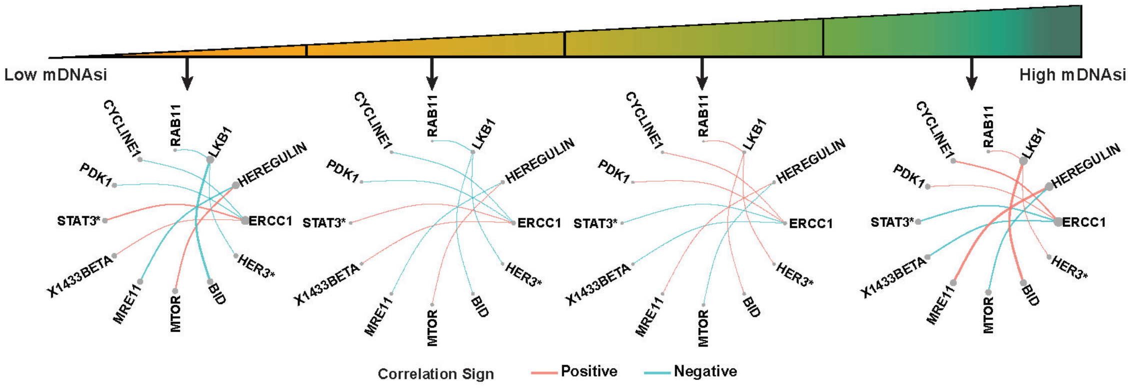


Figure D.3: Networks for proteins with top five connectivity degrees corresponded to each quarter of mRNAsi.

mDNAsi of the origin scale.

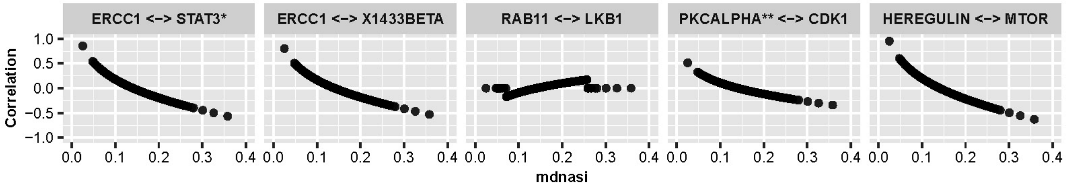


Figure D.4: Line plots of selected protein pairs showing changes of correlation along with mDNAsi of the original scale.

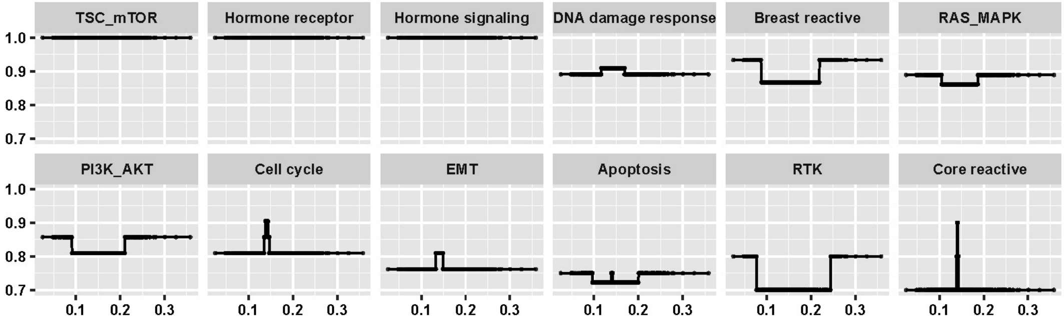


Figure D.5: Changing patterns of connectivity scores for pathways along with mDNAsi of the origin scale.

### E Pan-Cancer Proteomic Network-based Characterization of Breast and Gynecologic Cancers

#### E.1 Data description

The Gynecological and breast cancer (Pan-gyane) proteomics dataset was obtained from TCGA[26] and TCPA[8]. The Pan-gyane proteomics contains four types of cancers including breast invasive carcinoma (BRCA), cervical squamous cell carcinoma and endocervical adenocarcinoma (CESC), ovarian serous cystadenocarcinoma (OV) and uterine corpus endometrial carcinoma (UCEC). Abundance of 189 kinds of protein were measured across 1,941 patients among which 892 patients had BRCA, 173 had CESC, 436 had OV and 440 had UCEC.

#### E.2 Preprocessing and application

All settings are same as in Section C.2, except that:

(I) Four intrinsic factors: indicators of BRCA, CESC, OV, UCEC are included in the analyses.

(II) Significantly strong correlation is defined as connection with FDR based p-values $< 0.01$ and magnitude $| \rho |\geq 0.4)$

#### E.3 Results

Figure E.1 is a heatmap which shows posterior inclusion probability (PIP) of selected edges in each type of cancer.

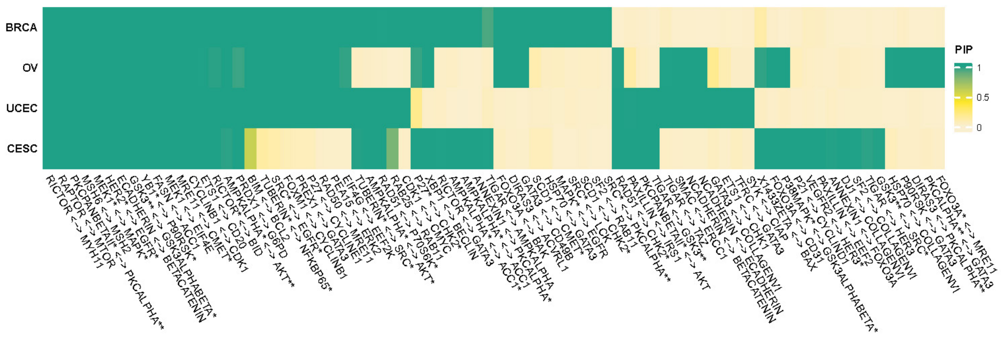


Figure E.1: Posterior inclusion probability (PIP) of selected edges in each type of cancer.

Figure E.2, E.3, E.4 and E.5 show networks of selected protein pairs in BRCA, CESC, OV and UCEC respectively. All other settings are same as figure C.2

**BRCA**

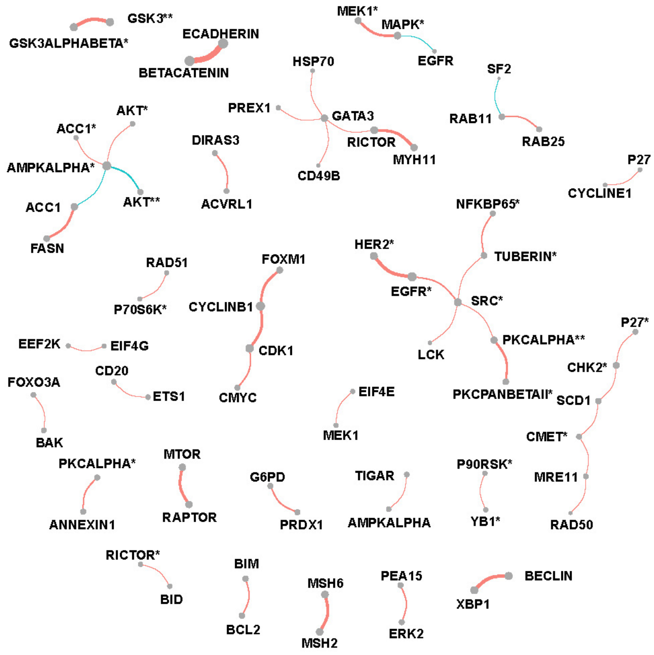


Figure E.2: Network of selected protein pairs in BRCA patients.

#### CSEC

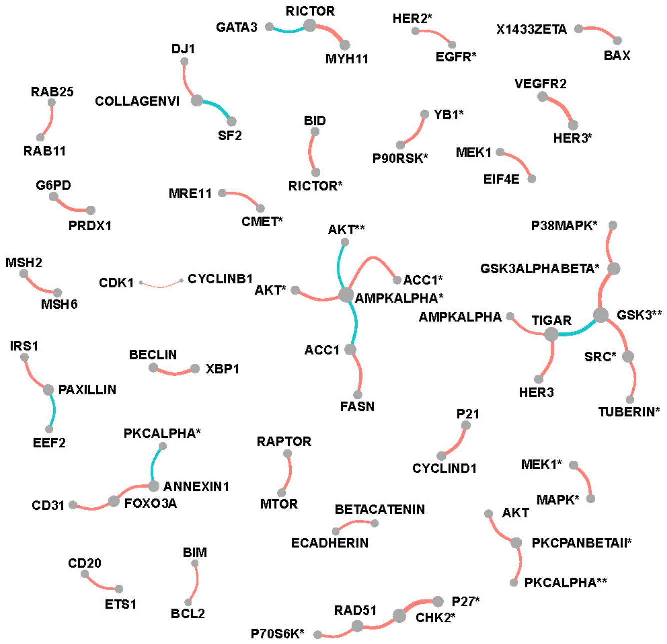


Figure E.3: Network of selected protein pairs in CESC patients.

## OV

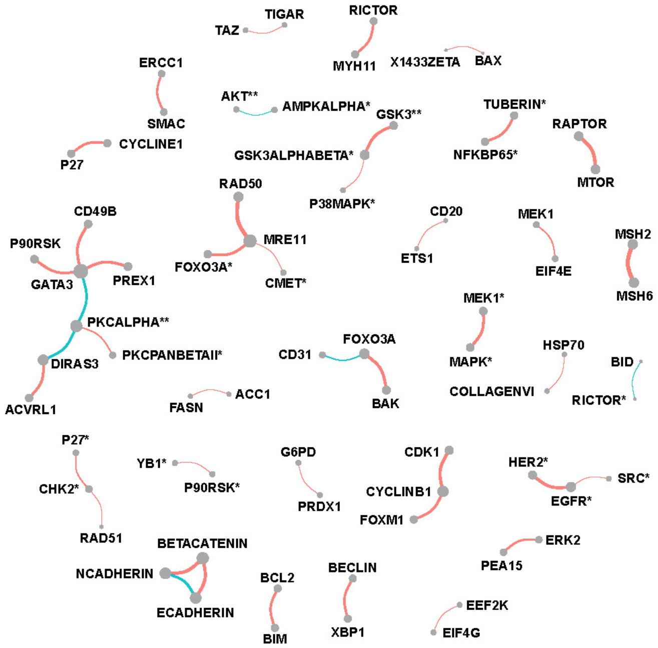


Figure E.4: Network of selected protein pairs in OV patients.

#### UCEC

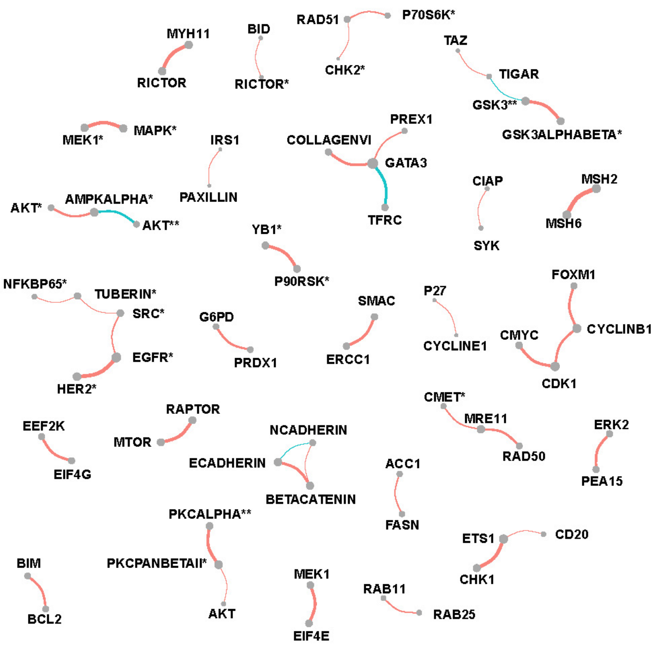


Figure E.5: Network of selected protein pairs in UCEC patients.

### F Gene Expression Network Estimation incorporating Spatial Heterogeneity

#### F.1 Data description

The human breast cancer data was collected from biopsy of breast cancer at a thickness of 16 $\mu$m[20]. Based on the Hematoxylin and Eosin (H&E) staining image, locations can be classified into three spatial regions as tumor, intermediate, and normal with the sizes 114, 67, and 69 spots respectively. The data includes measurement of 5262 genes expression at 250 spot locations.

#### F.2 Preprocessing and application

Here we only consider the 100 spatially expressed genes with the lowest Benjamini-Hochberg (BH) adjusted p-value by applying SPARK method[23]. Next, we apply the PQLseq[22] algorithm to adjust for the covariate effect and obtain the latent gene expressions which follow Normal distribution.

Two coordinates are scaled and treated as intrinsic factors in the GraphR model. Hyperparameters are set as previous section C.2. We include correlations with FDR based p-values $< 0.01$ in the results.

#### F.3 Results

Suppose the partial correlations between gene $i$ and gene $j$ and the corresponding posterior inclusion probabilities are vectors of length $n_1+n_2+n_3$ with $n_1, n_2, n_3$ representing number of spots in tumor, intermediate and normal region, namely

$$\begin{aligned} \rho_{ij} &= [\rho_{ij}^{tumor}, \rho_{ij}^{inter}, \rho_{ij}^{normal}] \in \mathbb{R}^{n_1+n_2+n_3}, \\ PIP_{ij} &= [PIP_{ij}^{tumor}, PIP_{ij}^{inter}, PIP_{ij}^{normal}] \in \mathbb{R}^{n_1+n_2+n_3}. \end{aligned} \quad (1)$$

We define weighted average of partial correlations between gene $i$ and gene $j$ in a region as $\hat{\rho}_{ij}^{region} = (\sum_{region} \rho_{ij}^{region} * PIP_{ij}^{region})/n_{region}$ where region can be tumor, intermediate or normal. Weighted connectivity degree of gene $i$ is defined as the sum of $|\hat{\rho}_{i\cdot}|$.

Figure F.1, F.2 and F.3 show networks w.r.t. each spatial regions. The edges are proportional to the weighted average of partial correlations and nodes are proportional to the weighted connectivity degrees.

**Tumor region**

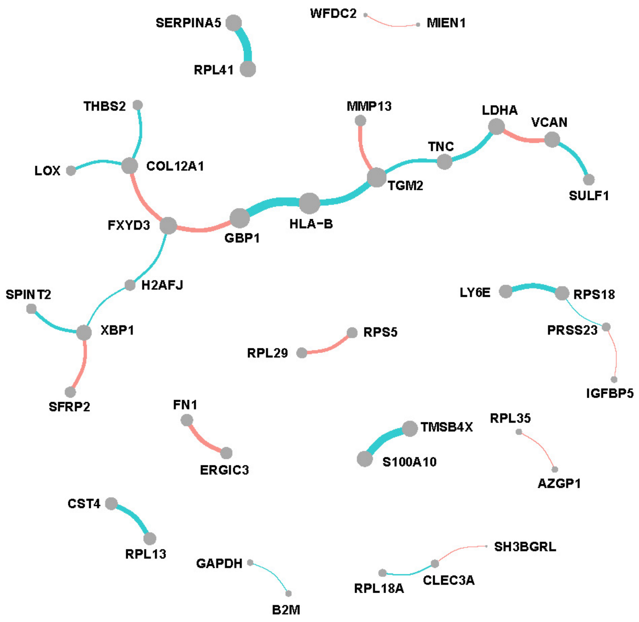


Figure F.1: Network of tumor region in breast cancer.

#### Intermediate region

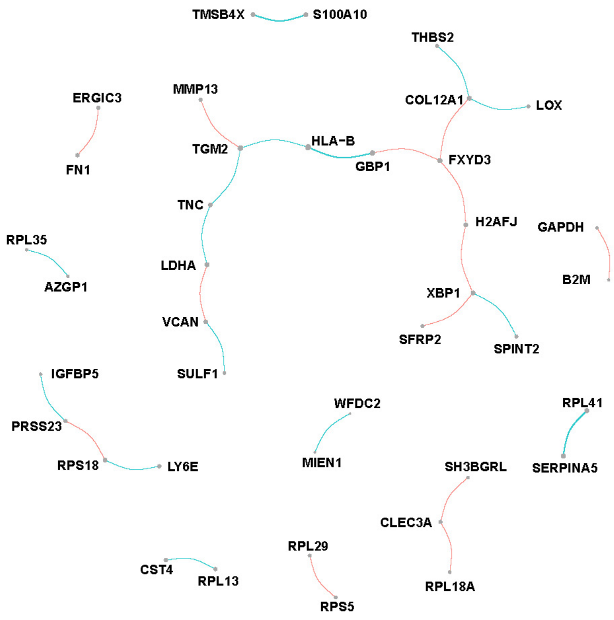


Figure F.2: Network of intermediate region in breast cancer.

#### Normal region

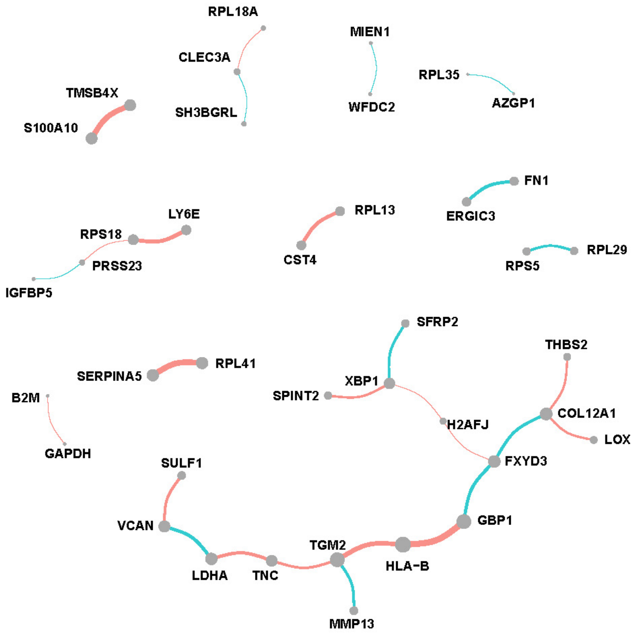


Figure F.3: Network of normal region in breast cancer.

We also display more spatial patterns of partial correlations (Figure F.4) and connectivity degrees (Figure F.5) for gene and gene pairs. The color bar indicates the values of correlations and connectivity degrees while shapes of point representing the spatial region.

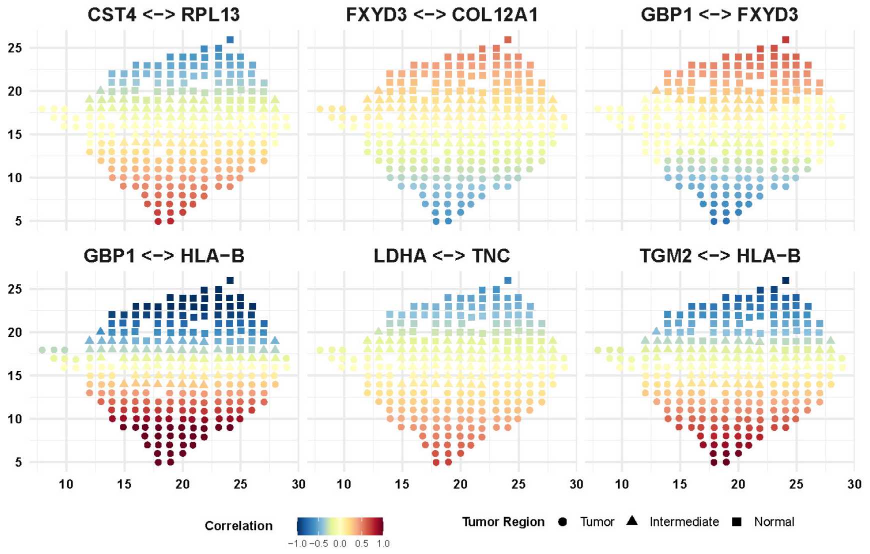


Figure F.4: Spatial pattern of partial correlations for selective gene pairs.

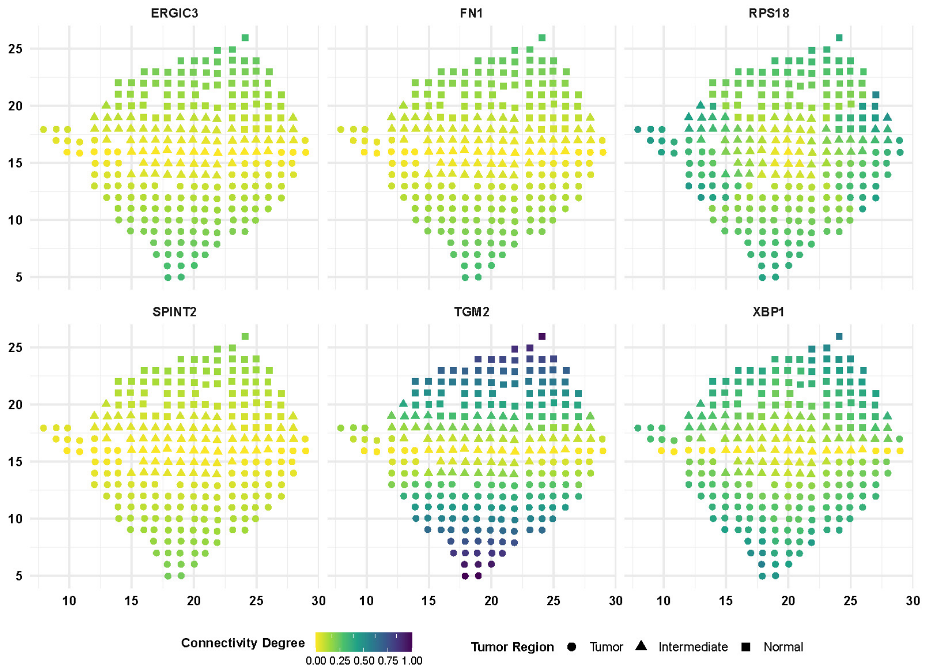


Figure F.5: Spatial pattern of connectivity degrees of selective genes.

### G. Implementation

The GraphR (Graphical Regression) is a flexible approach which incorporates sample heterogenity and enables covariate-dependent graphs. Our regression-based method provides a functional mapping from the covariate space to precision matrix for different types of heterogeneous graphical model settings. GraphR imposes sparsity in both edge and covariate selection and computationally efficient via use of variational Bayes algorithms. The method is versatile to incorporate different type of covariates such as (I) **binary** (control and disease specific graphs), (II) **categorical** (category specific graphs such as cancer subtypes), (III) **univariate continuous** (time varying graphs for single cell data), (IV) **categorical + univariate continuous** (graphs changing over category such as cancer sub-types and continuous scale as biomarkers), (V) **multivariate continuous** (spatial transcriptomics co-expression networks). More details about the method can found in the Methods Section of the manuscript and Section A of the Supplementary Materials. GraphR is implemented as an open-source R package (Section G.1) and Shiny app (Section G.3).

#### G.1 GraphR package

##### G.1.1 Installation

You can install the released version of GraphR from (https://github.com/bayesrx/GraphR) with:

```
devtools::install_github("bayesrx/GraphR")
library(GraphR)
```

##### G.1.2 GraphR_est() function

The **GraphR_est()** function can be used to estimate the graphical regression coefficients and inclusion probabilities of external covariates for the GraphR models. It is suggested to maintain $n/pq > 1$ and efficacy of the method increase with high values of $n/pq$ ratio. For priors, we assume $\pi \sim Beta(a_\pi, b_\pi)$ and $\tau \sim \Gamma(a_\tau, b_\tau)$.

The **mandatory inputs** of estimation function are given below.

- **Features (nodes)**: Nodes of the graphs among which edges are built (e.g. a gene expression matrix of dimensions $n \times p$). **Please standardize features before plugging into the function or set standardize_feature = TRUE in the function**.
- **Cont_external and dis_external (continuous and discrete external covariates)**: An $n \times q_1$ and an $n \times q_2$ matrices of continuous and discrete intrinsic factors respectively. $q_1 + q_2 = q$. **Please standardize continuous intrinsic factors before plug into the estimation function or set standardize_external = TRUE in the function.**

The **optional inputs** of estimation function are given below.

- $\boldsymbol{a_\pi}$, $\boldsymbol{b_\pi}$: Hyper-parameters from $\pi \sim Beta(a_\pi, b_\pi)$. By default $a_\pi = 1, b_\pi = 4$.
- $\boldsymbol{a_\tau}$, $\boldsymbol{b_\tau}$: Hyper-parameters from $\tau \sim Gamma(a_\tau, b_\tau)$. By default $a_\tau = 0.005, b_\tau = 0.005$.
- **Standardize_feature, standardize_external**: Standardize features or continuous intrinsic factors. Default as FALSE.
- **Max_iter**: Maximum number of iterations. Default as 2,000.
- **Max_tol**: Maximum tolerance. Default as 0.001.

**Outputs** of the **GraphR_est()** function are provided below.

- **Beta (the graphical regression coefficients)**: A $p \times p \times q$ array of coefficients for intrinsic factors. The $[i, j, k]$ element represents the effect of k-th external covariates on regression of j-th node on i-th node.
- **Phi (posterior inclusion probability)**: A $p \times p \times q$ array storing posterior inclusion probability (PIP) of external covariates. The $[i, j, k]$ elements represents the PIP of k-th intrinsic factors on regression of j-th node on i-th node.
- **Omega_diag (diagonal elements of precision matrix)**: A p vector with i-th element representing the inverse variance of error.

##### G.1.3 GraphR_pred() function

The **GraphR_pred()** function can be used to predict partial correlation between two nodes and the corresponding inclusion probabilities from the results of GraphR model alongwith Bayesian FDR-adjusted p-values.

The **mandatory inputs** of prediction function are given below.

- **New_df**: A matrix of new intrinsic factors based on which predictions are made. **Note: Please ensure that the order and scale of new intrinsic factors are same as those used in the estimation.**

The **optional inputs** of prediction function are given below.

- **GraphR_est_res**: Results from `GraphR_est` function. If graphR_est_res = NULL, then the following three inputs: (1) beta;

  (2) phi; (3) omega_diag are needed.
- **Beta**: A $p \times p \times q$ array storing coefficients of intrinsic factors. The $[i, j, k]$ elements represents the effect of k-th intrinsic factors on regression of j-th node on i-th node.
- **Omega_diag**: A p vector with i-th element representing the inverse variance of error.
- **Pip**: A $p \times p \times q$ array storing posterior inclusion probability (PIP) of intrinsic factors. The $[i, j, k]$ elements represents the PIP of k-th intrinsic factors on regression of j-th node on i-th node.

The **output** contains following information.

- **Feature_id1**, **feature_id2**: Indices of features or nodes.
- **Pr_inclusion**: Posterior inclusion probability of connections between two nodes based on "And" rules.
- **Correlation**: Partial correlation between two nodes. Values with maximum magnitudes are provided.
- **FDR_p**: Bayesian FDR-adjusted p values.

##### G.1.4 GraphR_visualization() function

The **GraphR_visualization()** function provides a circular network based on a given new intrinsic factors vector and thresholds for FDR-p values and magnitudes of partial correlations.

The **mandatory inputs** of prediction function are given below.

- **New_vec**: A vector of new intrinsic factors based on which plot is made. **Note: Please ensure that the order and scale of new intrinsic factors are same as those used in the estimation.**

The **optional inputs** of prediction function are given below.

- **GraphR_est_res**: Results from `GraphR_est` function. If graphR_est_res = NULL, then the following three inputs: (1) beta;

  (2) phi; (3) omega_diag are needed.

- **Beta**: A $p \times p \times q$ array storing coefficients of intrinsic factors. The $[i, j, k]$ elements represents the effect of k-th intrinsic factors on regression of j-th node on i-th node.
- **Omega_diag**: A p vector with i-th element representing the inverse variance of error.
- **Pip**: A $p \times p \times q$ array storing posterior inclusion probability (PIP) of intrinsic factors. The $[i, j, k]$ elements represents the PIP of k-th intrinsic factors on regression of j-th node on i-th node.
- **Fdr_thre**: A numeric value. Threshold for Bayesian FDR adjusted q-values.
- **Magnitude_thre**: A numeric value. Threshold for the magnitude of partial correlations.

The **output** provides a circular network plot. Node sizes represent connectivity degrees of the corresponding features while edge widths are proportional to the partial correlation between two features. Sign of the partial correlations are represented by the color

#### G.2 Example

An example code with one of the existing datasets to demonstrate how to run the functions and obtain inference.

##### G.2.1 Example

Here we provide an example to run the GraphR method with application to PAM50 protiomics data.

```
set.seed(100)
library(GraphR)
data("Pam50")

features <- as.matrix(apply(Pam50$features,2,scale))
features[c(1:5),c(1:5)]
#>      X1433EPSILON     X4EBP1 X4EBP1_pS65 X4EBP1_pT37T46    X53BP1
#> [1,]   -0.9298711 -1.0325344  -0.1814837      0.3870419 -1.125110
#> [2,]   -1.2265417 -0.8121828  -0.9249897     -0.4834352  1.084052
#> [3,]   -0.9250730 -0.1882466   0.8258123     -0.4022269  0.289943
#> [4,]   -0.6566337 -0.2473042   0.2114522      0.9897723  2.134105
```

```r
#> [5,]   -0.9476849  1.7654120   2.7128204      2.1739453  1.378139

external <- as.matrix(Pam50$external)
external[c(1:5),]
#>      basal_like her2_enriched luminal_ab
#> [1,]          0             1          0
#> [2,]          0             0          1
#> [3,]          0             0          1
#> [4,]          0             0          1
#> [5,]          0             0          1


system.time(res <- GraphR_est(
  features,
  external,
  a_pi = 1,
  b_pi = 4,
  a_tau = 0.005,
  b_tau = 0.005,
  max_iter = 2000,
  max_tol = 0.001
))
#>    user  system elapsed
#>  179.90   62.22  245.62

####### prediction
new_df <- diag(3)
colnames(new_df) <- colnames(external)

system.time(pred <- GraphR_pred(new_df, res))
#>    user  system elapsed
#>    6.10    0.04    6.24
head(pred)
#>   basal_like her2_enriched luminal_ab     feature1       feature2 Pr_inclusion
#> 1          1             0          0     PKCALPHA PKCALPHA_pS657            1
#> 2          1             0          0      ERALPHA             PR            1
#> 3          1             0          0 S6_pS235S236   S6_pS240S244            1
#> 4          1             0          0          BID       STATHMIN            1
#> 5          1             0          0          YAP       YAP_pS127            1
#> 6          1             0          0        RAD51     X4EBP1_pT70            1
#>   Correlation FDR_p
#> 1   0.7938597     0
#> 2   0.4530799     0
#> 3   0.8136693     0
#> 4   0.4157183     0
#> 5   0.7698744     0
#> 6   0.3773436     0

####### visualization
new_vec <- c(1,0,0)
GraphR_visualization(new_vec, graphR_est_res = res,
                     fdr_thre = 0.01, magnitude_thre = 0.4)
#> Joining with `by = join_by(feature)`
```

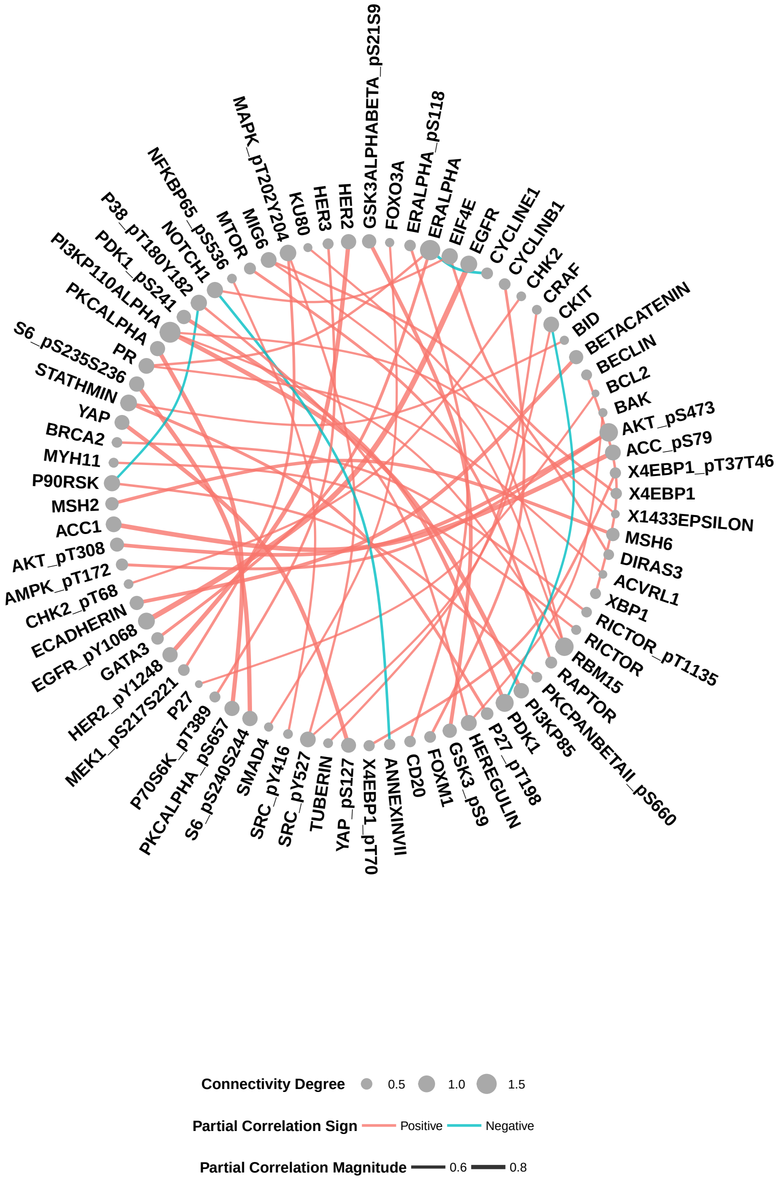


#### G.3 GraphR Shiny App and tutorial website

The Shiny App and tutorial website of GraphR can be found here.

#### G.4 Checkmarks

Here are a few checkmarks to follow while using the GraphR method.

- **We recommend to standardize continuous intrinsic factors while plugging into the functions.**
- **It is suggested to maintain** $n/pq > 1$ and efficacy of the method increase with high values of $n/pq$ ratio.
